# The Global Canopy Atlas: analysis-ready maps of 3D structure for the world’s woody ecosystems

**DOI:** 10.1101/2025.08.31.673375

**Authors:** Fabian Jörg Fischer, Becky Morgan, Toby Jackson, Jérôme Chave, David Coomes, KC Cushman, Ricardo Dalagnol, Michele Dalponte, Laura Duncanson, Sassan Saatchi, Rupert Seidl, Krzysztof Stereńczak, Gaia Vaglio Laurin, Stephen Adu-Bredu, Jesús Aguirre-Gutiérrez, Benedetta Antonielli, John David Armston, Mauro LR de Assis, Nicolas Barbier, Andrew Burt, Ricardo Gomes César, Jaroslav Cervenka, Nicholas Coops, Laury Cullen, James W Dalling, Andrew Davies, Miro Demol, Jakob Ebenbeck, Fabian Fassnacht, Lola Fatoyinbo, Mariano García, N. Ignacio Gasparri, Terje Gobakken, Tristan R.H. Goodbody, Eric Bastos Görgens, Tolga Gorum, Carl Gosper, Hongcan Guan, Janne Heiskanen, Marco Heurich, Martina Hobi, Bernhard Höfle, Al Hooijer, Andreas Huth, Alexander Kedrov, James R Kellner, Simon Koenig, Kamil Král, Misha Krassovski, Helga Kuechly, Martin Krůček, Kyaw Kyaw Htoo, Nicolas Labrière, Daphne Lai, Johannes Larson, Hjalmar Laudon, Dawn Lemke, Jonathan Lenoir, Yadvinder Malhi, Owais Ahmed Malik, Maxence Martin, Iain McNicol, Milutin Milenkovic, David Minor, Ed Mitchard, Vítězslav Moudrý, Helene C. Muller-Landau, Erik Næsset, Anuttara Nathalang, Jean Pierre Ometto, Masanori Onishi, Yusuke Onoda, Petri Pellikka, Henrik Persson, Matheus Pinheiro Ferreira, Pierre Ploton, Suzanne M Prober, Farhadur Rahman, Parvez Rana, Maxime Réjou-Méchain, Jannika Schäfer, Cornelius Senf, Aurélie Shapiro, Dmitry Schepaschenko, Guochun Shen, Miles Silman, Thiago Silva, Jenia Singh, Ferry Slik, Jonas Stillhard, Anto Subash, Ryuichi Takeshige, Shengli Tao, Evan Tenorio, Timo Tokola, Piotr Tompalski, Nithin Tripathi, Ruben Valbuena, Riccardo Valentini, Ronald Vernimmen, Greg Vincent, Jörgen Wallerman, Wan Shafrina Wan Mohd Jaafar, Yongcai Wang, Hannah Weiser, Joanne White, Lukas Winiwarter, Michael Wulder, Zuoqiang Yuan, Katherine Zdunic, Yelu Zeng, Houxi Zhang, Jian Zhang, Zhiming Zhang, Tommaso Jucker

## Abstract

Woody canopies regulate exchanges of energy, water and carbon, and their three-dimensional (3D) structure supports much of terrestrial biodiversity. Remote sensing technologies such as airborne laser scanning (ALS) now enable the 3D mapping of entire landscapes. However, we lack the large, harmonized and geographically representative ALS collections needed to build a global picture of woody ecosystem structure. To address this challenge, we developed the Global Canopy Atlas (GCA): 3,458 ALS acquisitions transformed into standardized and analysis-ready maps of canopy height and elevation at 1 m^2^ resolution. The GCA covers 56,554 km^2^ across all major biomes. 19% of this area has been scanned multiple times, and 87% of all GCA products are openly available, covering 95% of the total area. To showcase its wide range of applications, we applied the GCA in three case studies. First, we validated three global satellite-derived canopy height maps, finding poor performance at native resolution (1-30 m, R^2^ < 0.38) and moderate performance at 250 m resolution (R^2^ < 0.65). Second, analyzing global patterns in canopy gap size frequency we discovered an unexpectedly large variation of power law exponents from branch to stand level (α = 1.52 to 2.38), pointing to a fundamental scale-dependence of forest structure. Third, we developed a framework to standardize forest turnover quantification from multi-source, multi-temporal ALS. In a temperate forest in North America it revealed that 21% of canopy gaps closed within 12 years of opening and would thus be missed by infrequent monitoring. As demonstrated by these case studies, the GCA provides a novel data source for ecologists, foresters, remote sensing scientists and the ecosystem modelling community that substantially advances our ability to understand the structure and dynamics of woody ecosystems at global scales.

## 1. Introduction

Over the past two decades, airborne laser scanning (ALS) has become integral to the environmental sciences (Beland et al., 2019; Eitel et al., 2016). Laser pulses emitted from aircraft penetrate deep into vegetation canopies, and – by reflecting off leaves, branches and trunks – create a three-dimensional (3D) image of vegetation structure. This information is now routinely used for mapping forest resources (Fassnacht et al., 2024; Næsset et al., 2004), microclimates (Gril et al., 2023) and biodiversity (Davies & Asner, 2014; Toivonen et al., 2023). It is also central to the upscaling of vegetation properties such as aboveground carbon stocks (Asner et al., 2014; Coops et al., 2021; Ometto et al., 2023; Xu et al., 2017). At submetric accuracy, ALS reveals an ecological complexity that we are only just finding a language for (Atkins et al., 2023; Lines et al., 2022), and, with frequent, automated surveying on the horizon (Besson et al., 2022), it can help uncover the fundamental processes of how organisms, communities and their biomass develop in space and time (Battison et al., 2024; Jackson et al., 2024; Næsset et al., 2013). These insights will enable us to train a new generation of satellite monitoring tools (Kellogg et al., 2020; Quegan et al., 2019), calibrate high-resolution vegetation models (Fischer et al., 2020; Shugart et al., 2015), and shape conservation policies (Gonzalez et al., 2023). However, to build a comprehensive picture of the 3D structure and dynamics of ecosystems, we must move beyond single-time, single-site studies and harmonize ALS data and derivatives at global scales (Valbuena et al., 2020).

ALS data are increasingly available for this purpose. Most countries in Europe and North America now have substantial ALS coverage, with wall-to-wall acquisitions in many areas at regional scale or below (Moudrý et al., 2024; White et al., 2025), and facilities such as OpenTopography (https://opentopography.org), 3DEP (https://www.usgs.gov/3d-elevation-program) or ELVIS (https://elevation.fsdf.org.au) consolidate ALS data from a wide range of sources in the United States, New Zealand and Australia. As an alternative to wall-to-wall coverage, targeted ALS campaigns have been employed to collect representative samples over large biomes (Wulder, White, Nelson, et al., 2012). Examples include the Amazon (Ometto et al., 2023), the Congo Basin (Xu et al., 2017), Borneo (Melendy et al., 2018), Peru (Asner et al., 2014), and Canada’s boreal forests (Wulder, White, Bater, et al., 2012), as well as data-rich ecological “supersites” (Kampe, 2010; Karan et al., 2016; White et al., 2019). However, despite these advances, there is still a distinct lack of harmonization at global scales (Stereńczak et al., 2020). Three issues stand out in particular. First, the acquisition of ALS varies considerably across locations and projects. They are acquired in different seasons, with different instruments, at different altitudes, and often without consistent reporting standards – all of which can introduce substantial uncertainties into assessments of ecosystem 3D structure (Almeida et al., 2019; Fischer et al., 2024; Joerg et al., 2012; Næsset, 2005, 2009; Okyay et al., 2019; Riofrío et al., 2022; Roussel et al., 2017). Second, the processing of ALS data varies widely. It lacks minimum quality specifications and is carried out with different algorithms, for different aims, and often with little documentation. Ecologically relevant quantities such as canopy height, cover, and density can therefore come with errors that are difficult to correct for and make robust comparisons across studies difficult if not impossible (Fischer et al., 2024; Kissling & Shi, 2023; LaRue et al., 2022; Quan et al., 2021; Vincent et al., 2023; Zhang et al., 2024). Third, the accessibility of ALS data and derived products is inconsistent, with data provided in different formats, with varying data sharing policies and access protocols. It requires specialist knowledge and considerable time to find, download and harmonise ALS data, which substantially slows progress towards global syntheses.

To tackle these challenges, we here present the Global Canopy Atlas (GCA), a harmonised database of 3,458 ALS-derived maps of canopy height and elevation at 1 m^2^ resolution, covering 56,554 km^2^ of the world’s woody ecosystems. Its ALS samples have been processed using a harmonised pipeline designed to enable robust comparisons across acquisitions (Fischer et al., 2024). Products include several digital terrain models (DTMs, ground elevation), digital surface models (DSMs, total surface elevation), and canopy height models (CHMs, top canopy height), all delivered in an easy-to-share GeoTIFF format. Products are supplemented with a range of ancillary spatialized data quality layers and comprehensive metadata recording everything from acquisition dates, instrumentation, flight specifications and geolocation. 87% of the GCA products will be made freely available through the European Space Agency’s Multi-Mission Algorithm and Analysis Platform (MAAP), including thorough documentation and links to the original ALS data sources. In doing so, the GCA provides a novel resource for ecological research, global change studies, and the environmental sciences more broadly. Below, we describe its assembly and contents before demonstrating its range of applications through three case studies that focus on (1) the validation of global satellite-derived canopy height models, (2) global patterns in canopy gap size frequency, and (3) forest turnover quantification from multi-temporal ALS data.

## 2. Database construction

### 2.1 Data aggregation

To assemble the GCA, we searched and documented ALS acquisitions across the globe that could be converted into high-quality maps of terrain and canopy height. We employed a broad definition of ALS data, including acquisitions from airplanes and drones (fixed-wing and rotor-based). We did not include any form of terrestrial laser scanning (TLS) or mobile laser scanning (MLS).

We followed a number of guidelines during the assembly of the GCA. First, we put a focus on landscapes with natural woody vegetation cover, i.e., areas without large-scale human interventions such as urban and agricultural zones or plantations of introduced species. Second, we prioritised representativity and balanced sampling in geographical and environmental space. This means, when downloading data from publicly available ALS collections, we selected subsets covering areas of at least 1 km^2^, ideally 5-10 km^2^, a wide geographic range, and a diversity of ecosystem types. Third, we only included acquisitions where raw ALS data – so-called “point clouds” – were available, so that a standardized processing pipeline could be applied (Fischer et al., 2024). Fourth, we filtered data according to a set of minimum quality standards. These included a target sampling density of at least 2 pulses m^-2^ (based on metadata information), compliance with the LAS format standard (https://www.asprs.org/wp-content/uploads/2019/07/LAS_1_4_r15.pdf, last accessed on 26 August 2025), and the availability of metadata for flight configuration and acquisition dates. We also prioritised “leaf-on” over “leaf-off” acquisitions for ecosystems that are deciduous. This meant not including a wide range of openly available wall-to-wall ALS data, such as available through governmental agencies, as these are often insufficiently documented, low in pulse density or from winter acquisitions. We note, however, that these were broad guidelines. We made exceptions to improve geographic coverage, or to sample sites of particular ecological interest. Therefore, the GCA also includes acquisitions from heavily anthropogenically modified landscapes, with densities <2 pulses m^-2^, or with imprecise metadata on flight dates. Areal coverage can reach several 100 km^2^ in national parks, but also go down to a few hectares for drone acquisitions.

Using these guidelines, we assembled the GCA from a wide range of sources. An overview over all datasets that we included is provided in Table S1. We downloaded open collections from dedicated ecosystem observatories such as the Brazilian Sustainable Landscapes program (https://www.paisagenslidar.cnptia.embrapa.br/, this and all following links accessed on 26 August 2025), the United States’ National Ecological Observatory Network (Kampe, 2010) and Australia’s Terrestrial Ecosystem Research Network (TERN, http://portal.tern.org.au/). We supplemented them with ALS campaigns such as the Estimativa de Biomassa na Amazônia (EBA) initiative (https://www.ccst.inpe.br/projetos/eba-estimativa-de-biomassa-na-amazonia/), the World Wildlife Fund’s (WWF) acquisitions over the Democratic Republic of Congo (Xu et al., 2017) and individual research projects. We complemented targeted ALS acquisitions with openly available data from country-specific websites (e.g., https://geoservices.ign.fr/lidarhd in France) and collections such as OpenTopography, with a focus on protected areas and those with a low human footprint. Additionally, we contacted research groups who had acquired ALS data over large environmental gradients or at specific research sites (Stereńczak et al., 2020) and invited them to contribute to the database.

### 2.2 Data processing and quality checks

All data processing was carried out with the same standardized pipeline, which is described in detail in Fischer et al. (2024). The pipeline is fully automatic and based on a few key software tools, mainly the laser scanning software LAStools and the R packages data.table (Dowle & Srinivasan, 2025), terra (Hijmans, 2025), sf (Pebesma, 2018), and lidR (Roussel et al., 2020). It carries out several pre- and post-processing steps, such as the splitting of ALS data into coherent spatial units as well as noise filtering, deduplication, and ground classification. For each spatial unit, the output is a set of DTMs, DSMs, and CHMs, as well as several types of ancillary layers for quality control. For most applications, we recommend the default DTM and a CHM derived either from a triangulated irregular network (TIN) or a locally adaptive version of the spikefree algorithm (Khosravipour et al., 2016), both of which performed well in a robustness assessment (Fischer et al., 2024). An overview of the different types of layers generated by the pipeline is provided in Table 1, with further information in Table S2.

**Table 1:**
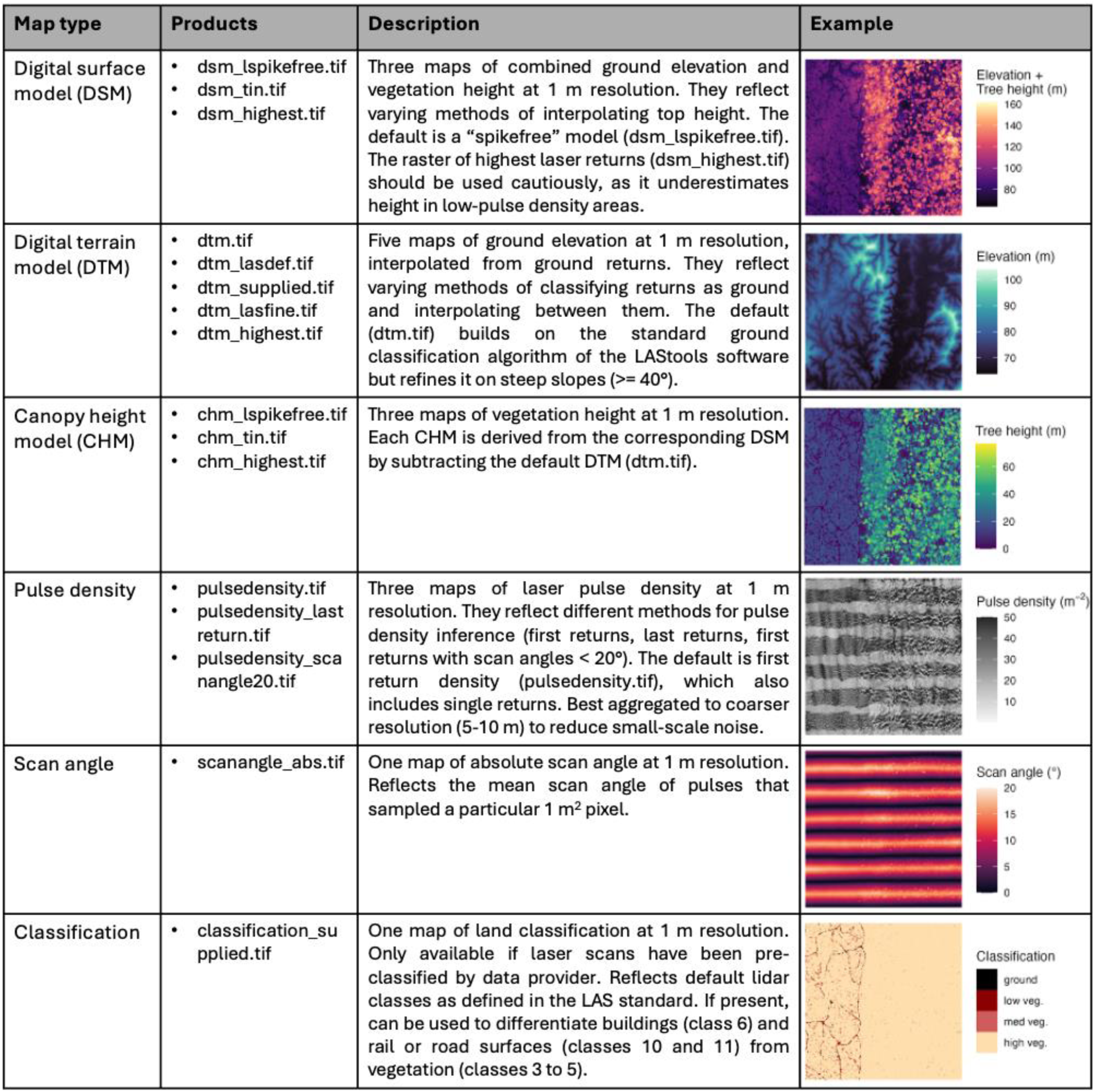
Overview of types of spatial layers provided as part of the Global Canopy Atlas. For several of the derived layers (e.g., CHMs and DTMs) we provide a range of alternative options generated using different algorithms which are described in Table S2 and in Fischer et al. (2024). We briefly summarise these in the description provided above and highlight the suggested default maps for most purposes. Sample data come from a 2020 acquisition at the Sepilok Forest Reserve in Malaysian Borneo and show a 1 km^2^ tile of where the reserve (right) borders an oil palm plantation (left).

Throughout processing, we kept all parameters constant across ALS acquisitions, with only few manual adjustments. For example, pre-classified noise points (LAS class 7 and 18) were always excluded, but for some datasets we also excluded custom noise classes defined by the data providers. These noise classes reflected manual corrections of scan artefacts or dense noise that would not be captured by automatic noise detection. In terms of geolocation, our default approach was to retain any provided metric coordinate system to ensure consistency with metadata. However, if acquisitions were provided in imperial units (ft), we converted them into SI units and Universal Transverse Mercator (UTM) coordinates to ensure global comparability. The same approach – retaining the original data configuration where possible – was applied to vertical geo-referencing, so acquisitions were generally processed with the provided ellipsoid or geoid reference. While this does not affect woody ecosystem structure quantification (CHMs), there may be a location-dependent offset between ellipsoidal and geoidal datum up to around 100m (DTMs and DSMs).

Data quality standards varied across data providers and instrumentation, with errors ranging from negligible (isolated noise points) to considerable but repairable (faulty geolocation, flight line misalignment) to substantial and beyond repair (insufficient canopy penetration or flight line overlap). To process large numbers of ALS acquisitions, we developed automatic procedures for quality assurance and control. The vast majority of these quality checks are directly embedded in the processing pipeline and provided as ancillary layers alongside the DTMs and CHMs (Table 1, Table S2). This gives users flexibility in choosing appropriate quality levels before and during data analysis. General quality layers include pulse density rasters that enable users to identify areas of poor sampling (pulsedensity.tif), derived pulse density masks (mask_pd02.tif and mask_pd04.tif, subsetting to regions with >2 or 4 pulses m^-2^, respectively), rasterizations of scan angles (scanangle_abs.tif), which enable identification of flightline patterns, as well as masks for potential high noise or cloud points (mask_cloud.tif). In addition, we provide several alternative DTM layers to assess robustness of ground detection, which is a major source of uncertainty on steep slopes and under dense forest cover (Stereńczak et al., 2016), as it requires both dense sampling and high laser power. Users can investigate how and where DTMs disagree or use pre-computed DTM masks that highlight areas with high uncertainty (mask_unstabledtm.tif) and those on steep slopes where features such as cliffs hamper robust ground classification (mask_steep.tif).

### 2.3 Processing reports

Each processed acquisition is accompanied by an automatic two-page PDF processing report (Fig. S1, also provided as separate pdf in the Supplementary). The first page provides an informal quality check, showing the acquisition outline overlaid on local maps from OpenStreetMap and optical imagery from ESRI, downloaded via the R basemaps package (Schwalb-Willmann, 2024). The second page provides a more detailed and quantitative overview. It shows a pulse density rasterization where areas with pulse densities <2 m^-2^ are flagged to provide a rapid assessment of lower-quality scan areas, as well as a DTM reliability mask (mask_unstabledtm.tif). It also shows the timing of the acquisition with respect to vegetation greenness/phenology (NDVI and EVI) and snow cover obtained from the MODIS Aqua and Terra sensors (Didan, 2025). Finally, the second page of the processing report also compares ALS-derived products with three satellite-derived layers: the Copernicus World DSM at 90 m resolution (European Space Agency, 2024), top canopy height (RH98) estimated from GEDI/ICESat-2 at 100 m resolution (Saatchi & Favrichon, 2023), and global canopy height derived from Landsat at 30 m resolution (Potapov et al., 2022). ALS data provide more accurate representations of terrain and vegetation structure than satellites, but we have found the comparison useful for practical purposes. First, it helps identify potential geo-location errors. For example, correlations between ALS DSMs and the Copernicus World DSM are generally high (R^2^ > 0.9), so low correlations can indicate assignment to the wrong UTM zone or constant horizontal offsets. Second, the comparison also provides a quick indicator of how well common satellite products represent local vegetation structure, including temporal offsets, and thus offers an insight into upscaling opportunities, a key application of ALS data (Coops et al., 2021).

## 3. Database overview

The GCA database consists of 3,458 ALS acquisitions converted into maps of vegetation height and terrain elevation at 1 m^2^ resolution (Fig. 1a). These acquisitions vary considerably in size from 0.1 to 203.7 km^2^ (95% interval). The vast majority (85%) cover 1 km^2^ or more, and the overall median area is 8.5 km^2^. All acquisitions together cover 56,554 km^2^ of land area spread across 56 countries and 224 ecoregions (Dinerstein et al., 2017). About 57% of this area (31,554 km^2^) is in intact forest landscapes (Potapov et al., 2017) or has some form of protection designation (World Database of Protected Areas). About 95% of the area – represented by 87% of the acquisitions – is covered by openly available GCA products. The total file size of GCA derived products is 3.2 TB.

**Fig. 1:**
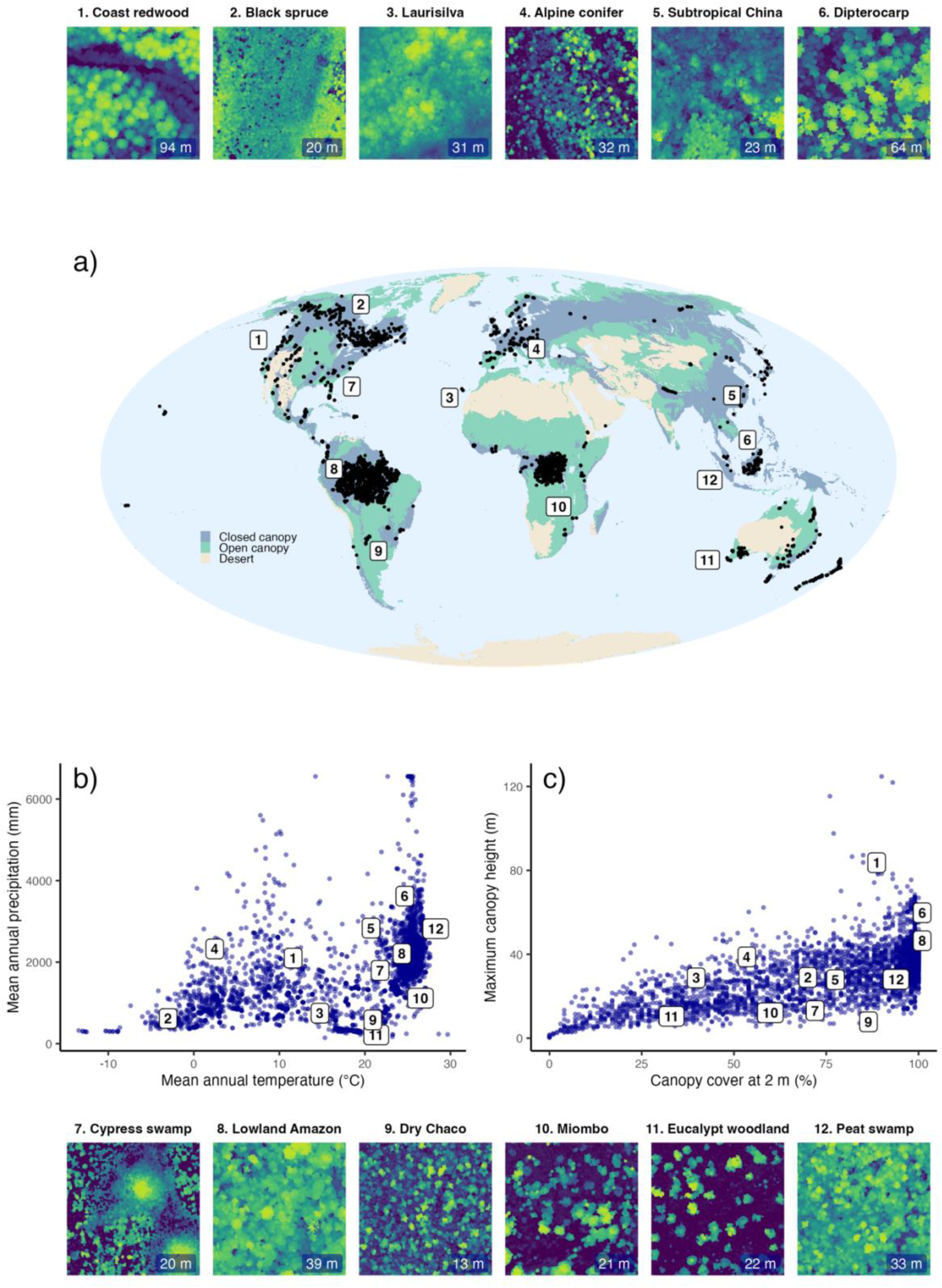
The Global Canopy Atlas. The distribution of each of the 3,458 individual ALS acquisitions is shown (a) on a map of the world, (b) in the environmental space, and (c) in terms of woody ecosystem structure. The top and bottom rows show sample canopy height models (CHMs, 200×200 m) for 12 sample sites that span a wide variety of ecosystems across the globe. Colours, ranging from dark blue to yellow with increasing height, are rescaled to fit the range of heights observed in each system, with the maximum height of the vegetation patch shown in the bottom right corner. The location of these 12 sites is also shown in geographic, environmental and structural space in panels (a–c) using numeric labels. Closed and open canopy ecosystem layers shown in (a) were derived by aggregating biomes in the Ecoregions2017 dataset (Dinerstein et al., 2017), for details cf. Table S3. Climatic variables in panel (b) were extracted from 1-km CHELSA climatologies for the years 1981-2010 (Karger et al., 2017) and averaged across the entire scan area. Maximum canopy height in panel (c) is defined as the 99^th^ percentile of canopy height across the entire scan area. Canopy cover at 2 m is calculated as the percentage of CHM pixels ≥ 2 m.

Geographic coverage of the GCA extends to all biomes and biogeographic realms (Fig. 1). The total number of acquisitions in the tropics is nearly twice as large (1,950) as in temperate regions (1,102) and six times as large as in the boreal zone (336). However, temperate and boreal acquisitions tend to cover larger areas and consequently the GCA comprises a total scan area of 20,271 km^2^ (36% of the total) in the tropics, 27,543 km^2^ in the temperate zone (49%, including Mediterranean forests), and 6,898 km^2^ in boreal regions (12%). A few acquisitions (70) cover other biomes, such as mangroves or highly xeric systems with little woody canopy cover (1,842 km^2^, 3%).

In terms of bioclimatic space, the GCA spans a large gradient of mean annual temperatures, going from as low as -10 °C up to 30 °C, and ranges in precipitation from less than 100 mm a year to over 6,000 mm (Fig. 1b). The woody ecosystem structures sampled in the GCA are also highly diverse, from open systems with canopy cover of less than 10% and trees that never exceed 10-20 m, to dense, tall forests where canopy cover reaches close to 100% and trees can exceed 100 m in height (Fig. 1c).

In addition to the extensive sampling of geography, environment and woody ecosystem structure, the GCA covers large tracts of land that have been scanned more than once (Fig. S2). The area covered by repeated ALS acquisitions is 10,532 km^2^ (19% of the total), but because several of these sites were scanned on multiple occasions the cumulative area of repeat acquisitions is 33,954 km^2^. This is more than 50% of the single-scan area coverage in the GCA, and brings the total scan area to 90,508 km^2^. Repeat acquisitions are available for all biomes, with the majority of the area covered by acquisitions on NEON sites in the United States of America (62%). The median interval between acquisitions is 1.2 years (95% range from 0.6 to 10.0 years).

In terms of sampling density and data quality, 93% of GCA acquisitions have an average pulse density >2 m^-2^, and the median density across all acquisitions is 5.1 m^-2^. There is, however, a large spread around these values, with a 95% interval ranging from 1.2 to 123.1 m^-2^. Temperate acquisitions in the GCA generally have higher pulse densities (median of 8.6 m^-2^) than their tropical (4.9 m^-2^) or boreal (3.5 m^-2^) counterparts, and drone acquisitions have much higher densities (up to 1,000 m^-2^) than airplane and helicopter-based ones, but these high pulse density acquisitions only account for a small fraction of the database (49 acquisitions > 300 pulses m^-2^).

In terms of temporal coverage, acquisition dates range from 2006 to 2024, with 90% acquired between 2012 and 2022. More than 74% of acquisitions were completed within a 1-week period (>94% within three months and >99% within six months). Most acquisitions have been acquired near or at peak EVI (87% of acquisitions) and NDVI (92%), with no clear geographic biases in acquisition timing. Uncertainty in ground classification is generally low. The average acquisition has topographic uncertainties over less than 2% of its area, and only 51 acquisitions (∼1.5%) have topographic uncertainties over more than 20% of the scanned area.

## 4. Case studies

All three case studies are based on analyses of CHMs generated using the locally adaptive spikefree algorithm (chm_lspikefree.tif), as described in Tables 1 and S2. R code and ancillary data to replicate them are archived on Zenodo (https://doi.org/10.5281/zenodo.16987211).

### 4.1 Case study 1: How accurate are satellite-derived global canopy height models?

Large-scale models of canopy height derived from satellite data are increasingly being used to map the structure and aboveground carbon storage of woody ecosystems (Liu et al., 2025; Schwartz et al., 2023; Wagner et al., 2025). At global scale, height maps are mostly created by combining wall-to-wall optical satellite imagery with laser scanning data acquired from space-borne or airborne sensors using machine learning. Three global examples include: (1) Potapov et al. (2021, 2022), who used Landsat imagery together with GEDI and ICESat-2 shots to map canopy height at 30 m resolution, (2) Lang et al. (2023), who fused GEDI data and Sentinel-2 imagery to map canopy height at 10 m resolution; and (3) Tolan et al. (2024), who trained models on ALS data and predicted height from Maxar imagery at 1 m. However, while these models generally replicate global-scale gradients in canopy height, questions persist about their accuracy at landscape scale (Besic et al., 2025; Moudrý et al., 2024; Wagner et al., 2024). Optical satellite data have limitations both in open systems, where they underestimate tree cover (Brandt et al., 2020), and in closed canopy forests due to saturation effects (Mutanga et al., 2023). To address these concerns, we assessed the landscape scale accuracy of all three global canopy height models against ALS data from the GCA.

We compared the three satellite-derived height maps against ALS-derived CHMs from across 20 landscapes that had not been used to train the models. These ALS acquisitions span forest types with distinct 3D structures and were collected within 1-2 years of the satellite data used to generate the global models (Table S1.1). Agreement between ALS-derived products and the global maps was quantified by calculating the coefficient of determination (R^2^) and the relative root mean square error (RMSE), i.e., the RMSE standardized by the mean canopy height. Since the global canopy height models differed in resolution, we first converted them to 1-m resolution via nearest-neighbour resampling and then calculated the R^2^ and RMSE across a wide range of resolutions between 1-250 m. By default, we averaged canopy height at each scale, but global models may represent different aspects of height (Besic et al., 2025), so we also calculated the R^2^ and RMSE for maximum height (99th percentile) at each resolution.

The three global canopy height models generally showed poor agreement with the reference ALS data (Fig. 2). At their nominal resolutions, R^2^ ranged from 0.28 (Lang, 10 m) to 0.31 (Tolan, 1 m) to 0.38 (Potapov, 30 m). The corresponding errors were lowest for Tolan (relative RMSE = 95%), higher for Potapov (112%) and highest by some margin for Lang (189%). However, agreement depended strongly on resolution and tended to increase at coarser scales (Fig. 2a-b). When assessed at a common resolution of 30 m, the Tolan model clearly outperformed the other two models (R^2^ = 0.54 and RMSE = 58% compared to 0.38 and 112% for Potapov, and 0.39 and 181% for Lang). The same was true at 250 m, where the Tolan model had an R^2^ of 0.65 and an RMSE of 43%, compared to the poorer 0.56 and 93% of Potapov and 0.56 and 170% of Lang. When comparing maximum heights instead of means, the differences in RMSE between models narrowed (57-69% at 30 m, 40-56% at 250 m; Fig. S1.1), but the accuracy of the predictions worsened (R2 = 0.38-0.55 at 250 m).

**Fig. 2.**
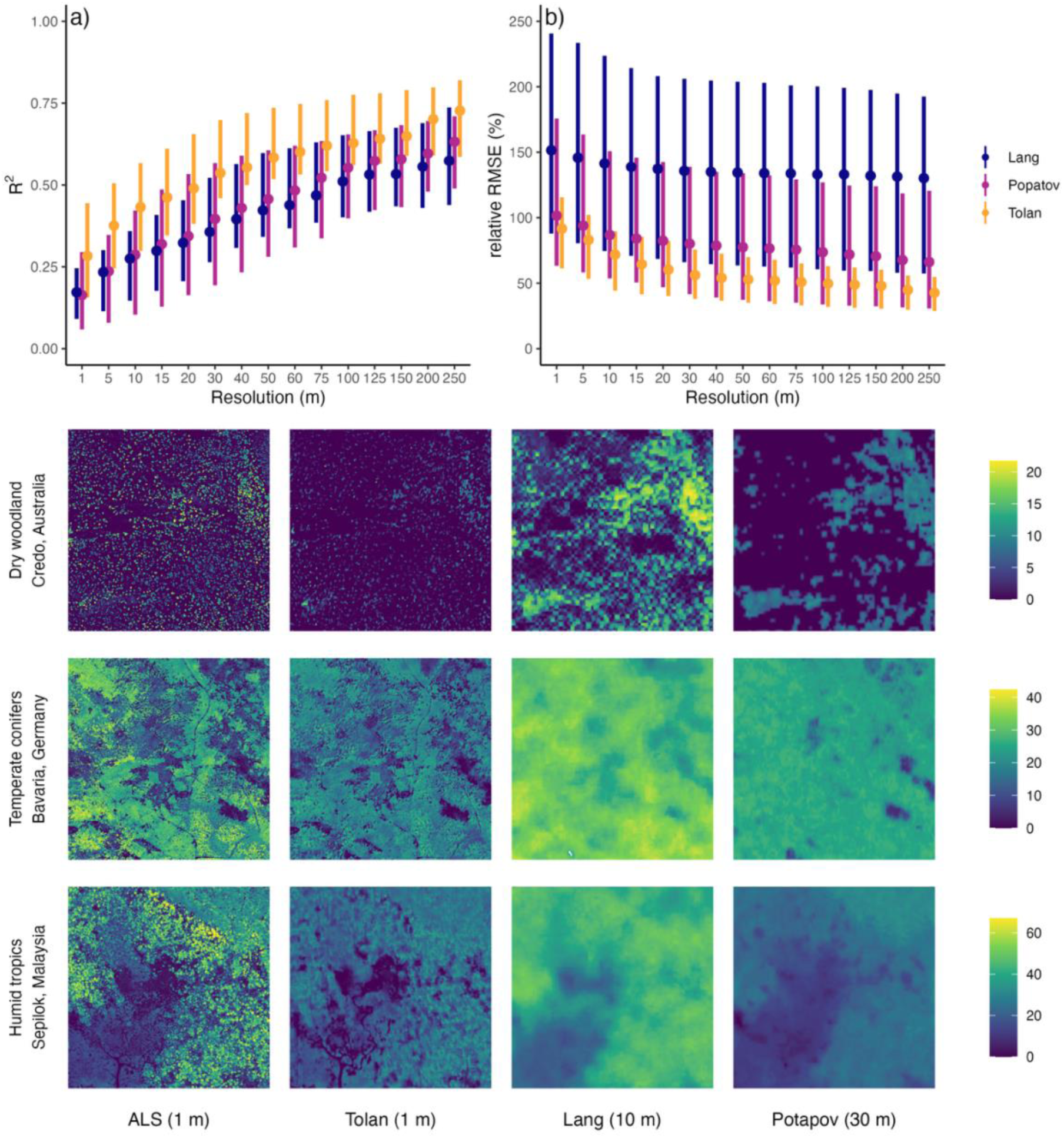
Accuracy of global canopy height models. Panels (a-b) show the agreement between the three global canopy height models and ALS data across 20 landscapes at different spatial resolutions on the basis of R^2^ and relative RMSE. Rasters are aggregated to different resolutions via averaging, with ALS data assumed to provide the true value. The distributions at each resolution are summarized via median values (dots) and 25^th^ and 75^th^ percentiles (error bars). Note that some data points exceeded 250% in relative RMSE, so have been excluded from the visualized range in panel (b). The panels below show sample canopy height models (2×2 km) from 3 landscapes, including the reference ALS product at 1 m resolution. The height range of these maps is provided by the scale bars on the right. Examples of landscapes with particularly large errors for each of the three models can be found in Figs S1.4-1.6, an alternative assessment based on maximum canopy height in Fig. S1.1.

Global scale canopy height products are expected to be less accurate than costly high-resolution airborne laser scans, but our results highlight strong deficiencies in currently available global canopy height maps. While these products broadly mirror global height trends (Fig. S1.3), they miss most of the variation in canopy height within landscapes at their nominal resolutions (1–30 m). Even at 250 m resolution and for the best performing model (Tolan), the mean relative error was >40% of the reference height and reached as high as 170% in certain landscapes. This makes it near-impossible to capture within-landscape variation in canopy height with reasonable accuracy (Moudrý et al., 2024). We found that some models, such as the Lang model, predicted maximum canopy height better than mean height (Besic et al., 2025), but errors still remained above 40%, and R^2^ values were even lower than for mean height. Across the vast majority of sites, the high-resolution approach of Tolan et al. (2024) was superior to other global maps tested here, with higher R^2^ and lower RMSE (Fig. 2), and better representation of landscape variability (Fig. S1.3). This may be due to an improved detection of trees from high-resolution imagery (Brandt et al., 2020) as well as the training on ALS data, which offer better resolution and geolocation than spaceborne lasers.

That being said, the Tolan height map also contained notable errors and uncertainties, underpredicting height at a tropical site in Amazonia by more than 20 m and consistently misrepresenting cloud cover as canopy gaps in tropical regions (Figs. S1.4-6). Global canopy height modelling is not even 15 years old (Simard et al., 2011) and we expect it to improve massively in the coming years, especially with the integration of new data sources (Quegan et al., 2019). It seems, however, that we are still a long way from operational products for global forest structure monitoring at medium to high resolution (1-30 m). Combining high-resolution satellite imagery with ALS for model training – as Tolan et al. (2024) have done – seems a promising approach. But it would require a more careful screening of input data (e.g., removing cloud artefacts) and a wider range of ALS data for calibration. A recent canopy height model for Amazonia shows considerable potential in this respect (Wagner et al., 2025). We anticipate that the GCA, with its robust canopy height products available across all biomes, can serve as a benchmark for the calibration and validation of such models.

### 4.2 Case study 2: Does the frequency of canopy gap sizes follow power laws?

Treefall gaps are an essential component of forest structure and dynamics (Franklin et al., 1987). They reinitiate succession, create diverse forest microclimates and habitats (Ritter et al., 2005), and shape species composition (Denslow, 1987). As legacies of forest disturbance, they are key proxies for measuring canopy and carbon turnover (Dalagnol et al., 2021; Hunter et al., 2015; Jucker, 2022) and indices of how humans reshape ecosystem structure (Zhang et al., 2023). Over the last two decades, large-scale ALS surveys have revealed a fundamental pattern: small gaps are legion, and large gaps are rare. Regional-scale studies have often found that these heavy-tailed gap size frequency distributions (GSFD) exhibit power law scaling (Goodbody et al., 2020; Kellner & Asner, 2009), possibly with a narrow range of exponents (Asner et al., 2013). If true at global scale, this would imply an astonishing level of generalization: (1) that forest dynamics are governed by multiplicative processes, (2) that these processes show little variation from the boreal to the tropics and (3) that they are scale-invariant, i.e., small-scale disturbances such as branchfalls follow similar rules as large-scale disturbances such as those associated with wildfires and storms. However, not all studies show power law scaling (Araujo et al., 2021), and scale invariance has never been systematically tested at global scales. Comparisons between regional studies are complicated if not impossible, as the inferred exponents depend on how CHMs are derived (Fischer et al., 2024; White et al., 2018), how gaps are defined (Asner et al., 2013; Lobo & Dalling, 2014), and how power laws are fitted (Hanel et al., 2017; Wedeux & Coomes, 2015). Since power laws are difficult to distinguish from other heavy-tailed distributions (Newman, 2005; Taubert et al., 2013), it is an open question whether GSFDs follow power laws across forest types, spatial scales, and the range of gap sizes.

To systematically tackle this question, we chose a subset of high-quality CHMs from the GCA with a minimum of 70% canopy cover at 2 m and divided them into 1 km^2^ cells using a flexible Voronoi tessellation (n = 13,300, details on selection in Supplementary S2.1). For each 1 km^2^ cell we calculated the mean canopy height (µ_CHM_) and defined gaps as contiguous areas (rook neighborhood) below a height threshold of δ_CHM_ = 0.5 × µ_CHM_ and no larger than 100,000 m^2^ (10% of the reference area). This excluded some larger gaps, but was not expected to alter power law assessments below 100,000 m^2^. Gap size was calculated as the number of 1 m^2^ pixels in a gap (Kellner & Asner, 2009), and discrete power laws were fitted with the poweRlaw R package (Gillespie, 2015). Tests for power law behaviour exist, but reduce a continuous spectrum of deviations to binary hypothesis testing and are difficult to interpret (Hanel et al., 2017). We therefore used an alternative heuristic that quantifies deviations from scale invariance (Fischer & Jucker, 2024). Under ideal power law behaviour, the scaling exponent α is scale-invariant. If a power law is fit to the full range of gap sizes and to a subset (e.g., only large gaps), α should be approximately identical. If α differs, we can calculate a deviation from power-law scaling Δ_PL_ as the difference between both α values. We here calculated Δ_PL_ over three different scale ranges. First, we calculated α for all gaps from small branch falls ≥1 m^2^ up to 100,000 m^2^ (α_branch_). This is the lower cutoff used in Kellner & Asner (2009). The mean area of all gaps within that range was 32.8 m^2^ (95% range: 9.4 to 91.8 m^2^). Second, we calculated α only for single treefall gaps or larger (α_crown_). Crown area scales with tree height (Jucker et al., 2022), so we approximated the typical tree crown area as circle with a diameter equivalent to the height cutoff δ_CHM_ in each 1 km^2^ cell (0.25 × δ_CHM_ ^2^ × π). The corresponding mean gap area was 420.4 m^2^ (60.3 to 1,446.7 m^2^). Finally, we calculated α only for the tail end of the gap size frequency distribution (α_stand_). For this, we calculated the cutoff as the midpoint between the typical tree crown size and the size of the largest disturbances (95^th^ percentile of gap sizes). The mean gap area was 1,830.9 m^2^ (210.0 to 5691.1 m^2^). Deviations were then quantified as Δ_PL, branch_ = α_crown_ – α_branch_ and Δ_PL, crown_ = α_stand_ – α_crown_ and calculated separately for boreal forests (n = 501), temperate forests (n = 3,903) and tropical forests (n = 8,896). The robustness of results was tested through several sensitivity analyses (Figs. S2.2-2.7).

We found that the power law exponent α was highly scale dependent, indicating that GSFDs do not follow power law behaviour (Fig. 3). When small gaps were included, α was low and narrowly distributed within and across biomes (mean α_branch_ = 1.52; 95% interval = 1.42-1.70). By contrast, when focussing on gaps from treefalls or larger disturbances, exponents were considerably higher and much more variable (α_crown_ = 2.22; 1.73-3.28), even though the gaps excluded in the α_crown_ calculation only made up 2.59% (0.74-5.21) of the total gap area. Across biomes, tropical forests generally had the largest α_crown_ and the largest variation around the mean (2.29; 1.73-3.45), compared to boreal (2.02; 1.76-2.38) and temperate forests (2.08; 1.71-2.61). From branch to crown scale, the typical deviation from power law scaling Δ_PL, branch_ was 0.70 (0.16-1.73) and was highest in the tropics (0.77) and smallest in boreal landscapes (0.49). Exponents increased again, albeit much less, when excluding treefall gaps and only fitting power laws to the largest disturbances (α_stand_ = 2.38; 1.73-3.63 and Δ_PL, crown_ = 0.25; - 0.23-0.91). Qualitatively, our results were robust to methodological choices, with α changing across scales and biomes also for different CHM types (Figs. S2.2–2.5) and when applying fixed thresholds to define gaps (< 2 m) and minimum gap size (≥ 9 m^2^, Figs. S2.6–2.7).

**Fig. 3:**
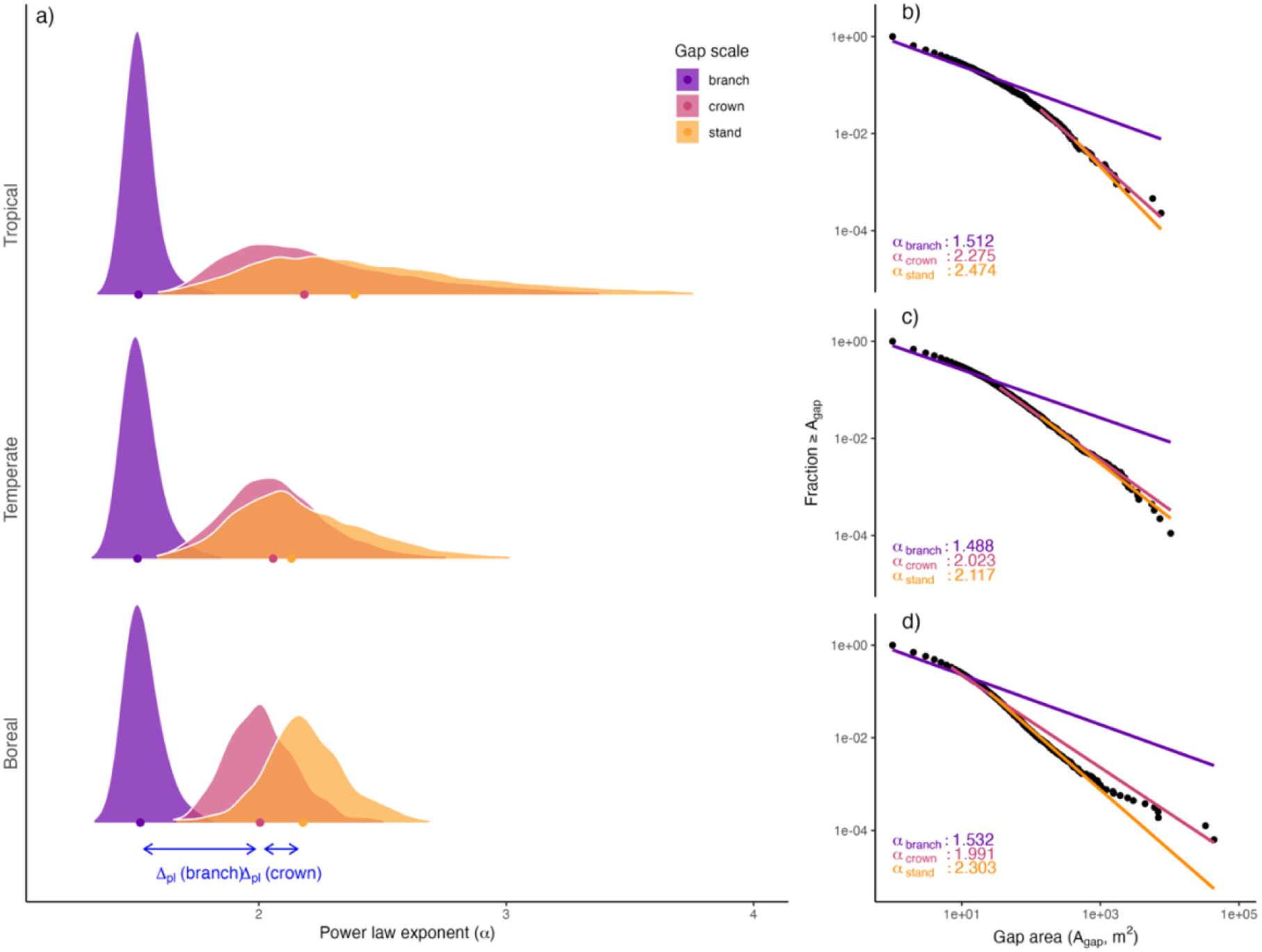
Deviations from power law scaling across major biomes. Panel (a) shows the distribution of power law exponents of gap size frequency distributions (α) for 1 km^2^ cells across tropical, temperate and boreal biomes and coloured by the size ranges over which they are calculated (α_crown_, α_branch_ and α_stand_). Coloured points represent the median values of the distributions. Under true power law behaviour, α should not depend on the size range over which it is calculated, so all distributions should be indistinguishable, with a near-identical median or mean (Δ_PL_ ≈ 0). The righthand column (b–d) shows results from a sample 1 km^2^ cell in each of the three biomes to illustrate how α varies across scales. Each black point represents the contribution of a single gap to the the observed complementary cumulative distribution function (CCDF) of gap sizes, plotted on double-logarithmic scales. Coloured lines show the fitted power laws. We chose a CCDF representation over size frequency plots, as it avoids binning into size classes and noise in the tail end (Newman, 2005). Sample data are from a tropical site in Indonesian Borneo (b), a temperate site in Northwestern United States (c), and a boreal site in Québec, Canada (d) and were selected to be close to each biome’s median in α_branch_ and α_crown_.

Overall, our results provide strong evidence that GSFDs do not follow power laws. The violation of scaling behaviour points to fundamentally different processes shaping forest structure at small and at large scales (Shugart et al., 2010). When GSFDs include small, branch-level gaps, inferred power law coefficients tend to be small and highly consistent across systems (Fig. 3), but at larger scales they exhibit tremendous variability that reflects the “fingerprints” of different disturbance regimes (Jucker, 2022). Violations of fundamental scaling assumptions likely explain why previous studies have come to different conclusions about the typical values and variability of power law coefficients (Asner et al., 2013; Goodbody et al., 2020; Reis et al., 2022), with differences exacerbated by CHM construction (Fig. 3 vs. Fig. S2.2, White et al., 2018). Based on our results (Δ_PL, crown_ << Δ_PL, branch_) and in line with observations in other systems (Newman, 2005), it is possible that GSFDs converge to power law behaviour at the tail end of the distribution, but our case study reveals a fundamental scale dependence across a large size range. A wider range of functions to model GSFDs (Araujo et al., 2021), combined with collections such as the GCA, will help build a more comprehensive picture of global forest structure and how it is shaped by natural and anthropogenic disturbances.

### 4.3 Case study 3: How dynamic are forest canopies?

Accurate monitoring of forest growth and disturbance through time is an essential first step in understanding drivers of forest dynamics and forest responses to global change. Collections such as the GCA bring together ALS data from many sources and years. Many of these acquisitions overlap spatially, which creates the opportunity for large-scale repeat surveying of forests, in particular the monitoring of growth (Yu et al., 2004), disturbance (Huertas et al., 2022), recovery (Krüger et al., 2024), biomass change (Cao et al., 2016; Næsset et al., 2013), structural complexity (Rosen et al., 2024), and tree demographics (Battison et al., 2024). However, estimating change rates from multi-temporal and multi-source ALS comes with its own set of challenges and depends on instrument differences and methodological decisions (Jackson et al., 2024; Leitold et al., 2018). When intervals between acquisitions are short (1-5 years), small biological changes are easily confused with measurement errors (Heiskanen et al., 2024; Riofrío et al., 2022). By contrast, as intervals between acquisitions get longer, disturbances can be missed or underestimated in size due to rapid recovery between acquisitions or repeat disturbances in the same area. These transient disturbances and recovery events parallel the problems of unobserved tree mortality and growth in field inventories with long remeasurement intervals (Kohyama et al., 2019), but have not been explored for repeat ALS surveys. We here leverage a unique timeseries of 8 ALS acquisitions conducted over a period of 12 years at Harvard Forest in the Northeastern United States and propose a new approach to robustly estimate disturbance and recovery rates, instrumentation noise, and transient dynamics.

Harvard Forest is part of the National Ecological Observatory Network (NEON). It is a mixed temperate forest dominated by deciduous broadleaf trees (Quercus rubra, Acer rubrum) and white pine (Pinus *strobus*), with a mean canopy height of ∼17 m and canopy cover of ∼80%. The GCA contains 12 ALS acquisitions for Harvard Forest, but for the purposes of this analysis we focussed on 8 summer acquisitions conducted by NEON (2012, 2014, 2016, 2017, 2018, 2019, 2022, and 2024), as these cover a large area of overlap (∼145 km^2^, all pulse densities > 2 pulses m^-2^). Each acquisition’s CHM was split into tiles of 1 km^2^. For each cell, we defined gaps as contiguous areas (rook neighborhood) below 10 m canopy height. Non-forest areas were defined as those lower than 10 m in canopy height which changed by less than 2 m over 12 years, and masked out. We calculated dynamics between each pair of NEON acquisitions (28 combinations), differentiating disturbance (newly formed gap area as proportion of intact canopy area), and recovery (closed gap area as a proportion of previous gap area; Jackson et al., 2024). Assuming a typical yearly disturbance rate *d* and recovery rate *r*, expressed as proportion of initial canopy and gap area, we derived an analytic relationship between those rates and the observed disturbance and recovery for a given time interval between surveys (Supplementary Material S3.1). This allowed us to separate between-scan noise (*d_0_*and *r_0_* for disturbance and recovery) from biological signal and to estimate unobserved dynamics by comparing observed rates to the theoretical rates that would apply if there were no transient events (no short-term recovery after gap formation, no immediate re-disturbance after closure). We fitted the models via the *brms* R package (Bürkner, 2018, Supplementary Material S3.2) and explored the sensitivity of results to modelling decisions (Supplementary Material S3.3-3.4).

We found near steady-state dynamics at Harvard Forest, with disturbance slightly exceeding recovery (Fig. 4). We estimated a 1-year gap creation rate of *d* = 0.45 ha km^-2^ (95% interval = 0.36-0.64) or slightly less than 0.5% of the intact canopy area (∼90 ha km^-2^), and a 1-year gap recovery rate *r* = 0.41 ha km^-2^ (0.07-1.11), or slightly less than 5% of the disturbed area (∼10 ha km^-2^). After accounting for biases due to between-scan noise (*d_0_* = 1.09 ha km^-2^, 0.64-1.54, and *r_0_* = 0.93 ha km^-2^, 0.68-1.17), we inferred a total of 5.30 ha km^-2^ (4.26-7.34) of newly created gaps and 3.86 ha km^-2^ (0.85-7.26) of recovered canopy area over 12 years. Of the new gaps, an estimated 1.11 ha km^-2^ (∼21%) had closed again by 2024, and 0.1 ha km^-2^ (∼3%) of closed gaps had re-opened after closure. In canopies with higher turnover, we estimated that such transient dynamics could account for up to 57% (short-term recovery of gaps) and 7% (re-disturbance of recovered area), respectively (*d* = *r* = 2.0 ha km^-2^, Fig. S3.1). Our estimates varied slightly, but not fundamentally, when introducing a minimum gap size (*d* = 0.45 ha km^-2^, *r* = 0.34 ha km^-2^, Fig. S3.2) or subsetting to a smaller area where additional acquisitions were available (*d* = 0.58 ha km^-2^, *r* = 0.38 ha km^-2^, Fig. S3.3). Inferences from single pairs of acquisitions were more uncertain (*d* ∼ 0.24-1.11 ha km^-2^ and *r* ∼ 0.05-1.04 ha km^-2^), but the overall averages were consistent (*d =* 0.45 ha km^-2^, *r* = 0.40 ha km^-2^, Fig. S3.4). Estimates of biases due to between-scan noise were confirmed independently via temporally coincident acquisitions (*d_0_*= 0.89 ha km^-2^, 0.19-3.19, *r_0_* = 0.94 ha km^-2^, 0.28-2.82, Fig. S3.5).

**Fig. 4.**
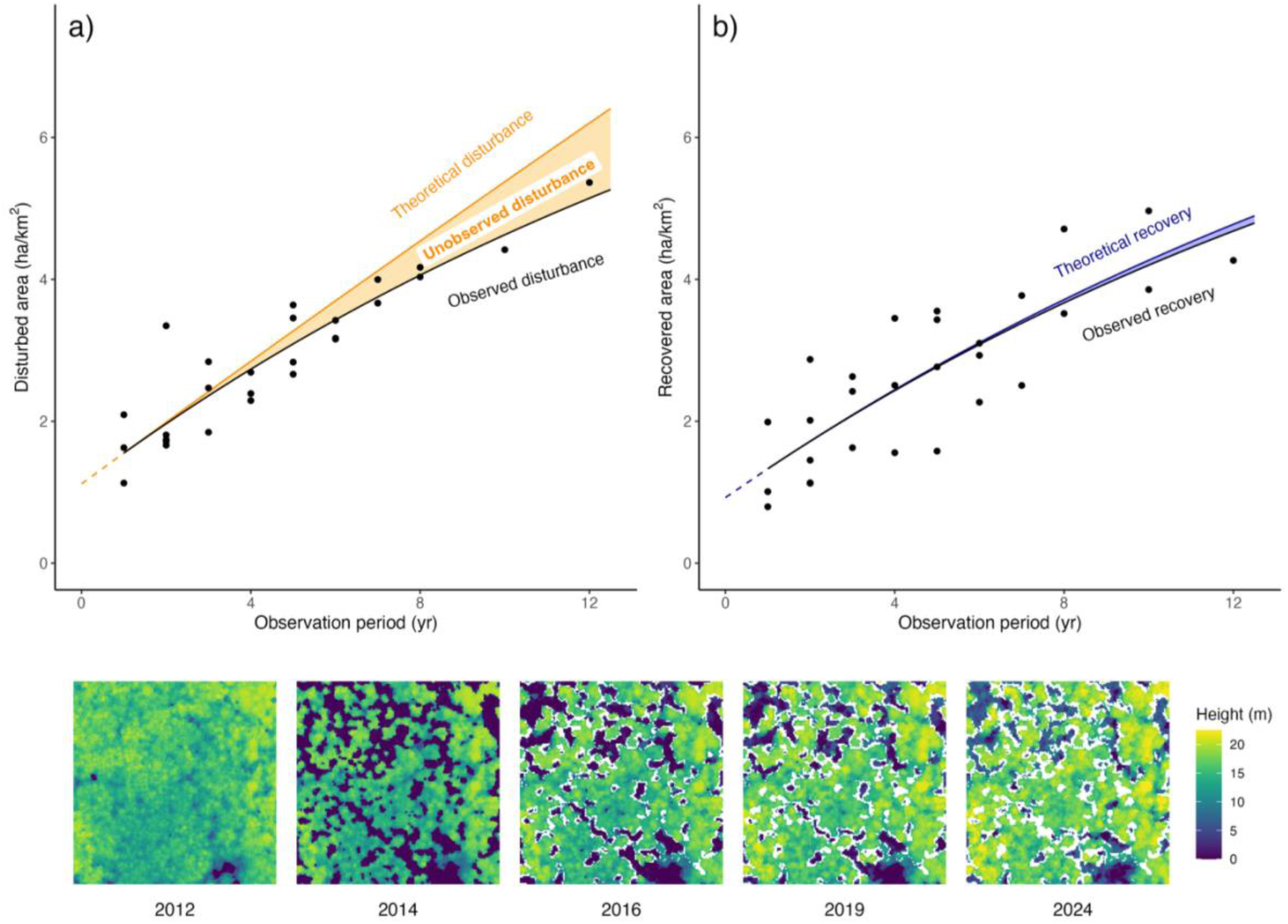
Short-term dynamics at Harvard Forest. Panels show observed rates of (a) disturbance *d* and (b) recovery *r* as a function of the time interval (years) between successive ALS surveys (black points). As the intervals between observations get longer (a), an increasing proportion of disturbances recover before being re-surveyed, leading to an underestimation of true disturbance rates (and equivalently for recovery in panel b). Rates were inferred as proportion of existing canopy and gap area and then converted into area rates per grid cell (ha km^-2^). Black lines are fitted relationships, while coloured lines reflect the theoretical changes that would be expected if there was no short-term recovery of disturbed area (a) and no rapid re-disturbance of recovered area (b). They are calculated by extending *d* and *r* into the future, discounting only for loss of canopy and gap area, respectively. The intercepts *d_0_* and *r_0_* are non-zero, as a proportion of pixels is expected to switch randomly between gap and non-gap states between two ALS acquisitions (e.g., due to instrument differences or movement of leaves) and thus introduce a baseline bias in disturbance and recovery rates. The five panels below show a sample canopy height model at Harvard Forest (150×150 m) over a 12-year period. White contours indicate gaps or parts of gaps that formed between 2012 and 2014 and rapidly closed in the years afterwards. These would go undetected in a direct comparison between 2012 and 2024. The panels show that initial gap recovery stems predominantly from lateral ingrowth around gaps, and within-gap regeneration appears only in 2024 as lighter blue areas within gaps.

Our results show a moderately dynamic canopy at Harvard Forest, with close to 1% of the area (1 ha km^-2^) either disturbed or recovering year-to-year, and more than 20% of the gap area created by branch or treefalls closing again over 12 years (Fig. 4, bottom panels). This transient turnover, which we cannot observe when ALS acquisitions are repeated infrequently, is expected to be even larger in forest canopies with higher disturbance and recovery rates (Fig. S3.1). To quantify it, we need medium-to-high frequency monitoring programs (e.g., Cushman et al., 2022) and robust methods that can account for scanning noise. We found that nearly 2% of the pixels were expected to switch from canopy to gap status (∼1%) or vice versa (∼1%) simply due to measurement noise between ALS acquisitions. This was more than twice the mean yearly biological changes at Harvard Forest, and if not accounted for, would greatly inflate estimates of canopy dynamics. Single pairs of acquisitions may be particularly affected by these issues (Heiskanen et al., 2024), which can be further compounded by ecological variability, changes in acquisition timing (cf. “winter” outliers in Fig. S3.3a) or instable CHM algorithms (Fischer et al., 2024). Some instrumentation noise may be avoided by careful re-calibration (Riofrío et al., 2022) or by subsetting to larger or more severe disturbances (Fig. S3.2). However, the former is challenging, and the latter does not make biological sense when tracking recovery which is more gradual in nature (Körner, 2003). A more robust approach will consist in the pooling of information across multiple sites and acquisitions per site, ideally via Bayesian hierarchical models (Kohyama et al., 2018). Long-term ecological observatories such as NEON and harmonised collections like the GCA will provide an ideal backbone for such an approach and enable us to sketch a global picture of canopy turnover.

## 5. Outlook

### 5.1 In Humboldt’s and Bonpland’s tailwinds

Ever since Alexander von Humboldt and Aimé Bonpland set out to travel the Americas, comparing ecosystems and communities across large geographic gradients has been a cornerstone of modern Earth system science. On the one hand, ecosystems display a large degree of local idiosyncrasy, shaped as much by their unique geological and climatic past as the accidents of natural and human history. On the other hand, there are commonalities in how ecosystems develop that cut across biomes, continents and civilizations. These are encoded in vegetation structural patterns, such as how height and cover depend on water availability (Staver et al., 2011; Tao et al., 2016), how growing season length limits the elevation range of trees (Paulsen & Körner, 2014), and the successional mosaics left behind by natural and anthropogenic disturbance (Frelich et al., 2018; Jucker, 2022). Describing these patterns, identifying new ones, and linking them to the biophysical processes that underpin them not only enhances our understanding of the natural world, but also improves our ability to predict ecosystem responses to global change in decades to come (Blois et al., 2013). Remote sensing technologies such as ALS are uniquely poised to carry this project forward (Senf, 2022). Much of global ecology so far has been informed by a combination of local field samples and optical satellite layers – providing a “flat” view of the world as a tapestry of landcovers, punctuated by occasional data-rich information spikes grounded in field observations. ALS inserts 3D contours into this 2D world, converting discrete vegetation classes (e.g., forest and non-forest) into continuous vegetation metrics (e.g., vegetation height, cover and density). In doing so it helps us build a fuller and more nuanced picture of woody ecosystem structure and how it reflects current as well as past dynamics (Lenoir et al., 2022).

As we showed here, ALS can be a particularly powerful macroecological tool when acquisitions from different sources, sites and times are harmonized in large databases such as the GCA. High-resolution derived products such as CHMs and DTMs are comparatively small in size, robust, and easily analysed and interpreted with a wide range of software (Fischer et al., 2024). These detailed but accessible ecosystem maps shift the focus from resource-intensive data processing to ecological analysis, model building and technological innovation (White et al., 2019). As highlighted by our case studies, it requires comparatively little effort to use 3D maps to validate predictions from satellite-based sensors (case study 1). carry out robust global comparisons of complex structural features (case study 2) and precisely quantify ecosystem dynamics through time (case study 3). Naturally, our case studies offer only a glimpse into the opportunities provided by regional to global ALS harmonization. Comparisons of ALS-derived maps across and within biomes may fundamentally shift our conceptualization of ecosystem dynamics (Lines et al., 2022) and usher in a new generation of ecosystem models (Shugart et al., 2015). Future versions of the GCA and other similar projects could provide harmonised ALS collections covering even larger spatial extents, such as wall-to-wall maps of entire countries or continents (Moudrý et al., 2024). They could also be used to derive additional information from point clouds that goes beyond rasterized maps of height, such as leaf area densities (Vincent et al., 2023) and subcanopy structure (Jarron et al., 2020), further enhancing our picture of the 3D structure of the world’s terrestrial ecosystems.

### 5.2 Flying close to the ground

The new picture of woody ecosystem structure will take time to develop. When Humboldt and Bonpland arrived in Venezuela, they were so enthralled by the flora and fauna that they ran from one organism to the next like “madmen” (Humboldt, 1801). It then took them decades to systematize their knowledge. The characterisation of ecosystem structure from remote sensing and ALS is likely in a similar situation. Laser scanning has come a long way since its first applications in forestry, which date back to the 1970s and 1980s (Nelson, 2013). We now possess vast amounts of data and a wide range of tools to analyse them (Coops et al., 2021; Næsset, 2002; Roussel et al., 2020), but, in our characterization of global ecosystems, we are likely still in the initial exploration phase and will require years of consolidation to arrive at robust categorizations and predictions (Atkins et al., 2018; Fassnacht et al., 2024; Knapp et al., 2020; Loke & Chisholm, 2022; Rosen et al., 2024; Valbuena et al., 2020). The GCA provides a strong foundation for this enterprise. Its ecosystem maps provide samples from a wide range of environments (Wulder, White, Nelson, et al., 2012), have been developed to be maximally robust to differences between acquisitions (Fischer et al., 2024), and are accompanied by extensive metadata, quality layers and masks that can be used to identify areas where robustness may be compromised (Table S2).

Automated data processing and quality checks have their limitations, and for certain applications (e.g., measurements of within-canopy structure) additional corrections or data filtering may be needed, including targeted ground corrections (Riofrío et al., 2022), manual re-classification, or adjustments to account for instrument properties (Roussel et al., 2017; Vincent et al., 2023). Furthermore, as our case studies show, the need for careful analysis goes beyond these initial data selection and processing steps and extends to ecological model building itself. Just because we can predict canopy height from ALS-trained satellite data (case study 1), fit power laws to gap sizes (case study 2) or calculate structural change between successive acquisitions (case study 3) does not mean the results are always reliable or useful. Robust global ecosystem ecology needs to be guided by a biophysical understanding of the underlying dynamics, a careful selection of metrics to monitor them (Fischer et al., 2024; Zhang et al., 2024), models that can account and correct for biases, and standardized methods to assess their predictive power and transferability (Ploton et al., 2020; Tompalski et al., 2019). A key challenge going forward is to link ALS observations back to field data and fundamental ecological units such as trees and species (White et al., 2016). This will be facilitated by a wide range of methods, such as manual and automatic tree segmentation (Aubry-Kientz et al., 2021; Weinstein et al., 2021), model inversion (Shugart et al., 2015), or algorithms that attempt to reconstruct inventories bottom-up (Fischer et al., 2020; Spriggs et al., 2015; Taubert et al., 2015)

### 5.3 Flying together

Science is a social enterprise, and global databases such as the GCA are community efforts made possible by the collective investments, work and ideas of researchers across nations, generations and disciplines. While this creates a unique opportunity for collaborative research and data sharing, it also brings a responsibility to promote fair use. Access to and utilization of data are shaped by country-specific procurement models and regulatory openness as well as historical power asymmetries, such as those that often divide the Global South and Global North (de Angeli Dutra et al., 2025) or countries in and outside the tropics (de Lima et al., 2022). Such asymmetries are a particular concern with regard to remote sensing technologies such as ALS, which rely on expensive instruments and can be employed by domestic or foreign actors without meaningful involvement of local communities or researchers. The open release of such data is key as it creates opportunities for local experts and a wider public to actively contribute to research. Equally important are clear licensing and documentation that acknowledge local data contributors and institutions, ongoing support for acquisitions in undersampled areas and investments in capacity building. Care is also required to avoid misuse of data. Easily accessible remote sensing data could be used for the surveillance of communities (Millner et al., 2024) or to reveal the location of environmental assets (e.g. logging and mining opportunities) and archaeological sites (Fisher et al., 2021), potentially exacerbating disparities where access and safeguards are uneven. Moreover, external actors may violate data-ownership laws or license terms (e.g. non-commercial clauses), or repackage publicly available data into commercial products without proper attribution.

The GCA project cannot solve these problems by itself, but it can take steps to mitigate risks. First, by provisioning standardized ALS-derived products, but not the raw data, the GCA provides added value, but does not supplant the original data sources, which are thoroughly documented and linked to (cf. Table S1). Second, with its focus on simple, light-weight raster products, the GCA reduces the need for expensive computing infrastructure to process and analyse data. Third, it brings visibility to acquisitions missing from large open data repositories, which we hope will increase collaboration opportunities for local experts. Finally, the GCA has a relatively equal distribution of data in space, with representative coverage across the tropics (Fig. 1), ensuring that new products – often trained and tested in high-latitude temperate and boreal zones – become more representative of ecosystems and research needs globally. On balance, we believe that the benefits of making the majority of the GCA’s products openly available outweigh the risks. Most raw data in the GCA are already open, so the publication of harmonised, easy-to-use derived products should mostly benefit local researchers and practitioners who do not have the means to find, store and process large volumes of raw data. Past experience has shown that pre-processed openly available global datasets, such as meteorological and climate layers (Fick & Hijmans, 2017) and global trait compilations (Zanne et al., 2009), can greatly benefit researchers world-wide and fill gaps in areas where specialized local products and data are not as readily accessible.

To ensure the GCA promotes scientific discovery in a fair and equitable manner, we aim to implement a number of measures to further widen participation in the maintenance and governance of the database. This includes appointing an advisory committee to guide how best to continue growing the GCA in a way that does not unintentionally reinforce historical power imbalances. Similarly, we aim to invite local experts to join the project to help assemble ancillary datasets for the GCA landscapes so that we can build a richer picture of their ecology and history. Further, we plan to organise targeted online and in-person training courses designed to lower the barrier to entry for working with ALS data and facilitate their integration into existing research programs underway across GCA landscapes. Above all, we hope that we can encourage users of the database, particularly in the Global North, to contribute to these goals by involving local experts in their studies and investing in the maintenance and growth of the GCA network.

## Acknowledgements

Databases such as the GCA would not be possible without the work and investments of countless scientific and non-scientific actors that support data collection efforts in the field as well as the checking, analysis, and maintainance of data. We deeply thank all of them for their dedication and support.

FJ Fischer and T Jucker acknowledge a Research Project Grant from the Leverhulme Trust to T Jucker (grant: RPG-2020-341). FJ Fischer and R Seidl acknowledge the European Research Council under the European Union’s Horizon 2020 research and innovation program (Grant Agreement 101001905, FORWARD). T Jucker acknowledges a UK NERC Independent Research Fellowship (grant: NE/S01537X/1) and UKRI Frontier Research grant (grant: EP/Y003810/1). J Chave and N Labrière acknowledge ESA FRM4BIOMASS and Labex CEBA (ANR-10-LABX-25-01). KC Cushman and M Krassovski were supported by the Laboratory Directed Research and Development Program of Oak Ridge National Laboratory (ORNL); ORNL is managed by the University of Tennessee-Battelle, LLC, under contract DE-AC05-00OR22725 with the U.S. Department of Energy. K Stereńczak acknowledges Project LIFE+ ForBioSensing (contract number LIFE13ENV/PL/000048) and Poland’s National Fund for Environmental Protection and Water Management (contract number 485/2014/WN10/OPNMLF/D) and project “AFTER FBS – maintenance of ForBioSensing project performance indicators,” which is funded by the Forest Research Institute’s own research fund (no. 261509). J Aguirre-Gutiérrez acknowledges the Natural Environment Research Council (NERC; NE/T011084/1 and NE/Z504191/1), the Leverhulme Trust (RPG-2024-342) and the Royal Society (RG/R1/251370). A Burt and M Demol acknowledge support for data acquisitions in Gilé NP from a. World Bank through the Forest Carbon Partnership Facility b. Government of Mozambique through the Monitoring, Reporting and Verification Unit of the National Fund for Sustainable Development and c. Innovate UK (project number 10004871). J Cervenka acknowledges support from cross-border cooperation programme Czech Republic–Bavaria Free State ETC goal 2014–2020, the Interreg V project No. 99. F Fassnacht, J Schäfer, H Weiser and B Höfle acknowledge Deutsche Forschungsgemeinschaft (DFG, German Research Foundation) Project SYSSIFOSS - 411263134/2019-2022. L Fatoyinbo was supported by NASA Land Cover Land Use Change and Carbon Monitoring System Program. Eric Bastos Görgens was supported by CNPq grants 403297/2016-8, 401053/2019-9, 306386-2022-4. Amazon Fund grant 14.2.0929.1. USAID AID-OAA-A-11–00012. Acquisitions of airborne LiDAR data, contributed by T Gorum, were funded by DELTA Lidar, Türkiye. H Guan acknowledges the National Natural Science Foundation of China (32401574). Acquisition of airborne LiDAR data acquisition, contributed by A Hooijer and R Vernimmen, was funded by the Australian Agency for International Development and Dutch government (EMRP-BlockE, 2011), Norwegian Agency for Development Cooperation (Kampar, 2014) and United Kingdom Climate Change Unit (Berbak). K Král and M Krůček acknowledge INTER-COST project LUC23023. KK Htoo, R Farhadur, R Takeshige and Y Onoda acknowledge JSPS KAKENHI (#21H05314 & #21H02564). D Lai and OA Malik acknowledge Research Grant - Universiti Brunei Darussalam - UBD/RSCH/1.18/FICBF(b)/ 2023/006. D Lemke acknowledges The Nature Conservancy for access to the Sharp Bingham Mountain Preserve. J Lenoir acknowledges The Agence Nationale de la Recherche (ANR), under the framework of the young investigators’ funding scheme (JCJC Grant N°ANR-19-CE32-0005-01: IMPRINT project). Iain McNicol and E Mitchard acknowledge The European Union for funding ERC grant FODEX (757526). M Milenkovic acknowledges funding from the European Union’s Horizon Europe research and innovation programme under grant agreement No. 101059548. H Muller-Landau acknowledges Smithsonian ForestGEO (2023 Panama lidar data). JP Ometto acknowledges the Amazon Fund/BNDES (grant 14.2.0929.1, Environmental Monitoring of Brazilian Biomes), the US Agency for International Development, Grant/ Award Number: AID-OAA-A-11-00012, as well as the support of FAPESP grant 2017/22269-2. P Ploton acknowledges the support of the One Forest Vision initiative, funded by the French Ministry of Higher Education and Scientific Research and the French Ministry of Europe and Foreign Affairs. S Prober acknowledges support from the Western Australian government through its State NRM program and through Ngadju Conservation Aboriginal Corporation for the Great Western Woodlands drone-based LiDAR data. Airborne LiDAR for the Great Western Woodlands was funded through the Western Australian government through its Landgate program with further support from the Western Australian Department of Biodiversity, Conservation and Attractions and Australia’s Terrestrial Ecosystems Research Network. P Rana acknowledge the support of the governments of Finland and Nepal for the ForestResource Assessment (FRA) project. C Senf acknowledges the AI4Forest project (grant no. 01IS23025A) funded through the Bundesministerium für Bildung und Forschung (BMBF). A Shapiro acknowledges the support of the International Climate Initiative (IKI) of the German Federal Ministry for the Environment, Nature Conservation, Building and Nuclear Safety, and the German Development Bank KfW. G Shen acknowledges support by Natural Science Foundation of Shanghai (NSFS) (23ZR1419200), National Key R&D Program of China (NKRDP) (2024YFF1308100). M Silman acknowledges USAID/Wake Forest University Cooperative Agreement No. 7205-2721-CA-00005, NASA NISAR CAL/VAL, and the Andes-Amazon Fund. G Vincent acknowledges funding for airborne lidar data acquistions by the French Government through various agencies and research bodies and by UK NERC in 2019. WS Wan Mohd Jaafar acknowledges Ministry of Higher Education of Malaysia (Grant no: FRGS/1/2020/WAB03/UKM/02/1). J White acknowledges the Canadian Forest Service and the Ontario Ministry of Natural Resources. Z Yuan acknowledges the National Key Research and Development Program of China (2024YFF1306501). Y Zeng acknowledges the National Key R&D Program of China (2024YFF0810500).

## Author contributions

T Jucker conceived of the idea of the project. FJ Fischer and T Jucker led the data assembly and construction of the database, with assistance from J Chave, D Coomes, KC Cushman, R Dalagnol, M Dalponte, L Duncanson, S Saatchi, R Seidl, K Stereńczak and G Vaglio Laurin. FJ Fischer led the development of processing workflows and data processing, with assistance from T Jucker, B Morgan and T Jackson. FJ Fischer led the analysis and writing of a first draft of the manuscript, with assistance from T Jucker, T Jackson and B Morgan. All co-authors contributed substantially through data collection, data assembly and critical revisions of the manuscript.

## Data availability

All openly available GCA products (87% of the database, covering 95% of the area) will be made freely available after review through the European Space Agency’s Multi-Mission Algorithm and Analysis Platform (MAAP). All code and materials necessary to reproduce the analyses in this manuscript will be made publicly available on Zenodo (https://doi.org/10.5281/zenodo.16987211).

## References

1. Almeida, D. R. A. de, Stark, S. C., Shao, G., Schietti, J., Nelson, B. W., Silva, C. A., Gorgens, E. B., Valbuena, R., Papa, D. de A., & Brancalion, P. H. S. (2019). Optimizing the Remote Detection of Tropical Rainforest Structure with Airborne Lidar: Leaf Area Profile Sensitivity to Pulse Density and Spatial Sampling. Remote Sensing, 11(1). 10.3390/rs11010092

2. Araujo, R. F., Grubinger, S., Celes, C. H. S., Negrón-Juárez, R. I., Garcia, M., Dandois, J. P., & Muller-Landau, H. C. (2021). Strong temporal variation in treefall and branchfall rates in a tropical forest is related to extreme rainfall: results from 5 years of monthly drone data for a 50 ha plot. Biogeosciences, 18(24), 6517–6531. 10.5194/bg-18-6517-2021

3. Asner, G. P., Kellner, J. R., Kennedy-Bowdoin, T., Knapp, D. E., Anderson, C., & Martin, R. E. (2013). Forest Canopy Gap Distributions in the Southern Peruvian Amazon. PLOS ONE, 8(4), e60875-. 10.1371/journal.pone.0060875

4. Asner, G. P., Knapp, D. E., Martin, R. E., Tupayachi, R., Anderson, C. B., Mascaro, J., Sinca, F., Chadwick, K. D., Higgins, M., Farfan, W., Llactayo, W., & Silman, M. R. (2014). Targeted carbon conservation at national scales with high-resolution monitoring. Proceedings of the National Academy of Sciences, 111(47), E5016–E5022. 10.1073/pnas.1419550111

5. Atkins, J. W., Bohrer, G., Fahey, R. T., Hardiman, B. S., Morin, T. H., Stovall, A. E. L., Zimmerman, N., & Gough, C. M. (2018). Quantifying vegetation and canopy structural complexity from terrestrial LiDAR data using the forestr r package. Methods in Ecology and Evolution, 9(10), 2057–2066. 10.1111/2041-210X.13061

6. Atkins, J. W., Costanza, J., Dahlin, K. M., Dannenberg, M. P., Elmore, A. J., Fitzpatrick, M. C., Hakkenberg, C. R., Hardiman, B. S., Kamoske, A., LaRue, E. A., Silva, C. A., Stovall, A. E. L., & Tielens, E. K. (2023). Scale dependency of lidar-derived forest structural diversity. Methods in Ecology and Evolution, 14(2), 708–723. 10.1111/2041-210X.14040

7. Aubry-Kientz, M., Laybros, A., Weinstein, B., Ball, J. G. C., Jackson, T., Coomes, D., & Vincent, G. (2021). Multisensor Data Fusion for Improved Segmentation of Individual Tree Crowns in Dense Tropical Forests. IEEE Journal of Selected Topics in Applied Earth Observations and Remote Sensing, 14, 3927–3936. 10.1109/JSTARS.2021.3069159

8. Battison, R., Prober, S. M., Zdunic, K., Jackson, T. D., Fischer, F. J., & Jucker, T. (2024). Tracking tree demography and forest dynamics at scale using remote sensing. New Phytologist, 244(6), 2251–2266. 10.1111/nph.20199

9. Beland, M., Parker, G., Sparrow, B., Harding, D., Chasmer, L., Phinn, S., Antonarakis, A., & Strahler, A. (2019). On promoting the use of lidar systems in forest ecosystem research. Forest Ecology and Management, 450, 117484. 10.1016/j.foreco.2019.117484

10. Besic, N., Picard, N., Vega, C., Bontemps, J.-D., Hertzog, L., Renaud, J.-P., Fogel, F., Schwartz, M., Pellissier-Tanon, A., Destouet, G., Mortier, F., Planells-Rodriguez, M., & Ciais, P. (2025). Remote-sensing-based forest canopy height mapping: some models are useful, but might they provide us with even more insights when combined? Geoscientific Model Development, 18(2), 337–359. 10.5194/gmd-18-337-2025

11. Besson, M., Alison, J., Bjerge, K., Gorochowski, T. E., Høye, T. T., Jucker, T., Mann, H. M. R., & Clements, C. F. (2022). Towards the fully automated monitoring of ecological communities. Ecology Letters, 25(12), 2753–2775. 10.1111/ele.14123

12. Blois, J. L., Williams, J. W., Fitzpatrick, M. C., Jackson, S. T., & Ferrier, S. (2013). Space can substitute for time in predicting climate-change effects on biodiversity. Proceedings of the National Academy of Sciences, 110(23), 9374–9379. 10.1073/pnas.1220228110

13. Brandt, M., Tucker, C. J., Kariryaa, A., Rasmussen, K., Abel, C., Small, J., Chave, J., Rasmussen, L. V., Hiernaux, P., Diouf, A. A., Kergoat, L., Mertz, O., Igel, C., Gieseke, F., Schöning, J., Li, S., Melocik, K., Meyer, J., Sinno, S., … Fensholt, R. (2020). An unexpectedly large count of trees in the West African Sahara and Sahel. Nature, 587(7832), 78–82. 10.1038/s41586-020-2824-5

14. Bürkner, P. C. (2018). Advanced Bayesian multilevel modeling with the R package brms. R Journal, 10(1), 395–411. 10.32614/rj-2018-017

15. Cao, L., Coops, N. C., Innes, J. L., Sheppard, S. R. J., Fu, L., Ruan, H., & She, G. (2016). Estimation of forest biomass dynamics in subtropical forests using multi-temporal airborne LiDAR data. Remote Sensing of Environment, 178, 158–171. 10.1016/j.rse.2016.03.012

16. Coops, N. C., Tompalski, P., Goodbody, T. R. H., Queinnec, M., Luther, J. E., Bolton, D. K., White, J. C., Wulder, M. A., van Lier, O. R., & Hermosilla, T. (2021). Modelling lidar-derived estimates of forest attributes over space and time: A review of approaches and future trends. Remote Sensing of Environment, 260, 112477. 10.1016/j.rse.2021.112477

17. Cushman, K. C., Detto, M., García, M., & Muller-Landau, H. C. (2022). Soils and topography control natural disturbance rates and thereby forest structure in a lowland tropical landscape. Ecology Letters, 25(5), 1126–1138. 10.1111/ele.13978

18. Dalagnol, R., Wagner, F. H., Galvão, L. S., Streher, A. S., Phillips, O. L., Gloor, E., Pugh, T. A. M., Ometto, J. P. H. B., & Aragão, L. E. O. C. (2021). Large-scale variations in the dynamics of Amazon forest canopy gaps from airborne lidar data and opportunities for tree mortality estimates. Scientific Reports, 11(1), 1388. 10.1038/s41598-020-80809-w

19. Davies, A. B., & Asner, G. P. (2014). Advances in animal ecology from 3D-LiDAR ecosystem mapping. Trends in Ecology & Evolution, 29(12), 681–691. 10.1016/j.tree.2014.10.005

20. de Angeli Dutra, D., Erikson, A., Genes, L., Dirzo, R., & Venturini, A. M. (2025). Elevating local perspectives for equity in ecological research. Trends in Ecology & Evolution, 40(5), 415–418. 10.1016/j.tree.2025.03.006

21. de Lima, R. A. F., Phillips, O. L., Duque, A., Tello, J. S., Davies, S. J., de Oliveira, A. A., Muller, S., Honorio Coronado, E. N., Vilanova, E., Cuni-Sanchez, A., Baker, T. R., Ryan, C. M., Malizia, A., Lewis, S. L., ter Steege, H., Ferreira, J., Marimon, B. S., Luu, H. T., Imani, G., … Vásquez, R. (2022). Making forest data fair and open. Nature Ecology & Evolution, 6(6), 656–658. 10.1038/s41559-022-01738-7

22. Denslow, J. S. (1987). Tropical Rainforest Gaps and Tree Species Diversity. Annual Review of Ecology and Systematics, 18, 431–451. http://www.jstor.org.bris.idm.oclc.org/stable/2097139

23. Didan, K. (2025, March 16). MODIS/Terra Vegetation Indices Monthly L3 Global 0.05Deg CMG V061 [Data set]. NASA EOSDIS Land Processes Distributed Active Archive Center.

24. Dinerstein, E., Olson, D., Joshi, A., Vynne, C., Burgess, N. D., Wikramanayake, E., Hahn, N., Palminteri, S., Hedao, P., Noss, R., Hansen, M., Locke, H., Ellis, E. C., Jones, B., Barber, C. V., Hayes, R., Kormos, C., Martin, V., Crist, E., … Saleem, M. (2017). An Ecoregion-Based Approach to Protecting Half the Terrestrial Realm. BioScience, 67(6), 534–545. 10.1093/biosci/bix014

25. Dowle, M., & Srinivasan, A. (2025). data.table: Extension of ‘data.framè. https://CRAN.R-project.org/package=data.table

26. Eitel, J. U. H., Höfle, B., Vierling, L. A., Abellán, A., Asner, G. P., Deems, J. S., Glennie, C. L., Joerg, P. C., LeWinter, A. L., Magney, T. S., Mandlburger, G., Morton, D. C., Müller, J., & Vierling, K. T. (2016). Beyond 3-D: The new spectrum of lidar applications for earth and ecological sciences. Remote Sensing of Environment, 186, 372–392. 10.1016/j.rse.2016.08.018

27. European Space Agency. (2024). Copernicus Global Digital Elevation Model. Distributed by OpenTopography.

28. Fassnacht, F. E., White, J. C., Wulder, M. A., & Næsset, E. (2024). Remote sensing in forestry: current challenges, considerations and directions. Forestry: An International Journal of Forest Research, 97(1), 11–37. 10.1093/forestry/cpad024

29. Fick, S. E., & Hijmans, R. J. (2017). WorldClim 2: new 1-km spatial resolution climate surfaces for global land areas. International Journal of Climatology, 37(12), 4302–4315. 10.1002/joc.5086

30. Fischer, F. J., Jackson, T., Vincent, G., & Jucker, T. (2024). Robust characterisation of forest structure from airborne laser scanning—A systematic assessment and sample workflow for ecologists. Methods in Ecology and Evolution, 15(10), 1873–1888. 10.1111/2041-210X.14416

31. Fischer, F. J., & Jucker, T. (2024). No evidence for fractal scaling in canopy surfaces across a diverse range of forest types. Journal of Ecology, 112(3), 470–486. 10.1111/1365-2745.14244

32. Fischer, F. J., Labrière, N., Vincent, G., Hérault, B., Alonso, A., Memiaghe, H., Bissiengou, P., Kenfack, D., Saatchi, S., & Chave, J. (2020). A simulation method to infer tree allometry and forest structure from airborne laser scanning and forest inventories. Remote Sensing of Environment, 251, 112056. 10.1016/j.rse.2020.112056

33. Franklin, J. F., Shugart, H. H., & Harmon, M. E. (1987). Tree death as an ecological process. BioScience, 37(8), 550–556. 10.2307/1310665

34. Frelich, L. E., Jõgiste, K., Stanturf, J. A., Parro, K., & Baders, E. (2018). Natural Disturbances and Forest Management: Interacting Patterns on the Landscape. In A. H. Perera, U. Peterson, G. M. Pastur, & L. R. Iverson (Eds.), Ecosystem Services from Forest Landscapes: Broadscale Considerations (pp. 221–248). Springer International Publishing. 10.1007/978-3-319-74515-2_8

35. Gillespie, C. S. (2015). Fitting Heavy Tailed Distributions: The poweRlaw Package. Journal of Statistical Software, 64(2), 1–16. http://www.jstatsoft.org/v64/i02/

36. Gonzalez, A., Vihervaara, P., Balvanera, P., Bates, A. E., Bayraktarov, E., Bellingham, P. J., Bruder, A., Campbell, J., Catchen, M. D., Cavender-Bares, J., Chase, J., Coops, N., Costello, M. J., Czúcz, B., Delavaud, A., Dornelas, M., Dubois, G., Duffy, E. J., Eggermont, H., … Torrelio, C. Z. (2023). A global biodiversity observing system to unite monitoring and guide action. Nature Ecology & Evolution, 7(12), 1947–1952. 10.1038/s41559-023-02171-0

37. Goodbody, T. R. H., Tompalski, P., Coops, N. C., White, J. C., Wulder, M. A., & Sanelli, M. (2020). Uncovering spatial and ecological variability in gap size frequency distributions in the Canadian boreal forest. Scientific Reports, 10(1), 6069. 10.1038/s41598-020-62878-z

38. Gril, E., Laslier, M., Gallet-Moron, E., Durrieu, S., Spicher, F., Le Roux, V., Brasseur, B., Haesen, S., Van Meerbeek, K., Decocq, G., Marrec, R., & Lenoir, J. (2023). Using airborne LiDAR to map forest microclimate temperature buffering or amplification. Remote Sensing of Environment, 298, 113820. 10.1016/j.rse.2023.113820

39. Hanel, R., Corominas-Murtra, B., Liu, B., & Thurner, S. (2017). Fitting power-laws in empirical data with estimators that work for all exponents. PLOS ONE, 12(2), 1–15. 10.1371/journal.pone.0170920

40. Heiskanen, J., Haurinen, H., Wekesa, C., & Pellikka, P. (2024). Repeat airborne laser scanning to assess canopy height changes in tropical montane forests. ISPRS Ann. Photogramm. Remote Sens. Spatial Inf. Sci., X-3*–*2024, 179–185. 10.5194/isprs-annals-X-3-2024-179-2024

41. Hijmans, R. J. (2025). terra: Spatial Data Analysis. https://CRAN.R-project.org/package=terra

42. Huertas, C., Sabatier, D., Derroire, G., Ferry, B., Jackson, Toby. D., Pélissier, R., & Vincent, G. (2022). Mapping tree mortality rate in a tropical moist forest using multi-temporal LiDAR. International Journal of Applied Earth Observation and Geoinformation, 109, 102780. 10.1016/j.jag.2022.102780

43. Humboldt, A. von. (1801). Briefe des Herrn Alexander von Humboldt. Neue Berlinische Monatschrift, 115–141.

44. Hunter, M. O., Keller, M., Morton, D., Cook, B., Lefsky, M., Ducey, M., Saleska, S., de Oliveira Jr, R. C., & Schietti, J. (2015). Structural Dynamics of Tropical Moist Forest Gaps. PLOS ONE, 10(7), 1–19. 10.1371/journal.pone.0132144

45. Jackson, T. D., Fischer, F. J., Vincent, G., Gorgens, E. B., Keller, M., Chave, J., Jucker, T., & Coomes, D. A. (2024). Tall Bornean forests experience higher canopy disturbance rates than those in the eastern Amazon or Guiana shield. Global Change Biology, 30(9), e17493. 10.1111/gcb.17493

46. Jarron, L. R., Coops, N. C., MacKenzie, W. H., Tompalski, P., & Dykstra, P. (2020). Detection of sub-canopy forest structure using airborne LiDAR. Remote Sensing of Environment, 244, 111770. 10.1016/j.rse.2020.111770

47. Joerg, P. C., Morsdorf, F., & Zemp, M. (2012). Uncertainty assessment of multi-temporal airborne laser scanning data: A case study on an Alpine glacier. Remote Sensing of Environment, 127, 118–129. 10.1016/j.rse.2012.08.012

48. Jucker, T. (2022). Deciphering the fingerprint of disturbance on the three-dimensional structure of the world’s forests. New Phytologist, 233(2), 612–617. 10.1111/nph.17729

49. Jucker, T., Fischer, F. J., Chave, J., Coomes, D. A., Caspersen, J., Ali, A., Loubota Panzou, G. J., Feldpausch, T. R., Falster, D., Usoltsev, V. A., Adu-Bredu, S., Alves, L. F., Aminpour, M., Angoboy, I. B., Anten, N. P. R., Antin, C., Askari, Y., Muñoz, R., Ayyappan, N., … Zavala, M. A. (2022). Tallo: A global tree allometry and crown architecture database. Global Change Biology, 28(17). 10.1111/gcb.16302

50. Kampe, T. U. (2010). NEON: the first continental-scale ecological observatory with airborne remote sensing of vegetation canopy biochemistry and structure. Journal of Applied Remote Sensing, 4(1), 043510. 10.1117/1.3361375

51. Karan, M., Liddell, M., Prober, S. M., Arndt, S., Beringer, J., Boer, M., Cleverly, J., Eamus, D., Grace, P., Van Gorsel, E., Hero, J.-M., Hutley, L., Macfarlane, C., Metcalfe, D., Meyer, W., Pendall, E., Sebastian, A., & Wardlaw, T. (2016). The Australian SuperSite Network: A continental, long-term terrestrial ecosystem observatory. Science of The Total Environment, 568, 1263–1274. 10.1016/j.scitotenv.2016.05.170

52. Karger, D. N., Conrad, O., Böhner, J., Kawohl, T., Kreft, H., Soria-Auza, R. W., Zimmermann, N. E., Linder, H. P., & Kessler, M. (2017). Climatologies at high resolution for the earth’s land surface areas. Scientific Data, 4(1), 170122. 10.1038/sdata.2017.122

53. Kellner, J. R., & Asner, G. P. (2009). Convergent structural responses of tropical forests to diverse disturbance regimes. Ecology Letters, 12(9), 887–897. 10.1111/j.1461-0248.2009.01345.x

54. Kellogg, K., Hoffman, P., Standley, S., Shaffer, S., Rosen, P., Edelstein, W., Dunn, C., Baker, C., Barela, P., Shen, Y., Guerrero, A. M., Xaypraseuth, P., Sagi, V. R., Sreekantha, C. V, Harinath, N., Kumar, R., Bhan, R., & Sarma, C. V. H. S. (2020). NASA-ISRO Synthetic Aperture Radar (NISAR) Mission. 2020 IEEE Aerospace Conference, 1–21. 10.1109/AERO47225.2020.9172638

55. Khosravipour, A., Skidmore, A. K., & Isenburg, M. (2016). Generating spike-free digital surface models using LiDAR raw point clouds: A new approach for forestry applications. International Journal of Applied Earth Observation and Geoinformation, 52, 104–114. 10.1016/j.jag.2016.06.005

56. Kissling, W. D., & Shi, Y. (2023). Which metrics derived from airborne laser scanning are essential to measure the vertical profile of ecosystems? Diversity and Distributions, 29(10), 1315–1320. 10.1111/ddi.13760

57. Knapp, N., Fischer, R., Cazcarra-Bes, V., & Huth, A. (2020). Structure metrics to generalize biomass estimation from lidar across forest types from different continents. Remote Sensing of Environment, 237, 111597. 10.1016/j.rse.2019.111597

58. Kohyama, T. S., Kohyama, T. I., & Sheil, D. (2018). Definition and estimation of vital rates from repeated censuses: Choices, comparisons and bias corrections focusing on trees. Methods in Ecology and Evolution, 9(4), 809–821. 10.1111/2041-210X.12929

59. Kohyama, T. S., Kohyama, T. I., & Sheil, D. (2019). Estimating net biomass production and loss from repeated measurements of trees in forests and woodlands: Formulae, biases and recommendations. Forest Ecology and Management, 433, 729–740. 10.1016/j.foreco.2018.11.010

60. Körner, C. (2003). Slow in, Rapid out--Carbon Flux Studies and Kyoto Targets. Science, 300(5623), 1242–1243. 10.1126/science.1084460

61. Krüger, K., Senf, C., Jucker, T., Pflugmacher, D., & Seidl, R. (2024). Gap expansion is the dominant driver of canopy openings in a temperate mountain forest landscape. Journal of Ecology, 112(7), 1501–1515. 10.1111/1365-2745.14320

62. Lang, N., Jetz, W., Schindler, K., & Wegner, J. D. (2023). A high-resolution canopy height model of the Earth. Nature Ecology & Evolution, 7(11), 1778–1789. 10.1038/s41559-023-02206-6

63. LaRue, E. A., Fahey, R., Fuson, T. L., Foster, J. R., Matthes, J. H., Krause, K., & Hardiman, B. S. (2022). Evaluating the sensitivity of forest structural diversity characterization to LiDAR point density. Ecosphere, 13(9), e4209. 10.1002/ecs2.4209

64. Leitold, V., Morton, D. C., Longo, M., dos-Santos, M. N., Keller, M., & Scaranello, M. (2018). El Niño drought increased canopy turnover in Amazon forests. New Phytologist, 219(3), 959–971. 10.1111/nph.15110

65. Lenoir, J., Gril, E., Durrieu, S., Horen, H., Laslier, M., Lembrechts, J. J., Zellweger, F., Alleaume, S., Brasseur, B., Buridant, J., Dayal, K., De Frenne, P., Gallet-Moron, E., Marrec, R., Meeussen, C., Rocchini, D., Van Meerbeek, K., & Decocq, G. (2022). Unveil the unseen: Using LiDAR to capture time-lag dynamics in the herbaceous layer of European temperate forests. Journal of Ecology, 110(2), 282–300.

66. Lines, E. R., Fischer, F. J., Owen, H. J. F., & Jucker, T. (2022). The shape of trees: Reimagining forest ecology in three dimensions with remote sensing. Journal of Ecology, 110(8). 10.1111/1365-2745.13944

67. Liu, S., Brandt, M., Nord-Larsen, T., Chave, J., Reiner, F., Lang, N., Tong, X., Ciais, P., Igel, C., Pascual, A., Guerra-Hernandez, J., Li, S., Mugabowindekwe, M., Saatchi, S., Yue, Y., Chen, Z., & Fensholt, R. (2025). The overlooked contribution of trees outside forests to tree cover and woody biomass across Europe. Science Advances, 9(37), eadh4097. 10.1126/sciadv.adh4097

68. Lobo, E., & Dalling, J. W. (2014). Spatial scale and sampling resolution affect measures of gap disturbance in a lowland tropical forest: implications for understanding forest regeneration and carbon storage. Proceedings of the Royal Society B: Biological Sciences, 281(1778), 20133218. 10.1098/rspb.2013.3218

69. Loke, L. H. L., & Chisholm, R. A. (2022). Measuring habitat complexity and spatial heterogeneity in ecology. Ecology Letters, 25(10), 2269–2288. 10.1111/ele.14084

70. Melendy, L., Hagen, S. C., Sullivan, F. B., Pearson, T. R. H., Walker, S. M., Ellis, P., Kustiyo, Sambodo, A. K., Roswintiarti, O., Hanson, M. A., Klassen, A. W., Palace, M. W., Braswell, B. H., & Delgado, G. M. (2018). Automated method for measuring the extent of selective logging damage with airborne LiDAR data. ISPRS Journal of Photogrammetry and Remote Sensing, 139, 228–240. 10.1016/j.isprsjprs.2018.02.022

71. Moudrý, V., Remelgado, R., Forkel, M., Torresani, M., Laurin, G. V., Sarovcova, E., Millan, V. E. G., Fischer, F. J., Jucker, T., Gallay, M., Kacic, P., Hakkenberg, C. R., Kokalj, Ž., Stereńczak, K., Erfanifard, Y., Rocchini, D., Prošek, J., Roilo, S., Gdulova, K., … Barták, V. (2024). Harmonised airborne laser scanning products can address the limitations of large-scale spaceborne vegetation mapping [pre-print]. EarthArXiv. 10.31223/X5D70J

72. Mutanga, O., Masenyama, A., & Sibanda, M. (2023). Spectral saturation in the remote sensing of high-density vegetation traits: A systematic review of progress, challenges, and prospects. ISPRS Journal of Photogrammetry and Remote Sensing, 198, 297–309. 10.1016/j.isprsjprs.2023.03.010

73. Næsset, E. (2002). Predicting forest stand characteristics with airborne scanning laser using a practical two-stage procedure and field data. Remote Sensing of Environment, 80(1), 88–99. 10.1016/S0034-4257(01)00290-5

74. Næsset, E. (2005). Assessing sensor effects and effects of leaf-off and leaf-on canopy conditions on biophysical stand properties derived from small-footprint airborne laser data. Remote Sensing of Environment, 98(2), 356–370. 10.1016/j.rse.2005.07.012

75. Næsset, E. (2009). Effects of different sensors, flying altitudes, and pulse repetition frequencies on forest canopy metrics and biophysical stand properties derived from small-footprint airborne laser data. Remote Sensing of Environment, 113(1), 148–159. 10.1016/j.rse.2008.09.001

76. Næsset, E., Bollandsås, O. M., Gobakken, T., Gregoire, T. G., & Ståhl, G. (2013). Model-assisted estimation of change in forest biomass over an 11year period in a sample survey supported by airborne LiDAR: A case study with post-stratification to provide “activity data”. Remote Sensing of Environment, 128, 299–314. 10.1016/j.rse.2012.10.008

77. Næsset, E., Gobakken, T., Holmgren, J., Hyyppä, H., Hyyppä, J., Maltamo, M., Nilsson, M., Olsson, H., Persson, Å., & Söderman, U. (2004). Laser scanning of forest resources: the nordic experience. Scandinavian Journal of Forest Research, 19(6), 482–499. 10.1080/02827580410019553

78. Nelson, R. (2013). How did we get here? An early history of forestry lidar. Canadian Journal of Remote Sensing, 39, S6–S17. 10.5589/m13-011

79. Newman, M. E. J. (2005). Power laws, Pareto distributions and Zipf’s law. Contemporary Physics, 46(5), 323–351. 10.1080/00107510500052444

80. Okyay, U., Telling, J., Glennie, C. L., & Dietrich, W. E. (2019). Airborne lidar change detection: An overview of Earth sciences applications. Earth-Science Reviews, 198, 102929. 10.1016/j.earscirev.2019.102929

81. Ometto, J. P., Gorgens, E. B., de Souza Pereira, F. R., Sato, L., de Assis, M. L. R., Cantinho, R., Longo, M., Jacon, A. D., & Keller, M. (2023). A biomass map of the Brazilian Amazon from multisource remote sensing. Scientific Data, 10(1), 668. 10.1038/s41597-023-02575-4

82. Paulsen, J., & Körner, C. (2014). A climate-based model to predict potential treeline position around the globe. Alpine Botany, 124(1), 1–12. 10.1007/s00035-014-0124-0

83. Pebesma, E. (2018). Simple Features for R: Standardized Support for Spatial Vector Data. The R Journal, 10(1), 439–446. 10.32614/RJ-2018-009

84. Ploton, P., Mortier, F., Réjou-Méchain, M., Barbier, N., Picard, N., Rossi, V., Dormann, C., Cornu, G., Viennois, G., Bayol, N., Lyapustin, A., Gourlet-Fleury, S., & Pélissier, R. (2020). Spatial validation reveals poor predictive performance of large-scale ecological mapping models. Nature Communications, 11(1). 10.1038/s41467-020-18321-y

85. Potapov, P., Hansen, M. C., Laestadius, L., Turubanova, S., Yaroshenko, A., Thies, C., Smith, W., Zhuravleva, I., Komarova, A., Minnemeyer, S., & Esipova, E. (2017). The last frontiers of wilderness: Tracking loss of intact forest landscapes from 2000 to 2013. Science Advances, 3, e1600821. 10.1126/sciadv.1600821

86. Potapov, P., Hansen, M. C., Pickens, A., Hernandez-Serna, A., Tyukavina, A., Turubanova, S., Zalles, V., Li, X., Khan, A., Stolle, F., Harris, N., Song, X.-P., Baggett, A., Kommareddy, I., & Kommareddy, A. (2022). The Global 2000-2020 Land Cover and Land Use Change Dataset Derived From the Landsat Archive: First Results. Frontiers in Remote Sensing, 3. https://www.frontiersin.org/journals/remote-sensing/articles/10.3389/frsen.2022.856903

87. Potapov, P., Li, X., Hernandez-Serna, A., Tyukavina, A., Hansen, M. C., Kommareddy, A., Pickens, A., Turubanova, S., Tang, H., Silva, C. E., Armston, J., Dubayah, R., Blair, J. B., & Hofton, M. (2021). Mapping global forest canopy height through integration of GEDI and Landsat data. Remote Sensing of Environment, 253, 112165. 10.1016/j.rse.2020.112165

88. Quan, Y., Li, M., Hao, Y., & Wang, B. (2021). Comparison and Evaluation of Different Pit-Filling Methods for Generating High Resolution Canopy Height Model Using UAV Laser Scanning Data. Remote Sensing, 13(12). 10.3390/rs13122239

89. Quegan, S., Le Toan, T., Chave, J., Dall, J., Exbrayat, J. F., Minh, D. H. T., Lomas, M., D’Alessandro, M. M., Paillou, P., Papathanassiou, K., Rocca, F., Saatchi, S., Scipal, K., Shugart, H., Smallman, T. L., Soja, M. J., Tebaldini, S., Ulander, L., Villard, L., & Williams, M. (2019). The European Space Agency BIOMASS mission: Measuring forest above-ground biomass from space. Remote Sensing of Environment, 227, 44–60. 10.1016/j.rse.2019.03.032

90. Reis, C. R., Jackson, T. D., Gorgens, E. B., Dalagnol, R., Jucker, T., Nunes, M. H., Ometto, J. P., Aragão, L. E. O. C., Rodriguez, L. C. E., & Coomes, D. A. (2022). Forest disturbance and growth processes are reflected in the geographical distribution of large canopy gaps across the Brazilian Amazon. Journal of Ecology, 110(12), 2971–2983. 10.1111/1365-2745.14003

91. Riofrío, J., White, J. C., Tompalski, P., Coops, N. C., & Wulder, M. A. (2022). Harmonizing multi-temporal airborne laser scanning point clouds to derive periodic annual height increments in temperate mixedwood forests. Canadian Journal of Forest Research, 52(10), 1334–1352. 10.1139/cjfr-2022-0055

92. Ritter, E., Dalsgaard, L., & Einhorn, K. S. (2005). Light, temperature and soil moisture regimes following gap formation in a semi-natural beech-dominated forest in Denmark. Forest Ecology and Management, 206(1), 15–33. 10.1016/j.foreco.2004.08.011

93. Rosen, A., Jörg Fischer, F., Coomes, D. A., Jackson, T. D., Asner, G. P., & Jucker, T. (2024). Tracking shifts in forest structural complexity through space and time in human-modified tropical landscapes. *Ecography*, *n/a*(n/a), e07377. 10.1111/ecog.07377

94. Roussel, J.-R., Auty, D., Coops, N. C., Tompalski, P., Goodbody, T. R. H., Meador, A. S., Bourdon, J.-F., de Boissieu, F., & Achim, A. (2020). lidR: An R package for analysis of Airborne Laser Scanning (ALS) data. Remote Sensing of Environment, 251, 112061. 10.1016/j.rse.2020.112061

95. Roussel, J.-R., Caspersen, J., Béland, M., Thomas, S., & Achim, A. (2017). Removing bias from LiDAR-based estimates of canopy height: Accounting for the effects of pulse density and footprint size. Remote Sensing of Environment, 198, 1–16. 10.1016/j.rse.2017.05.032

96. Saatchi, S. S., & Favrichon, S. (2023). Global Vegetation Height Metrics from GEDI and ICESat2. ORNL DAAC. 10.3334/ORNLDAAC/2294

97. Schwalb-Willmann, J. (2024). *basemaps: Accessing Spatial Basemaps in R*. https://CRAN.R-project.org/package=basemaps

98. Schwartz, M., Ciais, P., De Truchis, A., Chave, J., Ottlé, C., Vega, C., Wigneron, J.-P., Nicolas, M., Jouaber, S., Liu, S., Brandt, M., & Fayad, I. (2023). FORMS: Forest Multiple Source height, wood volume, and biomass maps in France at 10 to 30m resolution based on Sentinel-1, Sentinel-2, and Global Ecosystem Dynamics Investigation (GEDI) data with a deep learning approach. Earth System Science Data, 15(11), 4927–4945. 10.5194/essd-15-4927-2023

99. Senf, C. (2022). Seeing the System from Above: The Use and Potential of Remote Sensing for Studying Ecosystem Dynamics. Ecosystems, 25(8), 1719–1737. 10.1007/s10021-022-00777-2

100. Shugart, H. H., Asner, G. P., Fischer, R., Huth, A., Knapp, N., Le Toan, T., & Shuman, J. K. (2015). Computer and remote-sensing infrastructure to enhance large-scale testing of individual-based forest models. Frontiers in Ecology and the Environment, 13(9), 503–511. 10.1890/140327

101. Shugart, H. H., Saatchi, S., & Hall, F. G. (2010). Importance of structure and its measurement in quantifying function of forest ecosystems. Journal of Geophysical Research: Biogeosciences, 115(G2), n/a-n/a. 10.1029/2009JG000993

102. Simard, M., Pinto, N., Fisher, J. B., & Baccini, A. (2011). Mapping forest canopy height globally with spaceborne lidar. Journal of Geophysical Research: Biogeosciences, 116(G4). 10.1029/2011JG001708

103. Spriggs, R. A., Vanderwel, M. C., Jones, T. A., Caspersen, J. P., & Coomes, D. A. (2015). A simple area-based model for predicting airborne LiDAR first returns from stem diameter distributions: An example study in an uneven-aged, mixed temperate forest. Canadian Journal of Forest Research, 45(10), 1338–1350. 10.1139/cjfr-2015-0018

104. Staver, A. C., Archibald, S., & Levin, S. A. (2011). The Global Extent and Determinants of Savanna and Forest as Alternative Biome States. Science, 334(6053), 230–232. 10.1126/science.1210465

105. Stereńczak, K., Ciesielski, M., Balazy, R., & Zawiła-Niedźwiecki, T. (2016). Comparison of various algorithms for DTM interpolation from LIDAR data in dense mountain forests. European Journal of Remote Sensing, 49(1), 599–621. 10.5721/EuJRS20164932

106. Stereńczak, K., Laurin, G. V., Chirici, G., Coomes, D. A., Dalponte, M., Latifi, H., & Puletti, N. (2020). Global Airborne Laser Scanning Data Providers Database (GlobALS)—A New Tool for Monitoring Ecosystems and Biodiversity. Remote Sensing, 12(11). 10.3390/rs12111877

107. Tao, S., Guo, Q., Li, C., Wang, Z., & Fang, J. (2016). Global patterns and determinants of forest canopy height. Ecology, 97(12), 3265–3270. 10.1002/ecy.1580

108. Taubert, F., Hartig, F., Dobner, H.-J., & Huth, A. (2013). On the Challenge of Fitting Tree Size Distributions in Ecology. PLOS ONE, 8(2), e58036-. 10.1371/journal.pone.0058036

109. Taubert, F., Jahn, M. W., Dobner, H.-J., Wiegand, T., & Huth, A. (2015). The structure of tropical forests and sphere packings. Proceedings of the National Academy of Sciences, 112(49), 15125–15129. 10.1073/pnas.1513417112

110. Toivonen, J., Kangas, A., Maltamo, M., Kukkonen, M., & Packalen, P. (2023). Assessing biodiversity using forest structure indicators based on airborne laser scanning data. Forest Ecology and Management, 546, 121376. 10.1016/j.foreco.2023.121376

111. Tolan, J., Yang, H.-I., Nosarzewski, B., Couairon, G., Vo, H. V, Brandt, J., Spore, J., Majumdar, S., Haziza, D., Vamaraju, J., Moutakanni, T., Bojanowski, P., Johns, T., White, B., Tiecke, T., & Couprie, C. (2024). Very high resolution canopy height maps from RGB imagery using self-supervised vision transformer and convolutional decoder trained on aerial lidar. Remote Sensing of Environment, 300, 113888. 10.1016/j.rse.2023.113888

112. Tompalski, P., White, J. C., Coops, N. C., & Wulder, M. A. (2019). Demonstrating the transferability of forest inventory attribute models derived using airborne laser scanning data. Remote Sensing of Environment, 227, 110–124. 10.1016/j.rse.2019.04.006

113. Valbuena, R., O’Connor, B., Zellweger, F., Simonson, W., Vihervaara, P., Maltamo, M., Silva, C. A., Almeida, D. R. A., Danks, F., Morsdorf, F., Chirici, G., Lucas, R., Coomes, D. A., & Coops, N. C. (2020). Standardizing Ecosystem Morphological Traits from 3D Information Sources. Trends in Ecology & Evolution, 35(8), 656–667. 10.1016/j.tree.2020.03.006

114. Vincent, G., Verley, P., Brede, B., Delaitre, G., Maurent, E., Ball, J., Clocher, I., & Barbier, N. (2023). Multi-sensor airborne lidar requires intercalibration for consistent estimation of light attenuation and plant area density. Remote Sensing of Environment, 286, 113442. 10.1016/j.rse.2022.113442

115. Wagner, F. H., Dalagnol, R., Carter, G., Hirye, M. C. M., Gill, S., Takougoum, L. B. S., Favrichon, S., Keller, M., Ometto, J. P. H. B., Alves, L., Creze, C., George-Chacon, S. P., Li, S., Liu, Z., Mullissa, A., Yang, Y., Santos, E. G., Worden, S. R., Brandt, M., … Saatchi, S. (2025). High Resolution Tree Height Mapping of the Amazon Forest using Planet NICFI Images and LiDAR-Informed U-Net Model. https://arxiv.org/abs/2501.10600

116. Wagner, F. H., Roberts, S., Ritz, A. L., Carter, G., Dalagnol, R., Favrichon, S., Hirye, M. C. M., Brandt, M., Ciais, P., & Saatchi, S. (2024). Sub-meter tree height mapping of California using aerial images and LiDAR-informed U-Net model. Remote Sensing of Environment, 305, 114099. 10.1016/j.rse.2024.114099

117. Wedeux, B. M. M., & Coomes, D. A. (2015). Landscape-scale changes in forest canopy structure across a partially logged tropical peat swamp. Biogeosciences, 12(22), 6707–6719. 10.5194/bg-12-6707-2015

118. Weinstein, B. G., Marconi, S., Bohlman, S. A., Zare, A., Singh, A., Graves, S. J., & White, E. P. (2021). A remote sensing derived data set of 100 million individual tree crowns for the national ecological observatory network. ELife, 10. 10.7554/eLife.62922

119. White, J. C., Chen, H., Woods, M. E., Low, B., & Nasonova, S. (2019). The Petawawa Research Forest: Establishment of a remote sensing supersite. The Forestry Chronicle, 95(03), 149–156. 10.5558/tfc2019-024

120. White, J. C., Coops, N. C., Wulder, M. A., Vastaranta, M., Hilker, T., & Tompalski, P. (2016). Remote Sensing Technologies for Enhancing Forest Inventories: A Review. In Canadian Journal of Remote Sensing (Vol. 42, Issue 5, pp. 619–641). 10.1080/07038992.2016.1207484

121. White, J. C., Tompalski, P., Bater, C. W., Wulder, M. A., Fortin, M., Hennigar, C., Robere-McGugan, G., Sinclair, I., & White, R. (2025). Enhanced forest inventories in Canada: implementation, status, and research needs. Canadian Journal of Forest Research, 55, 1–37. 10.1139/cjfr-2024-0255

122. White, J. C., Tompalski, P., Coops, N. C., & Wulder, M. A. (2018). Comparison of airborne laser scanning and digital stereo imagery for characterizing forest canopy gaps in coastal temperate rainforests. Remote Sensing of Environment, 208, 1–14. 10.1016/j.rse.2018.02.002

123. Wulder, M. A., White, J. C., Bater, C. W., Coops, N. C., Hopkinson, C., & Chen, G. (2012). Lidar plots — a new large-area data collection option: context, concepts, and case study. Canadian Journal of Remote Sensing, 38(5), 600–618. 10.5589/m12-049

124. Wulder, M. A., White, J. C., Nelson, R. F., Næsset, E., Ørka, H. O., Coops, N. C., Hilker, T., Bater, C. W., & Gobakken, T. (2012). Lidar sampling for large-area forest characterization: A review. Remote Sensing of Environment, 121, 196–209. 10.1016/j.rse.2012.02.001

125. Xu, L., Saatchi, S. S., Shapiro, A., Meyer, V., Ferraz, A., Yang, Y., Bastin, J. F., Banks, N., Boeckx, P., Verbeeck, H., Lewis, S. L., Muanza, E. T., Bongwele, E., Kayembe, F., Mbenza, D., Kalau, L., Mukendi, F., Ilunga, F., & Ebuta, D. (2017). Spatial Distribution of Carbon Stored in Forests of the Democratic Republic of Congo. Scientific Reports, 7(1), 15030. 10.1038/s41598-017-15050-z

126. Yu, X., Hyyppä, J., Kaartinen, H., & Maltamo, M. (2004). Automatic detection of harvested trees and determination of forest growth using airborne laser scanning. Remote Sensing of Environment, 90(4), 451–462. 10.1016/j.rse.2004.02.001

127. Zanne, A. E., Lopez-Gonzalez, G., Coomes, D. A., Ilic, J., Jansen, S., Lewis, S. L., Miller, R. B., Swenson, N. G., Wiemann, M. C., & Chave, J. (2009). Data from: Global wood density database. Dryad Digital Repository. 10.5061/dryad.234

128. Zhang, B., Fischer, F. J., Coomes, D. A., & Jucker, T. (2023). Logging leaves a fingerprint on the number, size, spatial configuration and geometry of tropical forest canopy gaps. Biotropica, 55(2), 354–367. 10.1111/btp.13190

129. Zhang, B., Fischer, F. J., Prober, S. M., Yeoh, P. B., Gosper, C. R., Zdunic, K., & Jucker, T. (2024). Robust retrieval of forest canopy structural attributes using multi-platform airborne LiDAR. Remote Sensing in Ecology and Conservation, n/a(n/a). 10.1002/rse2.398

