## Supplementary Materials for "The Global Canopy Atlas: analysis-ready maps of 3D structure for the world’s woody ecosystems"

### The Global Canopy Atlas – general supplementary

| dataset | citations | units | area (km²) | size (GB) | pd ($m^{-2}$) |
|---|---|---|---|---|---|
| **3DEP** | 3DEP. Map services and data available from U.S. Geological Survey, National Geospatial Program<br>https://rockyweb.usgs.gov/vdelivery/Datasets/Staged/Elevation/LPC/Projects/USGS_LPC_PR_PuertoRico_2015_LAS_2018<br>https://rockyweb.usgs.gov/vdelivery/Datasets/Staged/Elevation/LPC/Projects/PR_PuertoRico_VirginIslands_2018_D18/PR_PRVI_A_2018/<br>https://rockyweb.usgs.gov/vdelivery/Datasets/Staged/Elevation/LPC/Projects/USGS_LPC_PR_PRVI_F_2018<br>https://rockyweb.usgs.gov/vdelivery/Datasets/Staged/Elevation/LPC/Projects/CA_Eastern_San_Diego_Co_Lidar_2016_B16/CA_E_SanDiegoCo_2016/<br>https://rockyweb.usgs.gov/vdelivery/Datasets/Staged/Elevation/LPC/Projects/OR_McKenzieRiver_2021_B21/OR_McKenzieRiver_1_2021<br>https://rockyweb.usgs.gov/vdelivery/Datasets/Staged/Elevation/LPC/Projects/FL_Peninsular_FDEM_2018_D19_DRRA/FL_Peninsular_FDEM_Baker_2018/<br>https://rockyweb.usgs.gov/vdelivery/Datasets/Staged/Elevation/LPC/Projects/FL_Peninsular_FDEM_2018_D19_DRRA/FL_Peninsular_FDEM_Columbia_2018/<br>https://rockyweb.usgs.gov/vdelivery/Datasets/Staged/Elevation/LPC/Projects/GA_Statewide_2018_B18_DRRA/GA_Statewide_B3_2018<br>https://rockyweb.usgs.gov/vdelivery/Datasets/Staged/Elevation/LPC/Projects/CO_DRCOG_2020_B20/CO_DRCOG_3_2020<br>https://rockyweb.usgs.gov/vdelivery/Datasets/Staged/Elevation/LPC/Projects/CO_NorthwestCO_2020_D20/CO_NWCO_2_2020<br>https://rockyweb.usgs.gov/vdelivery/Datasets/Staged/Elevation/LPC/Projects/AK_Fairbanks_NSB_Lidar_2017_B17/AK_FairbanksNSB_QL2_2017<br>https://rockyweb.usgs.gov/vdelivery/Datasets/Staged/Elevation/LPC/Projects/NY_FEMAR2_Central_2018_D19/NY_FEMAR2_Central_4_2018<br>https://rockyweb.usgs.gov/vdelivery/Datasets/Staged/Elevation/LPC/Projects/NC_Phase5_2018_A18/NC_Phase5_Clay_2017<br>https://rockyweb.usgs.gov/vdelivery/Datasets/Staged/Elevation/LPC/Projects/NC_Phase5_2018_A18/NC_Phase5_Macon_2017<br>https://rockyweb.usgs.gov/vdelivery/Datasets/Staged/Elevation/LPC/Projects/legacy/SC_RICHLANDCO_2010<br>https://rockyweb.usgs.gov/vdelivery/Datasets/Staged/Elevation/LPC/Projects/Six_County_SC_LiDAR_Quality_Assurance/SC_6County_Calhoun_2012<br>https://rockyweb.usgs.gov/vdelivery/Datasets/Staged/Elevation/LPC/Projects/USGS_LPC_SC_East_Central_2017_LAS_2019<br>https://rockyweb.usgs.gov/vdelivery/Datasets/Staged/Elevation/LPC/Projects/SC_SavannahPeeDee_2019_B19/SC_SavannahPeeDee_5_2019<br>https://rockyweb.usgs.gov/vdelivery/Datasets/Staged/Elevation/LPC/Projects/Eastern_MN_State_LiDAR_Phases_3_4_and_5/MN_Arrowhead_B3_2011<br>https://rockyweb.usgs.gov/vdelivery/Datasets/Staged/Elevation/LPC/Projects/MN_RainyLake_2020_B20/MN_RainyLake_1_2020<br>https://rockyweb.usgs.gov/vdelivery/Datasets/Staged/Elevation/LPC/Projects/AZ_CochiseCounty_2020_B20/AZ_CochiseCounty_2_2020<br>https://rockyweb.usgs.gov/vdelivery/Datasets/Staged/Elevation/LPC/Projects/AZ_PimaCounty_2021_B21/AZ_PimaCo_2_2021<br>https://rockyweb.usgs.gov/vdelivery/Datasets/Staged/Elevation/LPC/Projects/AK_GlacierBay_2019_B19/AK_GlacierBay_4_2019<br>https://rockyweb.usgs.gov/vdelivery/Datasets/Staged/Elevation/LPC/Projects/UT_StatewideCenSouth_2020_A20/UT_StatewideCenSouth_1_2020<br>https://rockyweb.usgs.gov/vdelivery/Datasets/Staged/Elevation/LPC/Projects/legacy/ARRA_TN_SMOKYMTNS_2011<br>https://rockyweb.usgs.gov/vdelivery/Datasets/Staged/Elevation/LPC/Projects/TN_Eastern_TN_LiDAR_2016_B16/TN_Eastern_2_16_B16_Del2_2016<br>https://rockyweb.usgs.gov/vdelivery/Datasets/Staged/Elevation/LPC/Projects/CA_AZ_FEMA_R9_Lidar_2017_D18/CA_FEMA_Z4_B1_2018<br>https://rockyweb.usgs.gov/vdelivery/Datasets/Staged/Elevation/LPC/Projects/USGS_LPC_MA_NE_CMGP_Sandy_Z18_2013<br>https://rockyweb.usgs.gov/vdelivery/Datasets/Staged/Elevation/LPC/Projects/MI_Hiawatha_NF_2018_D18/MI_HiawathaNF_QL2_2018<br>https://rockyweb.usgs.gov/vdelivery/Datasets/Staged/Elevation/LPC/Projects/CA_NoCAL_3DEP_Supp_Funding_2018_D18/CA_NoCAL_Wildfires_B4_2018<br>https://rockyweb.usgs.gov/vdelivery/Datasets/Staged/Elevation/LPC/Projects/FL_Peninsular_2018_D18/FL_Peninsular_Marion_2018<br>https://rockyweb.usgs.gov/vdelivery/Datasets/Staged/Elevation/LPC/Projects/AK_Kenai_2008<br>https://rockyweb.usgs.gov/vdelivery/Datasets/Staged/Elevation/LPC/Projects/LA_Sabine_River_Lidar_2018_D18/LA_Sabine_River_Lidar_A1_2018<br>https://rockyweb.usgs.gov/vdelivery/Datasets/Staged/Elevation/LPC/Projects/LA_Sabine_River_Lidar_2018_D18/LA_Sabine_River_Lidar_A6_2018<br>https://rockyweb.usgs.gov/vdelivery/Datasets/Staged/Elevation/LPC/Projects/AR_Ouachita_FEMA_R6_Lidar_2016_D17/AR_Ouachita_B5_2016 | 84 | 4848.3 | 149.1 | 14.1 |

| | | | | | |
|---|---|---|---|---|---|
| | https://rockyweb.usgs.gov/vdelivery/Datasets/Staged/Elevation/LPC/Projects/CA_CarrHirzDeltaFires_2019_B19/CA_CarrHirzDeltaFires_2_2019<br>https://rockyweb.usgs.gov/vdelivery/Datasets/Staged/Elevation/LPC/Projects/AK_GlacierBay_2019_B19/AK_GlacierBay_5_2019<br>https://rockyweb.usgs.gov/vdelivery/Datasets/Staged/Elevation/LPC/Projects/AK_GlacierBay_2019_B19/AK_GlacierBay_B3_2019<br>https://rockyweb.usgs.gov/vdelivery/Datasets/Staged/Elevation/LPC/Projects/CO_NorthwestCO_2020_D20/CO_NWCO_1_2020<br>https://rockyweb.usgs.gov/vdelivery/Datasets/Staged/Elevation/LPC/Projects/CA_YosemiteNP_2019_D19/CA_YosemiteNP_2019<br>https://rockyweb.usgs.gov/vdelivery/Datasets/Staged/Elevation/LPC/Projects/USGS_LPC_MO_FEMAR7_North_A1_2017_LAS_2019<br>https://rockyweb.usgs.gov/vdelivery/Datasets/Staged/Elevation/LPC/Projects/FL_HurricaneMichael_2020_D20/FL_HurricaneMichael_3_2020<br>https://rockyweb.usgs.gov/vdelivery/Datasets/Staged/Elevation/LPC/Projects/CO_Southwest_NRCS_2018_D18/CO_Southwest_NRCS_B2_2018<br>https://rockyweb.usgs.gov/vdelivery/Datasets/Staged/Elevation/LPC/Projects/Olympic_Peninsula_WA_QL_1_LiDAR/WA_Olympic_Peninsula_2013<br>https://rockyweb.usgs.gov/vdelivery/Datasets/Staged/Elevation/LPC/Projects/USGS_LPC_WA_Olympic_Peninsula_C3_2017<br>https://rockyweb.usgs.gov/vdelivery/Datasets/Staged/Elevation/LPC/Projects/FL_Peninsular_2018_D18/FL_Peninsular_Putnam_2018<br>https://rockyweb.usgs.gov/vdelivery/Datasets/Staged/Elevation/LPC/Projects/AR_Ouachita_FEMA_R6_Lidar_2016_D17/AR_Ouachita_B6_2016<br>https://rockyweb.usgs.gov/vdelivery/Datasets/Staged/Elevation/LPC/Projects/WA_Olympic_Peninsula_Lidar_2017_B17/WA_Olympic_Peninsula_TL_2017<br>https://rockyweb.usgs.gov/vdelivery/Datasets/Staged/Elevation/LPC/Projects/USGS_LPC_WA_Olympic_Peninsula_C1_2017<br>https://rockyweb.usgs.gov/vdelivery/Datasets/Staged/Elevation/LPC/Projects/CA_SouthernSierra_2020_B20/CA_SouthernSierra_1_2020<br>https://rockyweb.usgs.gov/vdelivery/Datasets/Staged/Elevation/LPC/Projects/CA_SierraNevada_B22/CA_SierraNevada_14_B22<br>https://rockyweb.usgs.gov/vdelivery/Datasets/Staged/Elevation/LPC/Projects/WA_EasternCascades_2019_B19/WA_EasternCascades_6_2019<br>https://rockyweb.usgs.gov/vdelivery/Datasets/Staged/Elevation/LPC/Projects/CA_AZ_FEMA_R9_Lidar_2017_D18/CA_FEMA_Z4_B2_2018<br>https://rockyweb.usgs.gov/vdelivery/Datasets/Staged/Elevation/LPC/Projects/CO_Southwest_NRCS_2018_D18/CO_Southwest_NRCS_B3_2018<br>https://rockyweb.usgs.gov/vdelivery/Datasets/Staged/Elevation/LPC/Projects/WY_YellowstoneNP_2020_D20/WY_YellowstoneNP_3_2020<br>https://rockyweb.usgs.gov/vdelivery/Datasets/Staged/Elevation/LPC/Projects/HI_Hawaii_Island_Lidar_NOAA_2017_B17/HI_Hawaii_Island_2017 | | | | |
| **AHN** | Actueel Hoogtebestand Nederland. www.ahn.nl | 7 | 218.8 | 6.7 | 20 |
| **AMAPCameroon** | original data contributed to GCA | 2 | 43.8 | 1.8 | 364.8 |
| **AMAPCongo** | original data contributed to GCA | 1 | 10 | 0.3 | 109.6 |
| **AWIYakutia** | Kruse, Stefan; Jackisch, Robert; Heim, Birgit; Gloy, Josias; Herzschuh, Ulrike; Kolmogorov, Alexei; Zakharov, Evgenii S; Pestryakova, Luidmila A; Förster, Michael; Kleinschmit, Birgit (2023): Point clouds and high-level data products of 21 Forest success | 16 | 1.8 | 0.1 | 451.4 |
| **AfriSAR** | Fatoyinbo et al. 2021. The NASA AfriSAR campaign: Airborne SAR and lidar measurements of tropical forest structure and biomass in support of current and future space missions. RSE 264, 112533. https://doi.org/10.1016/j.rse.2021.112533 | 3 | 227.3 | 6.9 | 2.4 |
| **AgencijazaOkolje** | Agencija rs za okolje. http://gis.arso.gov.si/evode/profile.aspx?id=atlas_voda_Lidar@Arso&culture=en-US | 6 | 128.9 | 4.4 | 9.7 |
| **Amani** | Hansen et al. 2015. Modeling Aboveground Biomass in Dense Tropical Submontane Rainforest Using Airborne Laser Scanner Data. Remote Sensing, 788-807. https://doi.org/10.3390/rs70100788 | 1 | 131.8 | 4.7 | 27.5 |
| **AnkasaBiaBoin** | Vaglio Laurin et al. 2016. Above ground biomass and tree species richness estimation with airborne lidar in tropical Ghana forests. International Journal of Applied Earth Observation and Geoinformation, 371-379. https://doi.org/10.1016/j.jag.2016.07.008 | 4 | 114.4 | 4.5 | 10.2 |
| **BCOpen** | LidarBC. https://lidar.gov.bc.ca/ (downloaded from previous website: https://www2.gov.bc.ca/gov/content/data/geographic-data-services/lidarbc) | 8 | 239.7 | 8.1 | 15.9 |
| **BNmicroclim** | Becek et al. 2020. Brunei Darussalam rainforest temperature and light intensity data recorded in 2017. Data in Brief 33, 106425. | 1 | 0 | 0 | 17.5 |
| **BayernAtlas** | BayernAtlas. Laserpunkte. https://geodaten.bayern.de/opengeodata/OpenDataDetail.html?pn=laserdaten | 6 | 584.5 | 24.6 | 12.7 |
| **Berchtesgaden** | Mandl et al. 2023. Spaceborne LiDAR for characterizing forest structure across scales in the European Alps. Remote Sensing in Ecology and Conservation 9: 599-614 https://doi.org/10.1002/rse2.330 | 1 | 210.3 | 8.6 | 46 |

| | | | | | |
|---|---|---|---|---|---|
| **Bialowieza** | Sterenczak et al. 2020. Global Airborne Laser Scanning Data Providers Database (GlobALS) - A New Tool for Monitoring Ecosystems and Biodiversity. Remote Sens. 12, 1877. https://doi.org/10.3390/rs12111877 | 2 | 151.4 | 4.4 | 16.4 |
| **CFSLidarplots** | Wulder et al. 2012. Lidar plots - a new large-area data collection option: context, concepts, and case study. Can. J. Remote Sens., 600-618. https://doi.org/10.5589/m12-049 | 121 | 1527.2 | 62 | 1.4 |
| **CMSKalimantan** | Melendy et al. 2017. CMS: LiDAR-derived Canopy and Elevation for Sites in Kalimantan, Indonesia, 2014. ORNL DAAC, Oak Ridge, Tennessee, USA. https://doi.org/10.3334/ORNLDAAC/1540 | 92 | 1678.5 | 74.9 | 7.9 |
| **CMSZambezi** | Fatoyinbo & Trettin 2017. CMS: LiDAR Data for Mangrove Forests in the Zambezi River Delta, Mozambique, 2014. ORNL DAAC, Oak Ridge, Tennessee, USA. https://doi.org/10.3334/ORNLDAAC/1521 | 1 | 115 | 3.7 | 5.6 |
| **CZO** | Luquillo CZO Rio Blanco and Rio Mameyes LiDAR Survey 2010-2011. Distributed by OpenTopography. https://doi.org/10.5069/G9BZ63ZR<br>Southern Sierra Nevada Critical Zone Observatory: Snow Off. Distributed by OpenTopography. https://doi.org/10.5069/G9BP00QB | 17 | 583.6 | 19.7 | 7.6 |
| **Changbai** | original data contributed to GCA | 5 | 8 | 0.4 | 4.6 |
| **Choco** | Meyer et al. 2019. Forest degradation and biomass loss along the Choco region of Colombia. Carbon Balance Manage 14. https://doi.org/10.1186/s13021-019-0117-9 | 37 | 614.3 | 20.4 | 6.9 |
| **DEFRA** | National LIDAR Programme. https://www.data.gov.uk/dataset/f0db0249-f17b-4036-9e65-309148c97ce4/national-lidar-programme | 33 | 554.6 | 15.3 | 2.4 |
| **DWER** | DWER | 22 | 257 | 7.5 | 1.5 |
| **DavieslabCongo** | original data contributed to GCA | 3 | 16.7 | 0.7 | 245.9 |
| **DavieslabKenya** | original data contributed to GCA | 13 | 136.6 | 4.5 | 176.3 |
| **DavieslabMozambique** | original data contributed to GCA | 5 | 240.1 | 8.5 | 62.4 |
| **DavieslabSA** | original data contributed to GCA | 9 | 40.8 | 1.6 | 78 |
| **Dongsithouane** | Hou et al. 2011. Use of ALS, Airborne CIR and ALOS AVNIR-2 data for estimating tropical forest attributes in Lao PDR. ISPRS Journal of Photogrammetry and Remote Sensing 66, 776-786. https://doi.org/10.1016/j.isprsjprs.2011.09.005 | 1 | 393.7 | 13.5 | 0.8 |
| **EBA** | Ometto, Jean Pierre et al. 2023. L1A - Discrete airborne LiDAR transects collected by EBA in the Brazilian Amazon (Acre e Rondonia). https://doi.org/10.5281/zenodo.7689908<br>Ometto, Jean Pierre et al. 2023. L1A - Discrete airborne LiDAR transects collected by EBA in the Brazilian Amazon (Roraima e Amapa). https://doi.org/10.5281/zenodo.7689692<br>Ometto, Jean Pierre et al. 2023. L1A - Discrete airborne LiDAR transects collected by EBA in the Brazilian Amazon (Mato Grosso, Amazonas e Para). https://doi.org/10.5281/zenodo.7636453<br>Ometto, Jean Pierre et al. 2023. L1A - Discrete airborne LiDAR transects collected by EBA in the Brazilian Amazon (Maranhao e Tocantins). https://doi.org/10.5281/zenodo.7689209 | 878 | 7271.7 | 349.1 | 4.7 |
| **ELVIS Australia** | Elvis - Elevation and Depth - Foundation Spatial Data. https://elevation.fsdf.org.au/ | 99 | 3315.8 | 106.3 | 7.4 |
| **EODaSVlaanderen** | Vlaanderen EODAS. https://remotesensing.vlaanderen.be/apps/openlidar | 1 | 6.2 | 0.2 | 24 |
| **EarthShape** | Kuegler et al. 2022. 3D Point Clouds and Topographic Data from the Chilean Coastal Cordillera. V. 1.0. GFZ Data Services. https://doi.org/10.5880/fidgeo.2022.002 | 3 | 157 | 5.2 | 4.8 |
| **FG** | original data contributed to GCA | 7 | 112.5 | 4.1 | 32.3 |
| **FODEXGabon** | McNicol et al. 2021. To what extent can UAV photogrammetry replicate UAV LiDAR to determine forest structure? A test in two contrasting tropical forests. JGR: Biogeosciences 126, e2021JG006586. https://doi.org/10.1029/2021JG006586 | 1 | 5.6 | 0.3 | 332.1 |
| **FODEXPeru** | McNicol et al. 2021. To what extent can UAV photogrammetry replicate UAV LiDAR to determine forest structure? A test in two contrasting tropical forests. JGR: Biogeosciences 126, e2021JG006586. https://doi.org/10.1029/2021JG006586 | 2 | 9 | 0.4 | 343.7 |
| **G-LiHT-Bahamas** | Cook et al. 2013. NASA Goddard's Lidar, Hyperspectral and Thermal (G-LiHT) airborne imager. Remote Sensing 5:4045-4066, doi:10.3390/rs5084045. | 2 | 11.5 | 0.6 | 4.9 |
| **G-LiHT-Mexico2013** | Cook et al. 2013. NASA Goddard's Lidar, Hyperspectral and Thermal (G-LiHT) airborne imager. Remote Sensing 5:4045-4066, doi:10.3390/rs5084045. | 80 | 267.4 | 11.7 | 6 |
| **G-LiHT-PuertoRico** | Cook et al. 2013. NASA Goddard's Lidar, Hyperspectral and Thermal (G-LiHT) airborne imager. Remote Sensing 5:4045-4066, doi:10.3390/rs5084045. | 1 | 1.9 | 0.1 | 11.7 |
| **G-LiHT-US** | Cook et al. 2013. NASA Goddard's Lidar, Hyperspectral and Thermal (G-LiHT) airborne imager. Remote Sensing 5:4045-4066, doi:10.3390/rs5084045. | 88 | 644.1 | 24.6 | 8.1 |
| **GUGiK** | Dane pomiarowe LIDAR. https://www.geoportal.gov.pl/dane/dane-pomiarowe-lidar | 19 | 1148.6 | 36.6 | 12.3 |
| **GWWALS** | Jucker et al. 2023. Using multi-platform LiDAR to guide the conservation of the world's largest temperate woodland. Remote Sensing of Environment 296, 113745. | 37 | 2818.8 | 117.5 | 21.6 |

| | | | | | |
|---|---|---|---|---|---|
| **GWWDrone** | Jucker et al. 2023. Using multi-platform LiDAR to guide the conservation of the world's largest temperate woodland. Remote Sensing of Environment 296, 113745. | 131 | 14.3 | 0.7 | 49 |
| **GeoBasis-DE/LGB** | Laserscandaten. GeoBasis-DE / LGB. https://data.geobasis-bb.de/geobasis/daten/als/ | 1 | 7.3 | 0.3 | 12.8 |
| **GeodatenMV** | © GeoBasis-DE/M-V 2023; https://www.geoportal-mv.de/portal/Geowebdienste/Fachthemen/Hoehe_und_Gelaende | 1 | 40.3 | 1.5 | 18.4 |
| **GhanaUAV** | original data contributed to GCA | 9 | 0.2 | 0 | 143.8 |
| **GolaNP** | Kent, R., Lindsell, J. A., Vaglio Laurin, G., Valentini, R., & Coomes, D. A. (2015). Airborne LiDAR detects selectively logged tropical forest even in an advanced stage of recovery. Remote Sensing, 7(7), 8348-8367. | 27 | 173.9 | 7.3 | 9.3 |
| **Hainich** | Freistaat Thueringen. Download Offene Geodaten. https://geoportal.thueringen.de/gdi-th/download-offene-geodaten | 1 | 74 | 3 | 7.7 |
| **Heyelan** | Tolga Gorum 2019. Landslide recognition and mapping in a mixed forest environment from airborne LiDAR data. Engineering Geology, 105155. https://doi.org/10.1016/j.enggeo.2019.105155. | 1 | 62 | 2.2 | 11.6 |
| **HiWater** | Xiao & Wen 2014. HiWATER: Airborne LiDAR raw data in Hulugou catchment. A Big Earth Data Platform for Three Poles. DOI: 10.3972/hiwater.159.2014.db<br>Xiao & Wen 2014. HiWATER: Airborne LiDAR raw data in Qilian on Aug. 28, 2012. A Big Earth Data Platform for Three Poles. DOI: 10.3972/hiwater.159.2014.db<br>Xiao & Wen 2014. HiWATER: Airborne LiDAR raw data in Tianlaochi catchment. A Big Earth Data Platform for Three Poles. DOI: 10.3972/hiwater.159.2014.db | 3 | 80.9 | 2.8 | 2.8 |
| **HongKongCEDD** | Civil Engineering and Development Department of the Government of the Hong Kong Special Administrative Region; https://sdportal.cedd.gov.hk/ | 26 | 270 | 9.4 | 30.7 |
| **Hoydedata** | Hoydedata. https://hoydedata.no/LaserInnsyn2/ | 6 | 304.9 | 9.8 | 8.3 |
| **ICNF_aGIL** | Projeto aGIL - Dados LiDAR. ICNF. https://geocatalogo.icnf.pt/geovisualizador/agil/ | 1 | 123.7 | 4.6 | 13.9 |
| **IGNFrance** | LiDAR HD. https://geoservices.ign.fr/lidarhd | 24 | 270.9 | 9.4 | 29.7 |
| **IGNSpain** | PNOA-LiDAR. https://pnoa.ign.es/pnoa-lidar/presentacion | 41 | 3227.8 | 117.7 | 3 |
| **IMPRINT** | original data contributed to GCA | 3 | 152.1 | 5.2 | 76.3 |
| **IrkutskKomiPerm** | original data contributed to GCA | 15 | 24.1 | 0.6 | 22.3 |
| **JTSB** | original data contributed to GCA | 2 | 32.8 | 1.3 | 17.5 |
| **KhaoYai** | Jha et al. 2020. Forest aboveground biomass stock and resilience in a tropical landscape of Thailand. Biogeosciences 17, 121--134. DOI: 10.5194/bg-17-121-2020 | 1 | 47.3 | 1.9 | 25.9 |
| **KilimanjaroUFZ** | Getzin et al. 2017. Using airborne LiDAR to assess spatial heterogeneity in forest structure on Mount Kilimanjaro. Landscape Ecol 32, 1881-1894. https://doi.org/10.1007/s10980-017-0550-7 | 35 | 59.6 | 3.8 | 24.4 |
| **LINZ** | LINZ (2018). Abel Tasman and Golden Bay, Tasman, New Zealand 2016. Collected by AAM New Zealand Limited, distributed by OpenTopography and Land Information New Zealand (LINZ). https://doi.org/10.5069/G9BR8Q8X.<br>Auckland Council, Toit_ Te Whenua Land Information New Zealand (LINZ) (2020) . Auckland North, New Zealand 2016-2018. Collected by Aerial Surveys, distributed by OpenTopography and LINZ. https://doi.org/10.5069/G92R3PVR.<br>Hawke's Bay Regional Council et al. (2023). Hawke's Bay, New Zealand 2020-2021. Collected by Ocean Infinity, distributed by OpenTopography and LINZ. https://doi.org/10.5069/G9S75DH2.<br>Marlborough District Council, Toit_ Te Whenua Land Information New Zealand (LINZ) (2023). Marlborough, New Zealand 2020-2022. Collected by Aerial Surveys, distributed by OpenTopography and LINZ. https://doi.org/10.5069/G97D2SB0.<br>Northland Regional Council, Toit_ Te Whenua Land Information New Zealand (LINZ) (2022). Northland, New Zealand 2018-2020. Collected by RPS Group, distributed by OpenTopography and LINZ. https://doi.org/10.5069/G9BR8QDQ.<br>BOPLASS Limited, Toit_ Te Whenua Land Information New Zealand (LINZ) (2023). Bay of Plenty, New Zealand 2019-2022. Collected by Aerial Surveys, distributed by OpenTopography and LINZ. https://doi.org/10.5069/G9W66J0Z.<br>Gisborne District Council, LINZ (2021). Gisborne, New Zealand 2018-2020. Collected by Aerial Surveys, distributed by OpenTopography and Land Information New Zealand (LINZ). https://doi.org/10.5069/G92V2D9X.<br>Environment Southland Regional Council, Toit_ Te Whenua Land Information New Zealand (LINZ) (2024). Southland, New Zealand 2020-2024. Collected by Aerial Surveys, distributed by OpenTopography and LINZ. https://doi.org/10.5069/G9TM78B1.<br>Tasman District Council, Toit_ Te Whenua Land Information New Zealand (LINZ) (2023). Tasman, New Zealand 2020-2022. Collected by Aerial Surveys, distributed by OpenTopography and LINZ. https://doi.org/10.5069/G9S46Q5N.<br>Waikato Regional Council et al. (2024). Waikato, New Zealand 2021. Collected by Ocean Infinity, distributed by OpenTopography and LINZ. https://doi.org/10.5069/G9ZC8136.<br>Environment Canterbury Regional Council et al. (2025). Canterbury, New Zealand 2020-2025. Collected by Aerial Surveys, distributed by OpenTopography and LINZ. https://doi.org/10.5069/G9Q23XF8. | 93 | 2003.5 | 79.7 | 6.8 |

| | | | | | |
|---|---|---|---|---|---|
| | Wellington, New Zealand 2013-2014. Distributed by OpenTopography.<br>https://doi.org/10.5069/G9CV4FPT.<br>https://portal.opentopography.org/datasetMetadata?otCollectionID=OT.042017.2193.2<br>West Coast Regional Council, Toit_ Te Whenua Land Information New Zealand (LINZ) (2025). West Coast, New Zealand 2020-2025. Collected by Aerial Surveys, distributed by OpenTopography and LINZ. | | | | |
| **Lantmateriet** | Lantmateriet Geotorget. https://geotorget.lantmateriet.se/bestallning/produkter | 9 | 834.4 | 22.2 | 1.8 |
| **LidarForScotland** | Scottish Remote Sensing Portal. https://remotesensingdata.gov.scot/data#/list | 5 | 26 | 0.9 | 1.9 |
| **MRNFQuebec** | Ministere des ressources naturelles et des forets. Lidar - Modeles numeriques (terrain, canopee, pente, courbe de niveau).<br>https://www.donneesquebec.ca/recherche/dataset/produits-derives-de-base-du-lidar | 121 | 2339.3 | 80.4 | 5.9 |
| **NASASonoma** | UMD-NASA Carbon Mapping /Sonoma County Vegetation Mapping and LiDAR Program. Distributed by OpenTopography. https://doi.org/10.5069/G9G73BM1. | 1 | 12.4 | 0.5 | 7.3 |
| **NCALM** | Lumbrazo, C. (2021). Hydrologic Effects of Forest Restoration, WA 2021. NCALM. Distributed by OpenTopography. https://doi.org/10.5069/G989142F.<br>South Fork Eel River, CA: Understanding Terrace Formation and Abandonment. Distributed by OpenTopography. https://doi.org/10.5069/G93F4MH1.<br>South Florida Everglades. Distributed by OpenTopography.<br>https://doi.org/10.5069/G9XG9P2Q.<br>Everglades National Park. Distributed by OpenTopography.<br>https://doi.org/10.5069/G9BC3WG0.<br>Greys River, WY Bathymetric Lidar. Distributed by OpenTopography.<br>https://doi.org/10.5069/G9CC0XNT.<br>Herbst, T. (2020). Lassen Volcanic National Park, CA 2019. NCALM. Distributed by OpenTopography.<br>Yosemite, CA: El Portal, Mariposa Grove, Yosemite Canyon & Tuolumne Meadows. Distributed by OpenTopography. https://doi.org/10.5069/G9GQ6VP3.<br>Niobrara River, Nebraska. Distributed by OpenTopography.<br>https://doi.org/10.5069/G9NP22DP.<br>Benjaram, S.S. (2016): Northern Rockies, Montana lidar 2016 airborne lidar survey. NCALM. Distributed by OpenTopography. https://doi.org/10.5069/G9RR1W9H.<br>Yanites, B. (2022). Lidar Survey of Dump Creek, ID 2019. NCALM. Distributed by OpenTopography. https://doi.org/10.5069/G93J3B5W.<br>Tushar Mountains, Utah. Distributed by OpenTopography.<br>https://doi.org/10.5069/G97M05W1. | 31 | 624 | 21 | 7.7 |
| **NCALMHawaii** | Hawaii Kauai Survey. Distributed by OpenTopography. https://doi.org/10.5069/G91V5BWJ. | 1 | 202 | 7.1 | 3.3 |
| **NEON** | NEON (National Ecological Observatory Network). Discrete return LiDAR point cloud (DP1.30003.001). https://data.neonscience.org/data-products/DP1.30003.001<br>NEON.D04.GUAN.DP1.30003.001.2018-05<br>NEON.D04.GUAN.DP1.30003.001.2022-10<br>NEON.D04.CUPE.DP1.30003.001.2022-10<br>NEON.D16.ABBY.DP1.30003.001.2017-06<br>NEON.D16.ABBY.DP1.30003.001.2018-07<br>NEON.D16.ABBY.DP1.30003.001.2019-07<br>NEON.D16.ABBY.DP1.30003.001.2021-07<br>NEON.D16.ABBY.DP1.30003.001.2022-07<br>NEON.D01.BART.DP1.30003.001.2014-06<br>NEON.D01.BART.DP1.30003.001.2016-08<br>NEON.D01.BART.DP1.30003.001.2017-08<br>NEON.D01.BART.DP1.30003.001.2018-08<br>NEON.D01.BART.DP1.30003.001.2019-08<br>NEON.D01.BART.DPI.30003.001.2022-08<br>NEON.D19.BONA.DP1.30003.001.2017-08<br>NEON.D19.BONA.DP1.30003.001.2018-08<br>NEON.D19.BONA.DP1.30003.001.2019-08<br>NEON.D19.BONA.DP1.30003.001.2021-08<br>NEON.D19.BONA.DP1.30003.001.2023-08<br>NEON.D19.BONA.DP3.30024.001.2024-07<br>NEON.D11.CLBJ.DP1.30003.001.2016-04<br>NEON.D11.CLBJ.DP1.30003.001.2017-05 | | | | |

| | |
|---|---|
| | NEON.D11.CLBJ.DP1.30003.001.2018-04<br>NEON.D11.CLBJ.DP1.30003.001.2019-04<br>NEON.D11.CLBJ.DP1.30003.001.2021-06<br>NEON.D11.CLBJ.DP1.30003.001.2022-05<br>NEON.D11.CLBJ.DP3.30024.001.2023-05<br>NEON.D11.CLBJ.DP3.30024.001.2024-05<br>NEON.D19.DEJU.DP1.30003.001.2017-07<br>NEON.D19.DEJU.DP1.30003.001.2018-08<br>NEON.D19.DEJU.DP1.30003.001.2019-08<br>NEON.D19.DEJU.DP1.30003.001.2021-07<br>NEON.D19.DEJU.DP1.30003.001.2023-07<br>NEON.D19.DEJU.DP3.30024.001.2024-07<br>NEON.D08.DELA.DP1.30003.001.2015-07<br>NEON.D08.DELA.DP1.30003.001.2016-05<br>NEON.D08.DELA.DP1.30003.001.2017-05<br>NEON.D08.DELA.DP1.30003.001.2018-04<br>NEON.D08.DELA.DP1.30003.001.2019-04<br>NEON.D08.DELA.DP1.30003.001.2021-05<br>NEON.D03.DSNY.DP1.30003.001.2014-05<br>NEON.D03.DSNY.DP1.30003.001/2016-09<br>NEON.D03.DSNY.DP1.30003.001.2017-09<br>NEON.D03.DSNY.DP1.30003.001.2018-10<br>NEON.D03.DSNY.DP1.30003.001.2019-04<br>NEON.D03.DSNY.DP1.30003.001.2021-09<br>NEON.D07.GRSM.DP1.30003.001.2015-08<br>NEON.D07.GRSM.DP1.30003.001.2016-06<br>NEON.D07.GRSM.DP1.30003.001.2017-10<br>NEON.D07.GRSM.DP1.30003.001.2018-05<br>NEON.D07.GRSM.DP1.30003.001.2021-06<br>NEON.D01.HARV.DP1.30003.001.2014-06<br>NEON.D01.HARV.DP1.30003.001.2016-08<br>NEON.D01.HARV.DP1.30003.001.2017-08<br>NEON.D01.HARV.DP1.30003.001.2018-09<br>NEON.D01.HARV.DP1.30003.001.2019-08<br>NEON.D01.HARV.DP1.30003.001.2022-08<br>NEON.D01.HARV.DP3.30024.001.2024-08<br>NEON.D19.HEAL.DP3.30024.001.2017-07<br>NEON.D19.HEAL.DP3.30024.001.2018-08<br>NEON.D19.HEAL.DP3.30024.001.2019-08<br>NEON.D19.HEAL.DP3.30024.001.2021-07<br>NEON.D19.HEAL.DP3.30024.001.2023-08<br>NEON.D19.HEAL.DP3.30024.001.2024-07<br>NEON.D01.HOPB.DP1.30003.001.2016-08<br>NEON.D01.HOPB.DP1.30003.001.2017-08<br>NEON.D01.HOPB.DP1.30003.001.2019-08<br>NEON.D01.HOPB.DP1.30003.001.2022.08<br>NEON.D03.JERC.DP1.30003.001.2014-05<br>NEON.D03.JERC.DP1.30003.001.2016-09<br>NEON.D03.JERC.DP1.30003.001.2017-09<br>NEON.D03.JERC.DP1.30003.001.2018-09<br>NEON.D03.JERC.DP1.30003.001.2019-09<br>NEON.D03.JERC.DP1.30003.001.2021-09<br>NEON.D08.LENO.DP1.30003.001.2015-07<br>NEON.D08.LENO.DP1.30003.001.2016-05<br>NEON.D08.LENO.DP1.30003.001.2017-05<br>NEON.D08.LENO.DP1.30003.001.2018-04<br>NEON.D08.LENO.DP1.30003.001.2019-05<br>NEON.D08.LENO.DP1.30003.001.2021-05<br>NEON.D08.LENO.DP3.30024.001.2023-05<br>NEON.D08.LENO.DP3.30024.001.2024-05<br>NEON.D05.LIRO.DP1.30003.001.2017-09<br>NEON.D05.LIRO.DP1.30003.001.2020-08<br>NEON.D05.LIRO.DP1.30003.001.2022-06<br>NEON.D16.MCRA.DP1.30003.001.2018-07<br>NEON.D16.MCRA.DP1.30003.001.2021-07<br>NEON.D16.MCRA.DP1.30003.001.2022-07<br>NEON.D07.MLBS.DP1.30003.001.2015-08<br>NEON.D07.MLBS.DP1.30003.001.2017-08<br>NEON.D07.MLBS.DP1.30003.001.2018-05<br>NEON.D07.MLBS.DP1.30003.001.2021-06<br>NEON.D07.MLBS.DP1.30003.001.2022-09<br>NEON.D13.NIWO.DP1.30003.001.2017-09<br>NEON.D13.NIWO.DP1.30003.001.2018-08<br>NEON.D13.NIWO.DP1.30003.001.2019-08<br>NEON.D13.NIWO.DP1.30003.001.2020-08<br>NEON.D13.NIWO.DP3.30024.001.2023-08<br>NEON.D13.NIWO.DP3.30024.001.2024-07<br>NEON.D15.ONAQ.DP3.30024.001.2017-06<br>NEON.D15.ONAQ.DP3.30024.001.2019-05<br>NEON.D15.ONAQ.DP3.30024.001.2021-05 |

| | |
|---|---|
| | NEON.D15.ONAQ.DP3.30024.001.2022-05<br>NEON.D15.ONAQ.DP3.30024.001.2023-04<br>NEON.D07.ORNL.DP1.30003.001.2015-08<br>NEON.D07.ORNL.DP1.30003.001.2016-06<br>NEON.D07.ORNL.DP1.30003.001.2017-09<br>NEON.D07.ORNL.DP1.30003.001.2018-05<br>NEON.D03.OSBS.DP1.30003.001.2014-05<br>NEON.D03.OSBS.DP1.30003.001.2016-09<br>NEON.D03.OSBS.DP1.30003.001.2017-09<br>NEON.D03.OSBS.DP1.30003.001.2018-09<br>NEON.D03.OSBS.DP1.30003.001.2019-04<br>NEON.D03.OSBS.DP1.30003.001.2021-09<br>NEON.D11.PRIN.DP1.30003.001.2016-04<br>NEON.D11.PRIN.DP1.30003.001.2017-05<br>NEON.D11.PRIN.DP1.30003.001.2021-06<br>NEON.D11.PRIN.DP1.30003.001.2022-05<br>NEON.D10.RMNP.DP1.30003.001.2017-07<br>NEON.D10.RMNP.DP1.30003.001.2018-09<br>NEON.D10.RMNP.DP1.30003.001.2020-07<br>NEON.D10.RMNP.DP1.30003.001.2022-07<br>NEON.D10.RMNP.DP3.30024.001.2023-08<br>NEON.D10.RMNP.DP3.30024.001.2024-07<br>NEON.D02.SCBI.DP1.30003.001.2016-07<br>NEON.D02.SCBI.DP1.30003.001.2017-07<br>NEON.D02.SCBI.DP1.30003.001.2019-06<br>NEON.D02.SCBI.DP1.30003.001.2021-08<br>NEON.D02.SCBI.DP1.30003.001.2022-05<br>NEON.D02.SCBI.DP1.30003.001.2023-06<br>NEON.D02.SERC.DP1.30003.001.2016-07<br>NEON.D02.SERC.DP1.30003.001.2017-07<br>NEON.D02.SERC.DP1.30003.001.2017-08<br>NEON.D02.SERC.DP1.30003.001.2019-05<br>NEON.D02.SERC.DP1.30003.001.2021-08<br>NEON.D02.SERC.DP1.30003.001.2022-05<br>NEON.D17.SJER.DP1.30003.001.2013-06<br>NEON.D17.SJER.DP1.30003.001.2017-03<br>NEON.D17.SJER.DP1.30003.001.2018-03<br>NEON.D17.SJER.DP1.30003.001.2019-03<br>NEON.D17.SJER.DP1.30003.001.2021-03<br>NEON.D17.SJER.DP3.30024.001.2023-04<br>NEON.D17.SJER.DP3.30024.001.2024-04<br>NEON.D17.SOAP.DP1.30003.001.2013-06<br>NEON.D17.SOAP.DP1.30003.001.2017-07<br>NEON.D17.SOAP.DP1.30003.001.2018-06<br>NEON.D17.SOAP.DP1.30003.001.2019-06<br>NEON.D17.SOAP.DP1.30003.001.2021-07<br>NEON.D14.SRER.DP1.30003.001.2017-08<br>NEON.D14.SRER.DP1.30003.001.2018-08<br>NEON.D14.SRER.DP1.30003.001.2019-09<br>NEON.D14.SRER.DP1.30003.001.2021-09<br>NEON.D14.SRER.DP1.30003.001.2022-08<br>NEON.D05.STEI.DP1.30003.001.2016-09<br>NEON.D05.STEI.DP1.30003.001.2017-08<br>NEON.D05.STEI.DP1.30003.001.2019-06<br>NEON.D05.STEI.DP1.30003.001.2022-06<br>NEON.D08.TALL.DP1.30003.001.2015-07<br>NEON.D08.TALL.DP1.30003.001.2016-05<br>NEON.D08.TALL.DP1.30003.001.2017-05<br>NEON.D08.TALL.DP1.30003.001.2018-04<br>NEON.D08.TALL.DP1.30003.001.2019-04<br>NEON.D08.TALL.DP1.30003.001.2021-05<br>NEON.D08.TALL.DP3.30024.001.2023-05<br>NEON.D08.TALL.DP3.30024.001.2024-05<br>NEON.D17.TEAK.DP1.30003.001.2013-06<br>NEON.D17.TEAK.DP1.30003.001.2017-06<br>NEON.D17.TEAK.DP1.30003.001.2018-06<br>NEON.D17.TEAK.DP1.30003.001.2019-06<br>NEON.D17.TEAK.DP1.30003.001.2021-07<br>NEON.D05.TREE.DP1.30003.001.2016-09<br>NEON.D05.TREE.DP1.30003.001.2017-08<br>NEON.D05.TREE.DP1.30003.001.2019-06<br>NEON.D05.TREE.DP1.30003.001.2020-08<br>NEON.D05.TREE.DP1.30003.001.2022-06<br>NEON.D05.TREE.DP3.30024.001.2024-07<br>NEON.D06.UKFS.DP1.30003.001.2016-07<br>NEON.D06.UKFS.DP1.30003.001.2017-06<br>NEON.D06.UKFS.DP1.30003.001.2018-06<br>NEON.D06.UKFS.DP1.30003.001.2019-05<br>NEON.D06.UKFS.DP1.30003.001.2020-07<br>NEON.D06.UKFS.DP1.30003.001.2022-06 |

| | | | | | |
|---|---|---|---|---|---|
| | NEON.D06.UKFS.DP3.20024.001.2023-05<br>NEON.D06.UKFS.DP3.20024.001.2024-06<br>NEON.D05.UNDE.DP1.30003.001.2016-09<br>NEON.D05.UNDE.DP1.30003.001.2017-09<br>NEON.D05.UNDE.DP1.30003.001.2019-06<br>NEON.D05.UNDE.DP1.30003.001.2020-08<br>NEON.D05.UNDE.DP1.30003.001.2022-06<br>NEON.D13.WLOU.DP1.30003.001.2017-09<br>NEON.D13.WLOU.DP1.30003.001.2019-08<br>NEON.D13.WLOU.DP1.30003.001.2020-08<br>NEON.D16.WREF.DP1.30003.001.2017-06<br>NEON.D16.WREF.DP1.30003.001.2018-07<br>NEON.D16.WREF.DP1.30003.001.2019-07<br>NEON.D16.WREF.DP1.30003.001.2021-07<br>NEON.D16.WREF.DP1.30003.001.2022-07<br>NEON.D12.YELL.DP1.30003.001.2018-07<br>NEON.D12.YELL.DP1.30003.001.2019-07<br>NEON.D12.YELL.DP1.30003.001.2020-07<br>NEON.D12.YELL.DP1.30003.001.2022-06<br>NEON.D20.PUUM.DP1.30003.001.2019-01<br>NEON.D20.PUUM.DP1.30003.001.2020-01 | | | | |
| **NEONprototype** | NEON (National Ecological Observatory Network). Discrete return LiDAR point cloud (DP1.30003.001). https://data.neonscience.org | 1 | 172 | 5 | 2.3 |
| **NERCGroundDataSolutions** | NERC Airborne Research and Survey Facility (ARSF) Remote Sensing Data.<br>Coomes, D.; Jackson, T. (2022): Airborne LiDAR and RGB imagery from Sepilok Reserve and Danum Valley in Malaysia in 2020. NERC EDS Centre for Environmental Data Analysis, 03 October 2022. doi:10.5285/dd4d20c8626f4b9d99bc14358b1b50fe | 5 | 72.5 | 2.6 | 32.4 |
| **NERCMalaysia2014** | NERC Airborne Research and Survey Facility (ARSF) Remote Sensing Data.<br>ARSF flight MA14_08. https://data.ceda.ac.uk/neodc/arsf/2014/MA14_08<br>ARSF flight MA14_10. https://data.ceda.ac.uk/neodc/arsf/2014/MA14_10<br>ARSF flight MA14_11. https://data.ceda.ac.uk/neodc/arsf/2014/MA14_11<br>NERC ARSF, Burslem, Coomes (2019): ARSF 2014_309 - MA14_14 Flight: Airborne remote sensing measurements. Centre for Environmental Data Analysis, 25 September 2019. doi:10.5285/c708cad9950c45b1af0d4e9ca944f09a.<br>ARSF flight MA14_21. https://data.ceda.ac.uk/neodc/arsf/2014/MA14_21<br>NERC Airborne Research Facility (2020): NERC-ARF 2014_289 - RG13_06. Centre for Environmental Data Analysis. doi:10.5285/1a072df93a19434e85b4fff7fabfdafa. https://data.ceda.ac.uk/neodc/arsf/2014/RG13_06 | 37 | 1622.5 | 56.4 | 9.7 |
| **NERCVolcanoes** | NERC Airborne Research and Survey Facility (ARSF) Remote Sensing Data.<br>ARSF flight ET12_14. Centre for Environmental Data Analysis (CEDA). https://data.ceda.ac.uk/neodc/arsf/2012/ET12_14/ET12_14-2012_324_Boset<br>ARSF flight ET12_17. Centre for Environmental Data Analysis (CEDA). https://data.ceda.ac.uk/neodc/arsf/2012/ET12_17/ET12_17-2012_321_Corbetti_Alutu<br>ARSF flight ET12_17. Centre for Environmental Data Analysis (CEDA). https://data.ceda.ac.uk/neodc/arsf/2012/ET12_17/ET12_17-2012_320_Corbetti<br>ARSF flight ET12_18. Centre for Environmental Data Analysis (CEDA). https://data.ceda.ac.uk/neodc/arsf/2012/ET12_18/ET12_18-2012_318_Rift<br>ARSF flight ET12_18. Centre for Environmental Data Analysis (CEDA). https://data.ceda.ac.uk/neodc/arsf/2012/ET12_18/ET12_18-2012_319_Rift | 7 | 947.3 | 31.5 | 1.9 |
| **NERCarsf** | NERC Airborne Research and Survey Facility (ARSF) Remote Sensing Data.<br>ARSF flight GB14_04. https://data.ceda.ac.uk/neodc/arsf/2014/GB14_04/GB14_04-2014_219b_Aberfoyle<br>ARSF flight GB09_11. https://data.ceda.ac.uk/neodc/arsf/2009/GB09_11/GB09_11-2009_331_Abernethy<br>ARSF flight GB12_04. https://data.ceda.ac.uk/neodc/arsf/2012/GB12_04/GB12_04-2012_086_Eaves_Wood<br>ARSF flight GB12_04. https://data.ceda.ac.uk/neodc/arsf/2012/GB12_04/GB12_04-2012_291b_Eaves_Wood<br>ARSF flight GB12_04. https://data.ceda.ac.uk/neodc/arsf/2014/GB12_04/GB12_04-2014_083_Eaves_Wood<br>ARSF flight GB13_05. https://data.ceda.ac.uk/neodc/arsf/2013/GB13_05/GB13_05-2013_144_Glenmore<br>ARSF flight RG13_08. https://data.ceda.ac.uk/neodc/arsf/2014/RG13_08/RG13_08-2014_175b_Wytham_Woods_Plus_Airsar | 10 | 142.8 | 5 | 6.1 |
| **NISAR_Chaco** | original data contributed to GCA | 20 | 215.5 | 7.4 | 10.6 |
| **NISAR_Krodsherad** | de Lera Garrido A., Gobakken T., Ørka H. O., Næsset E., Bollandsås O. M. (2020). Reuse of field data in ALS-assisted forest inventory. Silva Fennica vol. 54 no. 5 article id 10272. https://doi.org/10.14214/sf.10272 | 1 | 72.8 | 2.5 | 11.9 |
| **NISAR_MadredeDios** | original data contributed to GCA | 5 | 17.2 | 0.7 | 112 |
| **NISAR_Madrid** | original data contributed to GCA | 7 | 29.1 | 1.3 | 9.5 |
| **NOAA** | NOAA digital coast: https://coast.noaa.gov/dataviewer/ | 16 | 568.4 | 19.8 | 10.1 |

| | | | | | |
|---|---|---|---|---|---|
| NRCAN | LiDAR Point Clouds - CanElevation Series. https://open.canada.ca/data/en/dataset/7069387e-9986-4297-9f55-0288e9676947 | 37 | 1596 | 50 | 12.1 |
| NZAlpineFault | Langridge, R.M. (2022). Alpine Fault, NZ 2015. Collected by AAM New Zealand Limited (AAM). Distributed by OpenTopography. https://doi.org/10.5069/G9M61HGF. | 4 | 139.7 | 6.2 | 19.4 |
| NZLakeSumner | Langridge, R.M.(2022). Western Hope Fault, NZ 2010. Collected by New Zealand Aerial Mapping Limited (NZAM). Distributed by OpenTopography. https://doi.org/10.5069/G9QZ285D. | 1 | 34.5 | 1.3 | 3 |
| National Land Survey of Finland | Laser scanning data 0,5 p. https://www.maanmittauslaitos.fi/en/maps-and-spatial-data/expert-users/product-descriptions/laser-scanning-data-05-p | 12 | 459.1 | 13.1 | 3 |
| NgangaoTaitaHills | Janne Heiskanen et al. 2015. Use of airborne lidar for estimating canopy gap fraction and leaf area index of tropical montane forests. International Journal of Remote Sensing, 2569-2583, DOI: 10.1080/01431161.2015.1041177 | 1 | 6.8 | 0.2 | 11.1 |
| OntarioSPL | Ontario Forest Resources Inventory leaf-on LiDAR. https://geohub.lio.gov.on.ca/maps/lio::forest-resources-inventory-leaf-on-lidar/about | 55 | 1509.9 | 45 | 43 |
| PaintRockUAV | original data contributed to GCA | 2 | 1.1 | 0.1 | 67.8 |
| PeatlandsIND | Vernimmen et al. 2019. Creating a Lowland and Peatland Landscape Digital Terrain Model (DTM) from Interpolated Partial Coverage LiDAR Data for Central Kalimantan and East Sumatra, Indonesia. Remote Sens. 11, 1152. https://doi.org/10.3390/rs11101152 | 3 | 81 | 2.6 | 5.9 |
| PontaldoParanapanema | original data contributed to GCA | 75 | 236.6 | 9.2 | 7.9 |
| SLU | Huo et al. 2022. Towards low vegetation identification: A new method for tree crown segmentation from LiDAR data based on a symmetrical structure detection algorithm. Remote Sensing of Environment 270, 112857 | 3 | 128.5 | 3.5 | 27.9 |
| STRIPanama | ForestGEO Smithsonian. (2024). 2023 high-resolution airborne LiDAR data for Barro Colorado Island and other Smithsonian ForestGEO Sites in Central Panama. Smithsonian Research Data Repository. doi:10.60635/C34W2W. | 15 | 113.3 | 4.3 | 19.2 |
| SYSSIFOSS | Weiser et al. 2022. Individual tree point clouds and tree measurements from multi-platform laser scanning in German forests, Earth Syst. Sci. Data, 14, 2989Ð3012, https://doi.org/10.5194/essd-14-2989-2022 | 23 | 19.4 | 0.9 | 155.3 |
| SaoPaulo | Sao Paulo City Hall (PMSP) (2024). Sao Paulo, Brazil Lidar Survey 2017. Distributed by OpenTopography. https://doi.org/10.5069/G9NV9GD1. | 2 | 270.3 | 8.8 | 16.1 |
| SilaNP | Puletti, Nicola. (2020). Sila National Park - 3D Point cloud data (Version 1) [Data set]. Zenodo. https://doi.org/10.5281/zenodo.3633629 | 1 | 29.1 | 1.1 | 6.5 |
| SumavaBayerwald | Latifi et al. 2021. A laboratory for conceiving Essential Biodiversity Variables (EBVs). The Data pool initiative for the Bohemian Forest Ecosystem. Methods in Ecology and Evolution, 12, 2073-2083. https://doi.org/10.1111/2041-210X.13695 | 2 | 1215.5 | 50.8 | 29.5 |
| SustainableLandscapes | dos-Santos et al. 2019. LiDAR Surveys over Selected Forest Research Sites, Brazilian Amazon, 2008-2018. ORNL DAAC, Oak Ridge, Tennessee, USA. https://doi.org/10.3334/ORNLDAAC/1644; https://www.paisagenslidar.cnptia.embrapa.br/geonetwork/<br>Keller, Michael; Batistella, Mateus; Gorgens, Eric Bastos, 2024, ""LiDAR survey on 765 hectares in Tumbira, Para, Brazil, 2021."", https://doi.org/10.48432/TMJT7O, Redape, V2<br>Gorgens et al. 2023. Out of steady state: Tracking canopy gap dynamics across Brazilian Amazon. Biotropica, 55, 755-766. https://doi.org/10.1111/btp.13226<br>dos Santos, Maiza Nara; Keller, Michael; Batistella, Mateus, 2023, ""LiDAR survey on 176.2 hectares in Ucayali, Peru in 2017."", https://doi.org/10.48432/I98OSL, Redape, V1<br>dos Santos, Maiza Nara; Keller, Michael; Batistella, Mateus, 2023, ""LiDAR survey on 223.5 hectares in Ucayali, Peru in 2017."", https://doi.org/10.48432/RW3KGS, Redape, V1<br>dos Santos, Maiza Nara; Keller, Michael; Batistella, Mateus, 2023, ""LiDAR survey on 374.7 hectares in Ucayali, Peru in 2017."", https://doi.org/10.48432/L0IRCB, Redape, V1<br>dos Santos, Maiza Nara; Keller, Michael; Batistella, Mateus, 2023, ""LiDAR survey on 244 hectares in Ucayali, Peru in 2017."", https://doi.org/10.48432/9OUYDX, Redape, V1 | 253 | 933.3 | 39.5 | 19.4 |
| TCDWY | Teton Conservation District, Wyoming Lidar. Distributed by OpenTopography. https://doi.org/10.5069/G9F769GN. | 1 | 457 | 13.8 | 1.8 |
| TEAMLaSelva2009 | Beckley 2019. TEAM lidar data over La Selva, Costa Rica 2009. Northrop Grumman Foundation. Distributed by OpenTopography. https://doi.org/10.5069/G9P8491K. Accessed: 2023-07-04 | 1 | 50 | 1.6 | 2.1 |
| TERNSuperSites | Department of the Environment, T. (2021). Airborne Hyperspectral and LiDAR Data - Australian Field Sites. Version 1.0. Terrestrial Ecosystem Research Network. Dataset. https://portal.tern.org.au/metadata/4ff0b4c9-cfa0-4d09-9520-b5402adc583f | 12 | 264.7 | 8.2 | 20.7 |
| TWDEF | Teich & Tarboton 2016. TW Daniels Experimental Forest (TWDEF) Lidar, HydroShare, https://doi.org/10.4211/hs.36f3314971a547bc8bc72dc60d6bd03c | 6 | 29.7 | 1.3 | 3.1 |
| TeraiArcLandscape | Parvez Rana et al. 2016. Optimizing the number of training areas for modeling above-ground biomass with ALS and multispectral remote sensing in subtropical Nepal. International Journal of Applied Earth Observation and Geoinformation 49, 52-62. | 20 | 1202.6 | 16.3 | 1.2 |

| | | | | | |
|---|---|---|---|---|---|
| **Tiantong** | original data contributed to GCA | 1 | 0.2 | 0 | 102.8 |
| **TrailValleyCreek** | Katharina et al. 2018. Airborne Laser Scanning (ALS) Point Clouds of Trail Valley Creek, NWT, Canada (2016). PANGAEA. Https://doi.org/10.1594/PANGAEA.894884 | 1 | 106.9 | 3.7 | 3.8 |
| **TraunsteinUFZ** | Knapp et al. 2020. Structure metrics to generalize biomass estimation from lidar across forest types from different continents. Remote Sensing of Environment 2020, 111597. | 2 | 50 | 1.8 | 16.5 |
| **Trentino** | Provincia Autonoma di Trento. https://siat.provincia.tn.it/stem/ | 2 | 69.6 | 2.9 | 9.9 |
| **UAVPKU** | original data contributed to GCA | 8 | 0.7 | 0 | 160.1 |
| **UAVplotsCZ** | original data contributed to GCA | 6 | 3 | 0.1 | 968.1 |
| **UAVplotsJP** | Takeshige et al. 2025. High-resolution digital canopy height models, terrain models, ortho-mosaic photos, and canopy tree crown shapes derived from UAV-borne LiDAR at 22 tree census plots across Japanese natural forests. Ecol. Research 40, 657-670. | 22 | 2.3 | 0.1 | 1011.1 |
| **UGKK SR** | UGKK SR. www.geoportal.sk | 10 | 19.8 | 0.8 | 29.5 |
| **USFS** | U.S. Forest Service Region 5 Remote Sensing Lab Information Management Staff. (2022). USFS Kern Plateau Lidar, CA 2011. Collected by Watershed Sciences, Inc. Distributed by OpenTopography. https://doi.org/10.5069/G9MC8X7N.<br>U.S. Forest Service Region 5 Remote Sensing Lab Information Management Staff. (2023). USFS Mill Flat Creek Lidar, CA 2012. Collected by Watershed Sciences, Inc. Distributed by OpenTopography. https://doi.org/10.5069/G9WM1BMS.<br>U.S. Forest Service Region 5 Remote Sensing Lab Information Management Staff. (2023). USFS Willow Creek Lidar, CA 2012. Collected by Watershed Sciences, Inc. Distributed by OpenTopography. https://doi.org/10.5069/G9G73BWM. | 6 | 127.3 | 4.3 | 10.2 |
| **WWFCongo** | UCLA, WWF, BMUB, KFW 2017. Carbon Map of DRC: High resolution carbon distribution in forests of Democratic Republic of Congo. Summary Report. Available at globil.panda.org. Last accessed on 28/02/2022 | 218 | 4365.7 | 153.4 | 3.2 |
| **WaterDayekou** | Ni et al. 2013. WATER: Dataset of airborne LiDAR mission at the super site in the Dayekou watershed flight zone on Jun. 23, 2008. A Big Earth Data Platform for Three Poles. DOI: 10.3972/water973.0220.db. | 1 | 56.8 | 1.8 | 1.7 |
| **WythamUAV** | original data contributed to GCA | 1 | 3.5 | 0.2 | 109.9 |
| **swissSURFACE3D** | (c)swisstopo. https://www.swisstopo.admin.ch/de/hoehenmodell-swisssurface3d | 10 | 231 | 8.3 | 12.7 |

**Table S1: GCA datasets.** Overview of datasets included in the GCA database. Shown is an informal dataset name, the associated citations and links to the original data, the count of geo-units belonging to the particular dataset, the total area covered by the dataset (in km2, including repeat acquisitions), the total size of the derived products (in GB), and the mean pulse density.

| PRODUCT | DESCRIPTION |
|---|---|
| **summary_processing.csv** | Metadata and processing summary statistics. Useful for quick assessments of scans. |
| **outline_localCRS.shp** | Outline of the scan in local coordinate reference system. Includes the same information as summary_processing.csv. Useful to obtain metadata and processing details. |
| **outline_WGS84.shp (cpg/dbh/prj)** | Outline of the airborne laser scan in longitude-latitude. Includes the same information as summary_processing.csv. Useful to obtain metadata and processing details and to combine multiple outlines for an overview of scan locations. |
| **dtm.tif** | Digital terrain model (DTM). The default and most robust DTM. Based on the default LAStools lasground_new classification but refined to improve robustness in steep areas (slopes > 40 degrees). Recommended for comparative analyses across space or time. |
| **dtm_lasdef.tif** | Digital terrain model (DTM). Based on the default LAStools lasground_new classification without improvements in steep areas. Recommended for comparative analyses across space or time except in montane regions. |
| **dtm_supplied.tif** | Digital terrain model (DTM). DTM supplied by data provider. Not always available and not comparable among data providers. Useful for quality checks particularly when containing manual corrections. Not recommended for comparative analyses across space or time. |
| **dtm_lasfine.tif** | Digital terrain model (DTM). DTM based on alternative LAStools settings. Finer resolution of the ground but not robust to pulse density. Useful for high quality scans (> 30 pulses $m^{-2}$) when additional details are required. Not recommended for comparative analyses across space or time. |
| **dtm_highest.tif** | Digital terrain model (DTM). Elevation estimates based on highest ground return per pixel. Not interpolated and therefore not useful for terrain or canopy modelling. Not recommended for analysis except for quality checks e.g. density of ground sampling. |
| **dsm_lspikefree.tif / chm_lspikefree.tif** | Digital surface model (DSM) and canopy height model (CHM), where CHM = DSM – DTM (dtm.tif). Robust DSM/CHM that creates a "spike-free" interpolation of surfaces while adjusting to local pulse densities. Interpretable as top canopy height. Applicable down to 2 pulses per m2. Limited use for individual tree delineation due to smoothing of edges. Recommended for comparative analyses across space and time. |
| **dsm_tin.tif / chm_tin.tif** | Digital surface model (DSM) and canopy height model (CHM), where CHM = DSM – DTM (dtm.tif). Robust DSM/CHM based on interpolation of first returns via a triangulated irregular network (TIN). Interpretable as the mean height of light interception. Limited use for individual tree delineation due to highly irregular surface. Recommended for comparative analyses across space and time. |
| **dsm_highest.tif / chm_highest.tif** | Digital surface model (DSM) and canopy height model (CHM), where CHM = DSM – DTM (dtm.tif). Height estimates based on highest return per pixel. Interpretable as canopy top height. Not robust. Useful for tree crown delineation. Not recommended for comparative analyses across space or time. |
| **mask_cloud.tif** | Mask layer for regions with cloud artefacts. Excludes (sets NA) any pixel where top canopy height exceeds maximum tree height at local (variable) or global (125 m) level. This mask is included in mask_combined.tif and should always be applied. |
| **mask_pd02.tif** | Mask layer for regions with low pulse density. Excludes (sets NA) areas with densities below 2 pulses $m^{-2}$. Included in mask_combined.tif and should always be applied. |
| **mask_pd04.tif** | Same as previous layer but with 4 pulses $m^{-2}$. |
| **mask_noground.tif** | Mask layer for regions devoid of ground points. Excludes (sets NA) areas with insufficient sampling. Included in mask_combined.tif and should always be applied. |
| **mask_unstabledtm.tif** | Mask layer for regions where DTM varies strongly depending on LAStools settings. Useful to improve robustness but may discard valid high-elevation or rugged regions. |
| **mask_steep.tif** | Mask layer for steep regions where DTM varies depending on LAStools settings. Useful to improve robustness but may discard valid high-elevation or rugged regions. |
| **mask_combined.tif** | Mask layer that combines: mask_cloud.tif mask_noground.tif mask_pd02.tif. Minimum mask to improve robustness of the inferred terrain and canopy models. Consider combining with mask_steep.tif or mask_unstabledtm.tif for maximum robustness. |

| | |
|---|---|
| **scanangle_abs.tif** | Absolute scan angle raster. Average scan angle of points registered in each grid cell. Useful to check flight patterns and assess scan quality. |
| **pulsedensity.tif** | Pulse density using first returns. The default pulse density product. Useful for quality checks and as predictor in models. |
| **pulsedensity_lastreturn.tif** | Pulse density using last returns. Useful for quality checks and as predictor in models. |
| **pulsedensity_scanangle20.tif** | Pulse density using first returns for laser pulses with absolute scan angles < 20 degrees. Useful to find high quality scan regions. |
| **classification_supplied.tif** | Rasterized classifications supplied by data provider. Useful to check for noise classifications that may have been overlooked or mapping of built-up area. |

**Table S2: GCA products.** Description of the GCA products and possible uses. All raster products are provided with a spatial grain of 1 $m^2$.

| Biome | Vegetation class |
|---|---|
| Deserts & Xeric Shrublands | Desert |
| N/A | Desert |
| Boreal Forests/Taiga | Closed canopy |
| Flooded Grasslands & Savannas | Open canopy |
| Mangroves | Closed canopy |
| Mediterranean Forests, Woodlands & Scrub | Open canopy |
| Montane Grasslands & Shrublands | Open canopy |
| Temperate Broadleaf & Mixed Forests | Closed canopy |
| Temperate Conifer Forests | Closed canopy |
| Temperate Grasslands, Savannas & Shrublands | Open canopy |
| Tropical & Subtropical Coniferous Forests | Closed canopy |
| Tropical & Subtropical Dry Broadleaf Forests | Closed canopy |
| Tropical & Subtropical Grasslands, Savannas & Shrublands | Open canopy |
| Tropical & Subtropical Moist Broadleaf Forests | Closed canopy |
| Tundra | Open canopy |

**Table S3: Mapping of biomes onto vegetation classes.** Mapping of biomes (Dinerstein et al. 2017) onto three discrete vegetation classes for Figure 1.

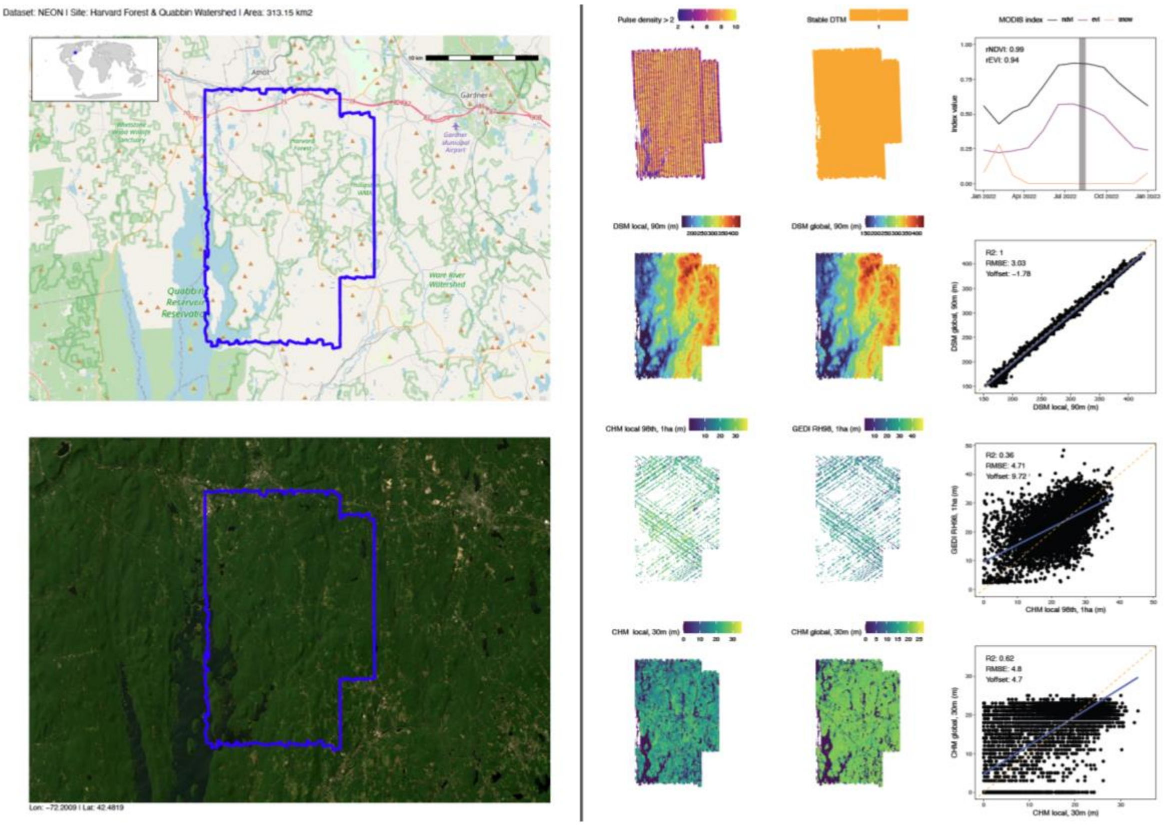

**Figure S1: Processing report at Harvard Forest.** Shown is the automatically generated processing report at Harvard Forest (Northeastern United States). A high-resolution version of this report is provided separately as part of the Supplementary Materials (report_sample_HarvardForest.pdf). Vegetation greenness and snow cover are not directly computed from MODIS-derived values during the acquisition period (Didan, 2025), but from a derived monthly climatology so that $rNDVI_{scan} = NDVI_{scan}/NDVI_{max}$ and $rEVI_{scan} = EVI_{scan}/EVI_{max}$, where $NDVI_{scan}$ and $EVI_{scan}$ are the average values of NDVI and EVI during the acquisition month(s), and $NDVI_{max}$ and $EVI_{max}$ the months with maximum greenness. For comparisons with the global DSM and CHM, we averaged the ALS-derived DSMs and CHMs at the target resolution (90 m and 30 m, respectively). For the comparison with GEDI-based height estimates, we first computed the 98th percentile of canopy height at 20 m resolution (~ GEDI footprint size) for each scan, and then took the mean of the 20 m pixels at 100 m resolution.

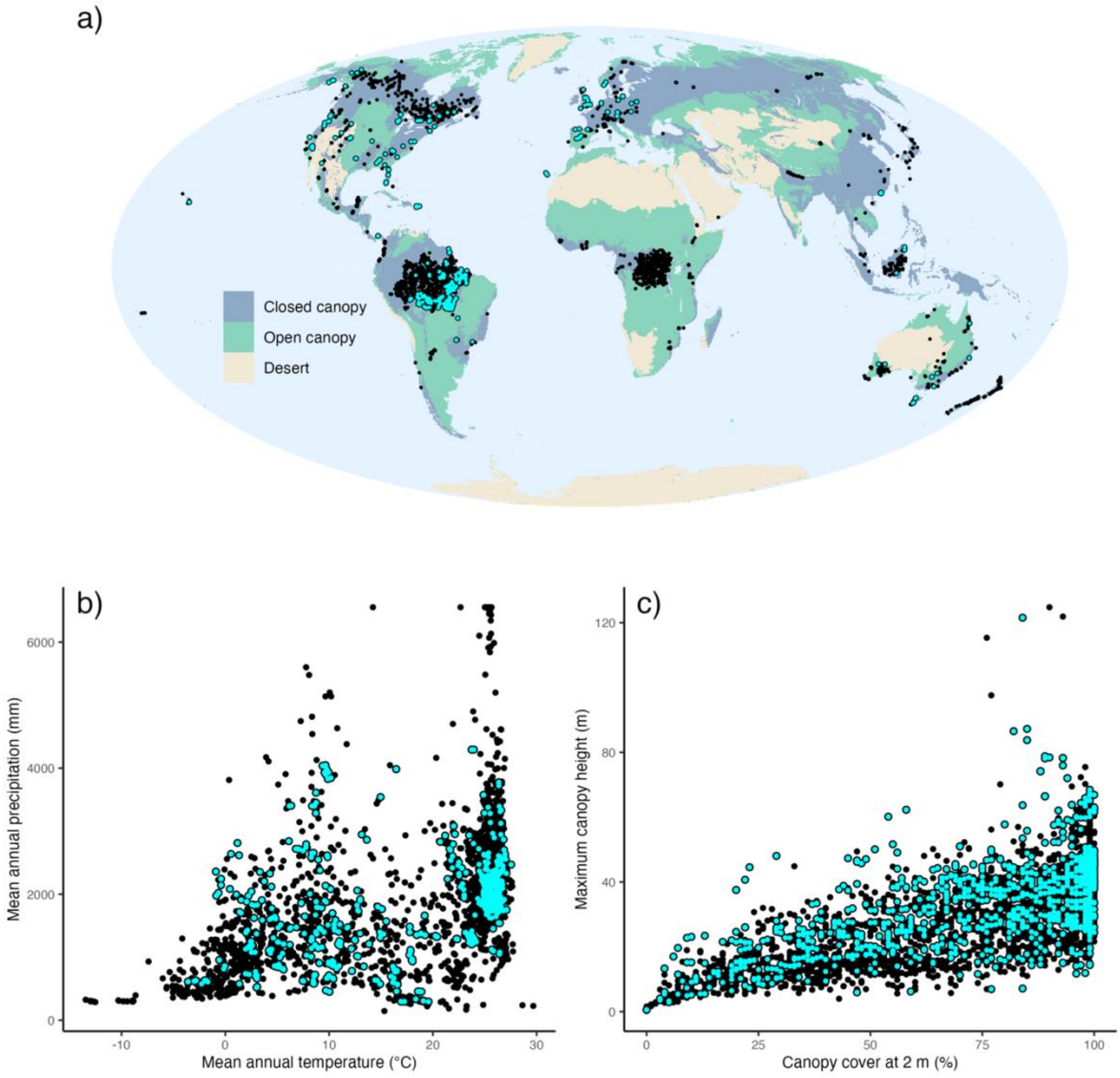

**Figure S2: The Global Canopy Atlas, with repeat scan areas.** This figure replicates panels a-c in Fig. 2 in the main text, but highlights locations with repeat scan data (light blue dots) vs. single scan locations (black dots).

### Case Study #1

| Site | Country | Year | Pulse density ($m^{-2}$) | Area ($km^2$) |
|---|---|---|---|---|
| Chaco | Argentina | 2023 | 2.8 | 11 |
| Normanby River | Australia | 2019 | 32.6 | 34 |
| Mount Fincham | Australia | 2021 | 8.4 | 36 |
| Credo | Australia | 2021 | 23.2 | 25 |
| Rio Jari | Brazil | 2020 | 7.5 | 4 |
| Peace Athabasca | Canada | 2019 | 9.3 | 11 |
| Nahuelbuta | Chile | 2008 | 2.8 | 32 |
| Odzala | Congo | 2021 | 252.7 | 8 |
| Bavarian Forest | Germany | 2020 | 15.8 | 122 |
| Castle Peak | Hong Kong | 2019 | 80.9 | 8 |
| Sila | Italy | 2019 | 6.5 | 29 |
| Sepilok | Malaysia | 2020 | 38.0 | 29 |
| Gorongosa | Mozambique | 2023 | 62.3 | 34 |
| Hawkes Bay Kaweka | New Zealand | 2020 | 10.2 | 15 |
| Hoydedata | Norway | 2019 | 4.0 | 53 |
| Gigante | Panama | 2023 | 29.6 | 2 |
| Rio Acre (FODEX) | Peru | 2019 | 394.1 | 3 |
| La Gomera | Spain | 2023 | 7.6 | 252 |
| Galiuro Wilderness | United States | 2020 | 13.8 | 41 |
| Rocky Mountains | United States | 2022 | 10.2 | 195 |

**Table S1.1: Characteristics of the 20 airborne laser scans.** Shown are the characteristics of the 20 airborne laser scans used to validate the global canopy height models.

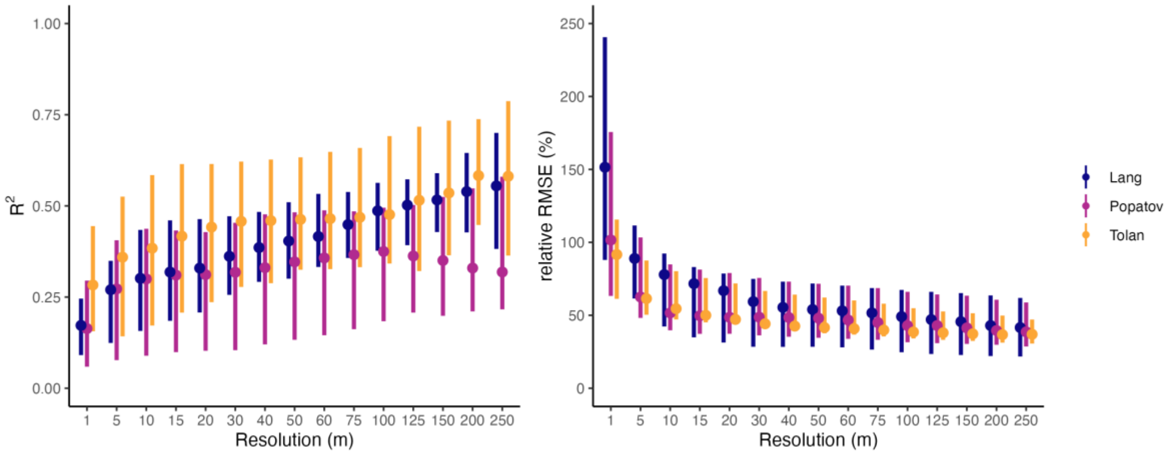

**Figure S1.1: Predictive power of maximum canopy height.** Same as panels a) and b) in Fig. 2 in the main text, but with predictions for maximum height (99th percentile of height) instead of mean height at each resolution.

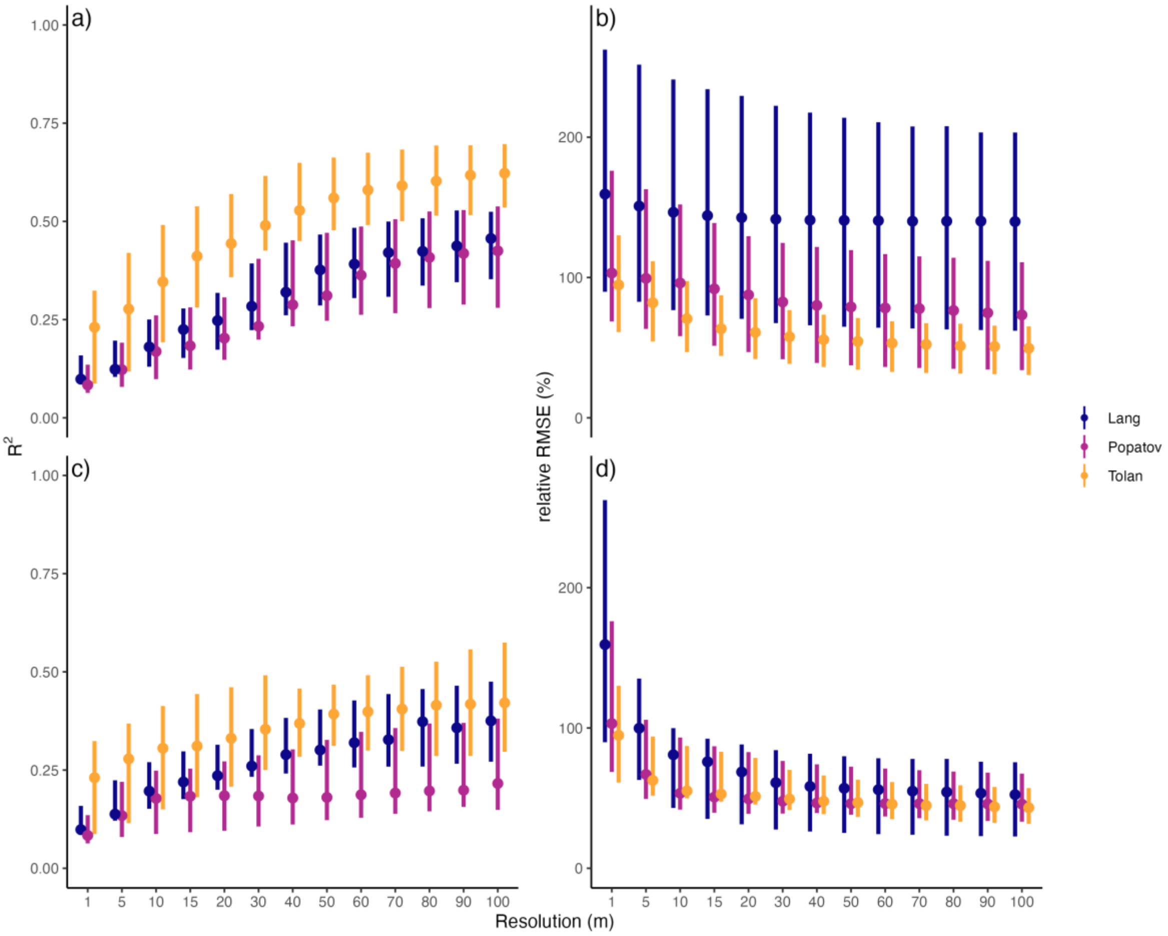

**Figure S1.2: Predictive power, standardized at 1 km².** Same as panels a) and b) in Fig. 2 in the main text and a) and b) in Fig. S1.1, but all statistics are now calculated at a standardized 1 $km^2$ scale instead of across entire landscapes. Panels a) and b) show results for aggregation via mean values, panels c) and d) aggregation with the $99^{th}$ percentile of canopy height. Since 1 $km^2$ are substantially smaller than entire landscapes, we only calculated statistics up to a resolution of 100 m to limit sampling noise.

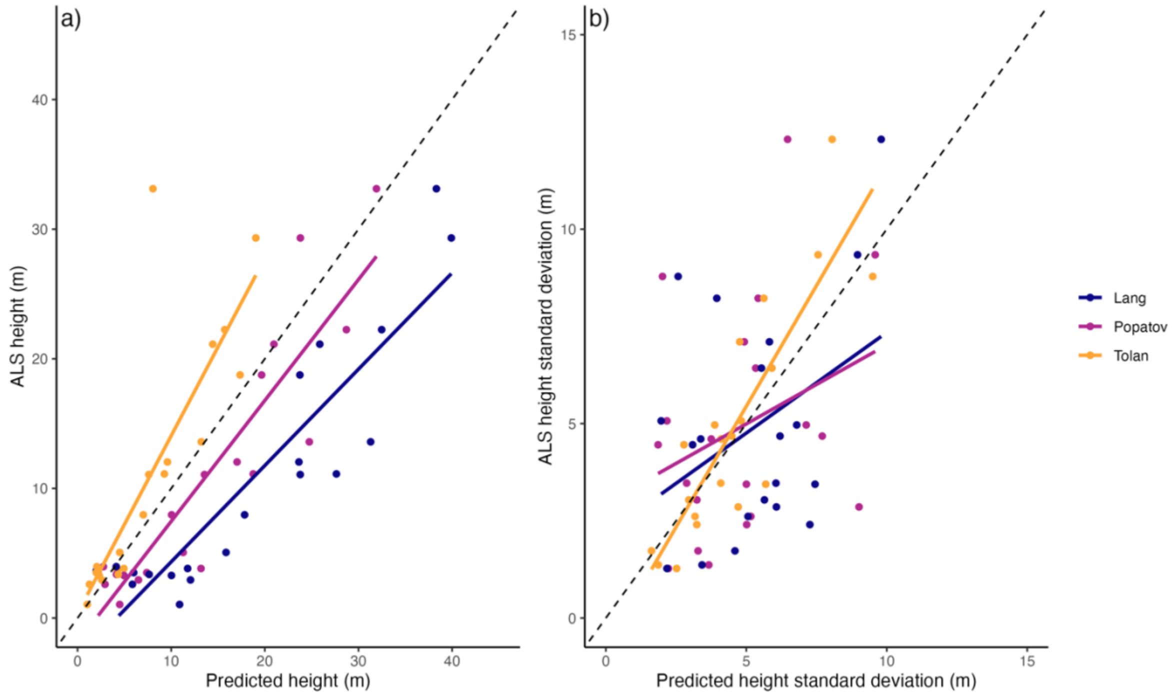

**Figure S1.3: Predictions across landscapes.** Shown are the predicted mean heights at 30 m resolution as well as the predicted standard deviations of 30 m pixels for each of the 20 landscapes from the Tolan, Lang and Potapov models, plotted against the corresponding values derived from ALS acquisitions. Note that Tolan has the clearest outlier in terms of canopy height (corresponding to the site seen in Fig. S1.7), but better represents local variability than the other two models.

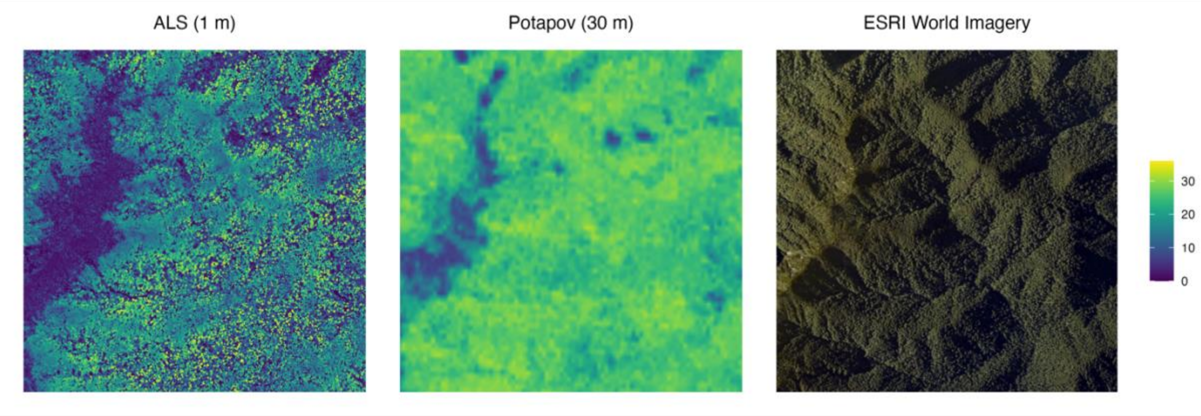

**Figure S1.4: Site with largest absolute errors for Potapov model.** Shown are canopy height models derived from ALS and the Potapov CHM, as well as the corresponding ESRI world imagery data over a 2 km x 2km extent in Hawkes Bay, New Zealand. This is the site with the largest absolute RMSE for the Potapov model. The colour scale on the right refers the height of the two canopy height models (in m). The mean height in the ALS model across the entire site is 13.6 m, while Potapov predicts 24.7 m.

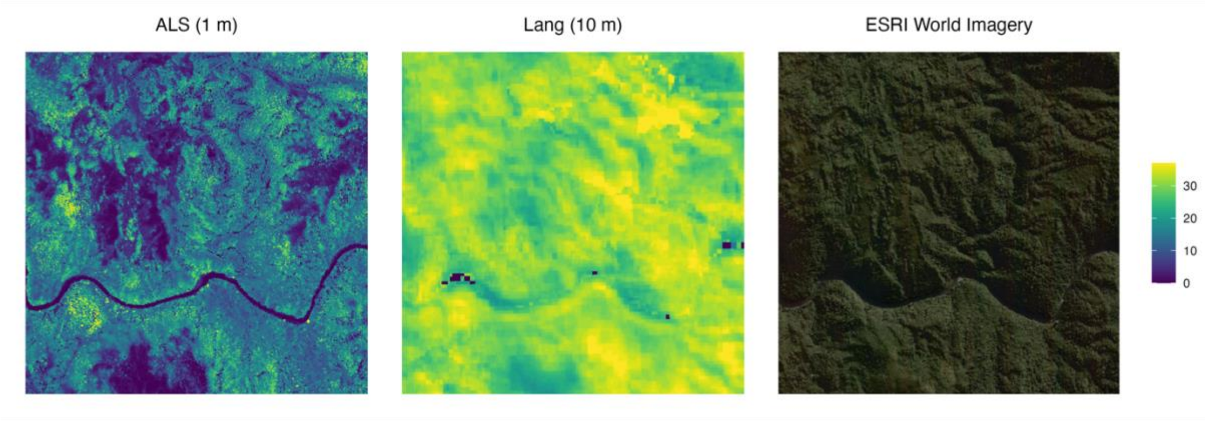

**Figure S1.5: Site with largest absolute errors for Lang model.** Same as Fig. 3.4, but with the largest absolute errors for the Lang CHM. The site is Mount Fincham in Tasmania. The mean height in the ALS model across the entire site is 11.1 m, while Lang predicts 27.6 m.

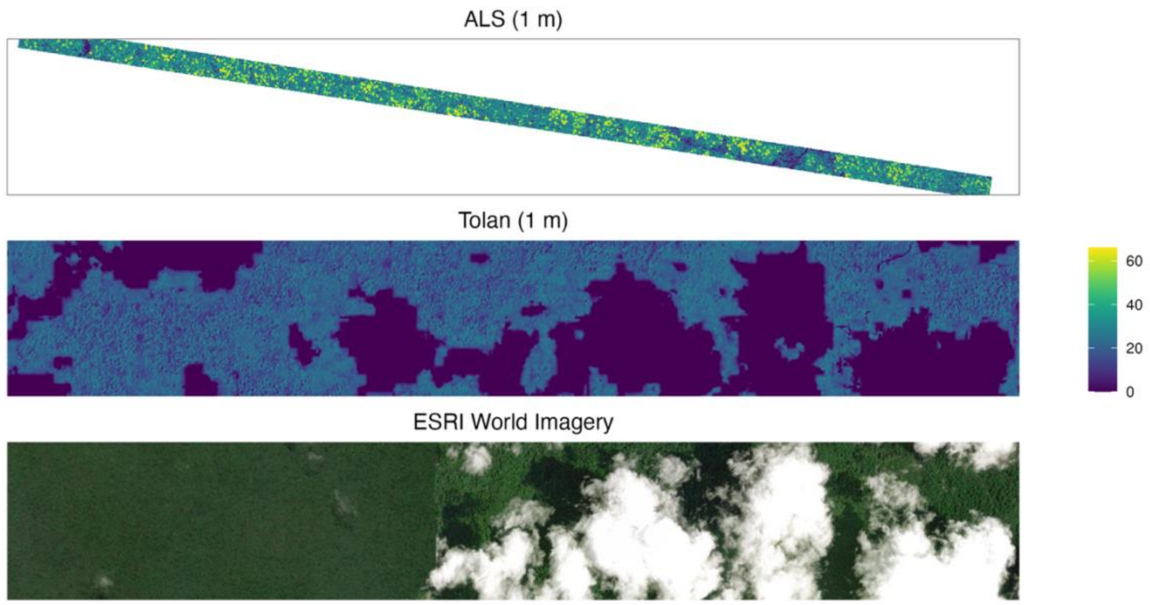

**Figure S1.6: Site with largest absolute errors for Tolan model.** Same as Figs. 1.4 and 1.5, but with the largest absolute errors for the Tolan CHM. The site is near Rio Jari in Brazil. The mean height in the ALS model across the entire transect is 33.1 m, while the Tolan model predicts 8.0 m for the same area. For visualization purposes, we also show areas surrounding the transect. Note that there is a general bias in the Tolan model at this site and that this is further exacerbated by visible cloud artefacts. The clear correspondence of cloud outlines in ESRI World Imagery data with gaps in the righthand part of the image suggest that part of this image was very likely used for predictions with the Tolan model, leading to substantial cloud artefacts.

### Case Study #2

##### S2.1 Data filtering and voronoi tessellation for gridding into 1 $km^2$ cells

To create comparable units of analysis, we removed a 50 m buffer from all CHMs in the GCA and divided them into cells of 1 $km^2$. Instead of a fixed rectangular grid of 1 $km^2$ cells, we used an iterative procedure that creates a Voronoi tessellation with a cell size of 1 $km^2$ for each scan (Fig. S2.1). The initial Voronoi tessellation is derived via k-mean clustering so that the mean of all cell sizes is 1 $km^2$, with some variation around the mean. To minimize this variation, we iteratively moved the initial Voronoi seed points by a small random distance and accepted new locations only if they reduced the cell size coefficient of variation (CV). This was repeated until either the CV was < 0.01 or 5,000 iterations were reached. An example of the resulting Voronoi grid is shown in Fig. S2.1. The utilization of Voronoi cells allowed us to include a wide range of scans (e.g., including scan lines with a width < 1 km) and maximized the area that could be used for forest structure analysis (e.g., doubling the total usable extent in Fig. S2.1). We note that highly elongated Voronoi cells could bias results due to increased edge effects along cell borders (e.g., splitting of canopy gaps), so we filtered out cells with a perimeter > 8 km (twice the perimeter of a square grid cell with an area of 1 $km^2$). Further, we only retained cells where at least 99.9% of the terrain was stably inferred (assessed via mask_unstabledtm.tif), with more than 70% canopy cover at 2 m, less than 1% of Landsat-observed surface water (Potapov et al., 2022), and with a MODIS EVI > 80% of peak EVI during scan acquisition (Didan, 2025). To avoid duplication, for sites with repeat scans, we selected only cells from the last scan date. We also excluded any cells with fewer than 20 canopy gaps.

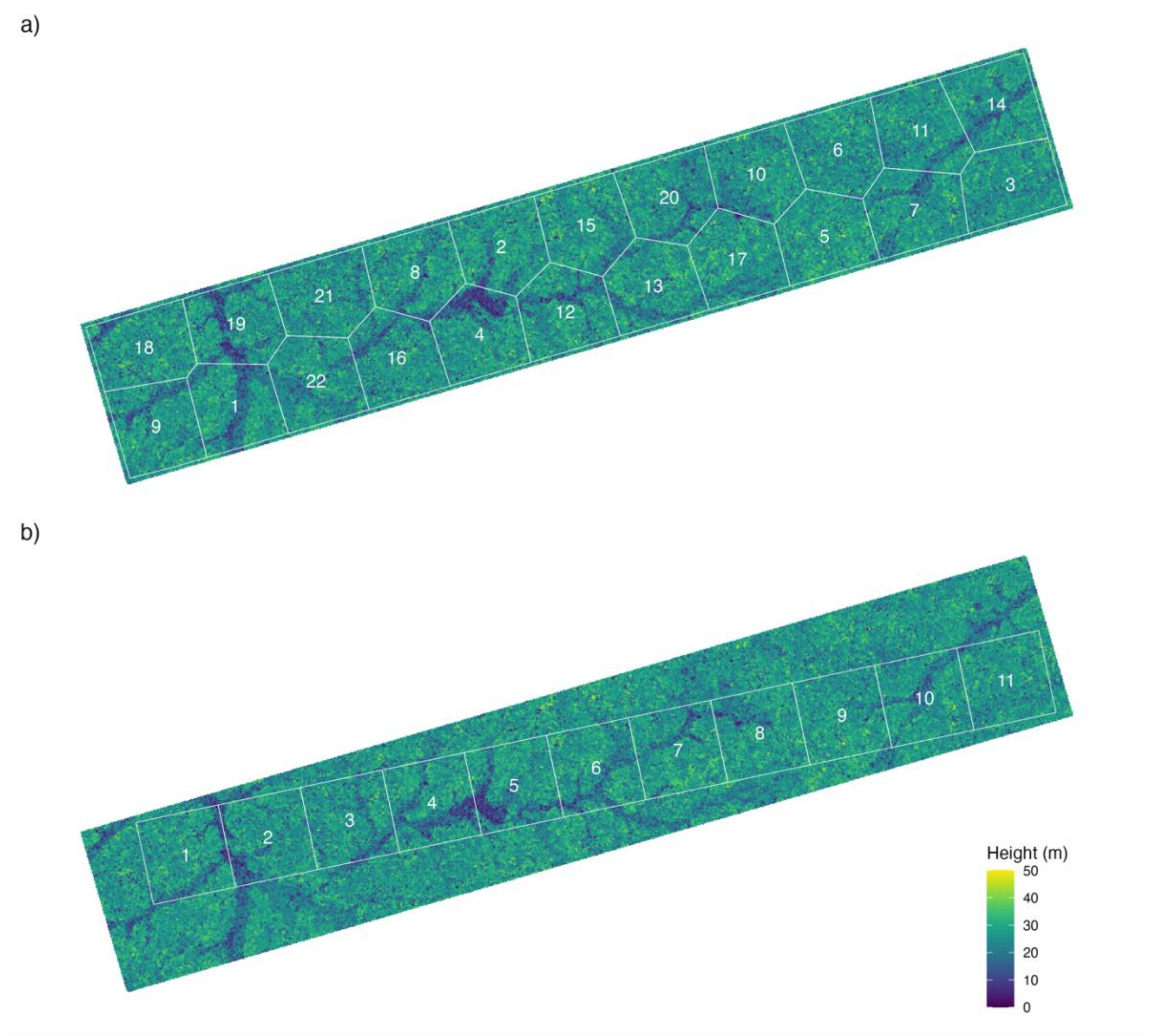

**Figure S2.1: Example of Voronoi tessellation with 1 km² grid cells.** Shown is the canopy height model of a tropical rain forest scan over ~22 km² in the Democratic Republic of Congo. Overlaid are a) a Voronoi tessellation where each cell has an area of 1 km², and b) a corresponding rectangular grid with 1 km² grid size, which was randomly rotated until maximizing the number of cells (11). The maximum number of rectangular grid cells stands in stark contrast to the corresponding Voronoi grid which can allocate twice as many cells over the same area (22). Note that in both cases, a 50 m boundary layer was cut off the scan extent to remove edge effects.

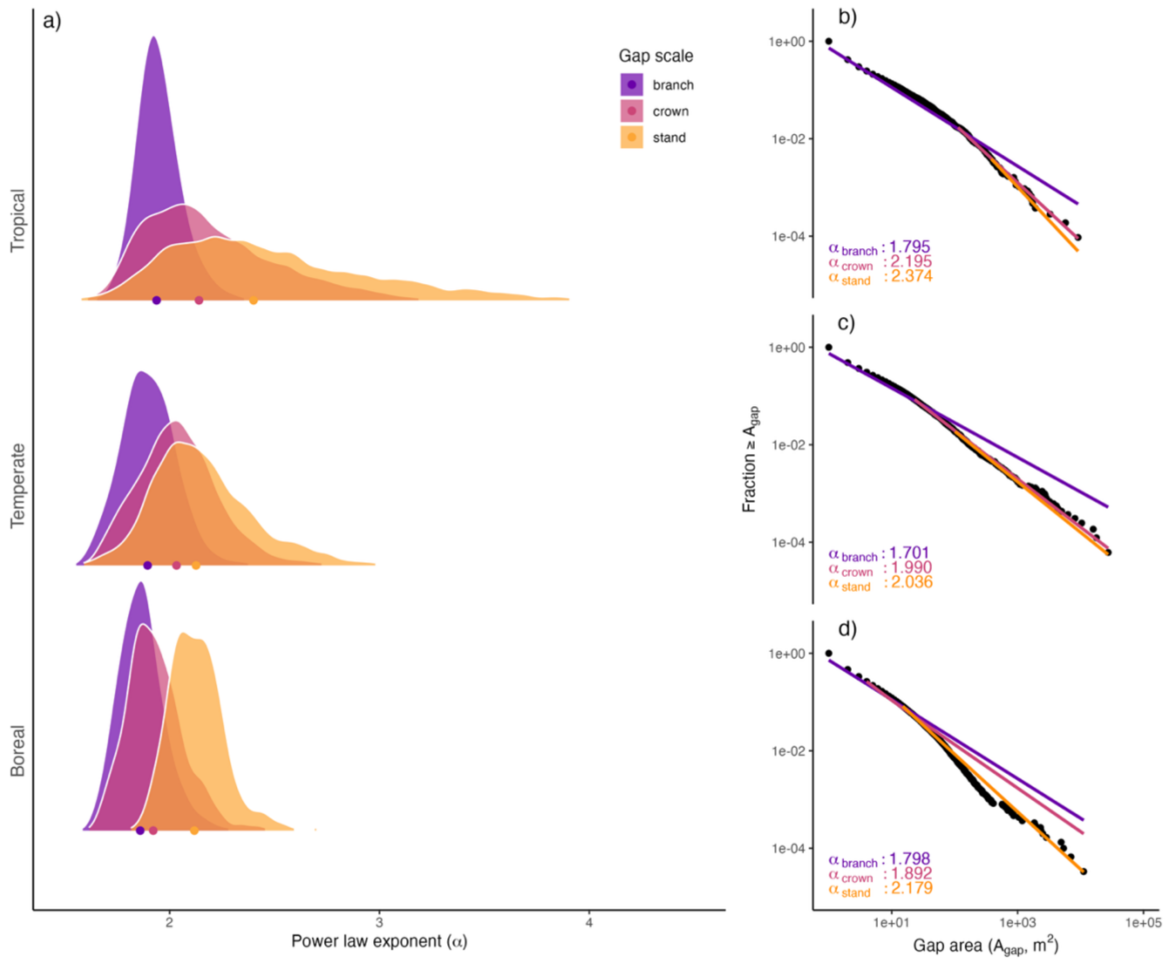

**Figure S2.2: Deviations from power law scaling across major biomes (TIN).** This is exactly the same figure as Fig. 3 in the main text, but with all gap statistics calculated from a TIN-based canopy height model (triangulated irregular network of first returns, or a "light interception height" model) instead of the default locally adaptive spikefree canopy height model ("canopy top height" model). The choice of CHM algorithm strongly affects the size and number of recorded gaps (Fischer et al., 2024), so we also expected a strong effect on the power law coefficient $\alpha$. Note how $\alpha$ here depends less on the range of gap sizes over which it is calculated than in Fig. 3, but that the same qualitative shifts exist, with $\alpha$ increasing when the calculation is limited to larger gaps, and substantial variation in $\alpha$ at the crown and stand scale, particularly in tropical forests.

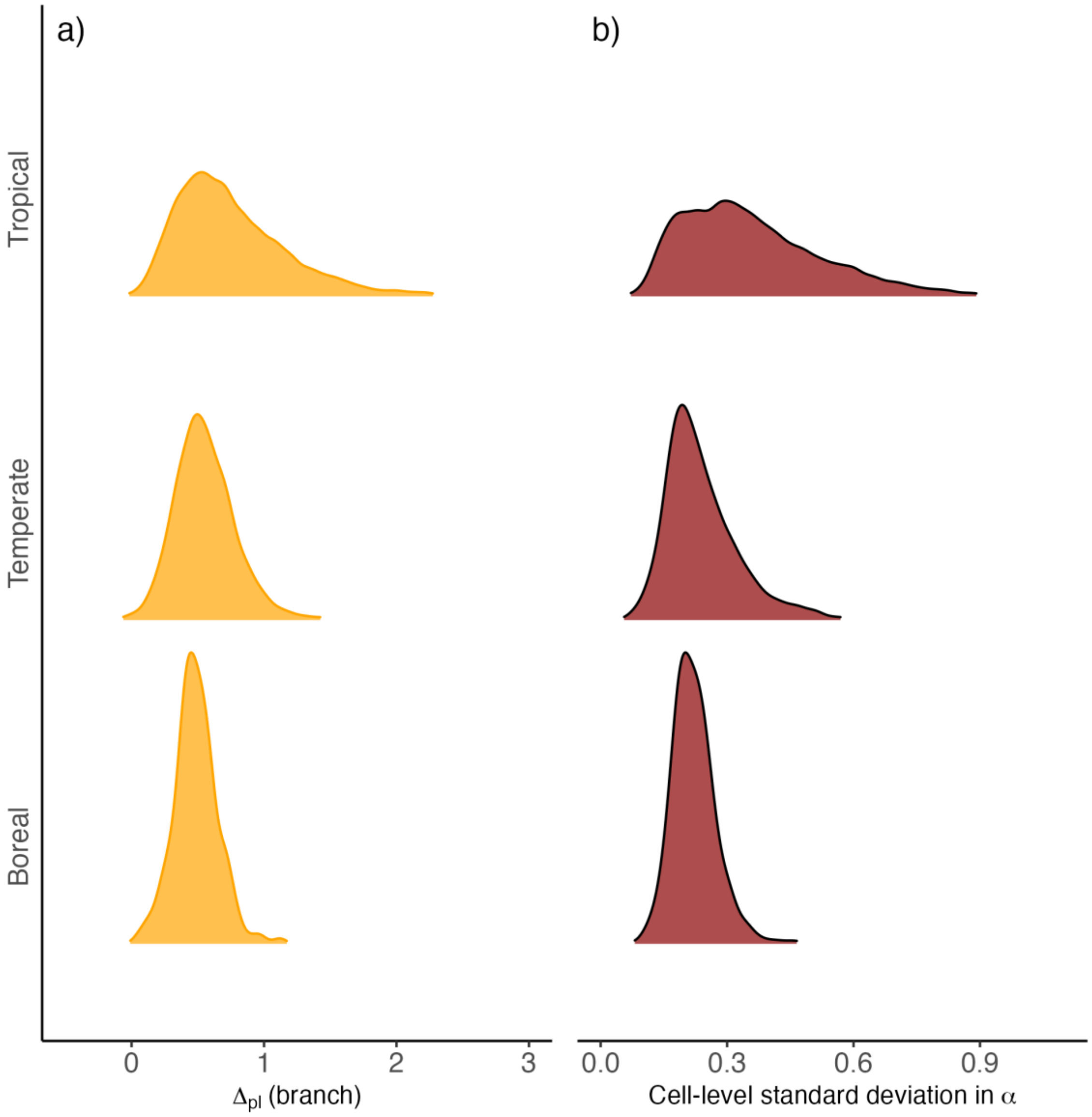

**Figure S2.3: Deviations from power law scaling across biomes and sites.** Shown are the calculated deviations from power law scaling (panel a) between $\alpha_{branch}$ and $\alpha_{crown}$, split by biome, as well as an estimate of how much $\alpha$ varies within each cell depending on scale and measurement methodology (panel b). The former shows that the average difference between $\alpha$ calculated across all gap sizes ($\alpha_{branch}$) and $\alpha$ calculated only for canopy treefalls or larger ($\alpha_{crown}$) is at least 0.5 across biomes. The latter shows that, for a single 1 km$^2$ cell, the average standard deviation of $\alpha$ is least 0.25 across all the methodologies employed in this study.

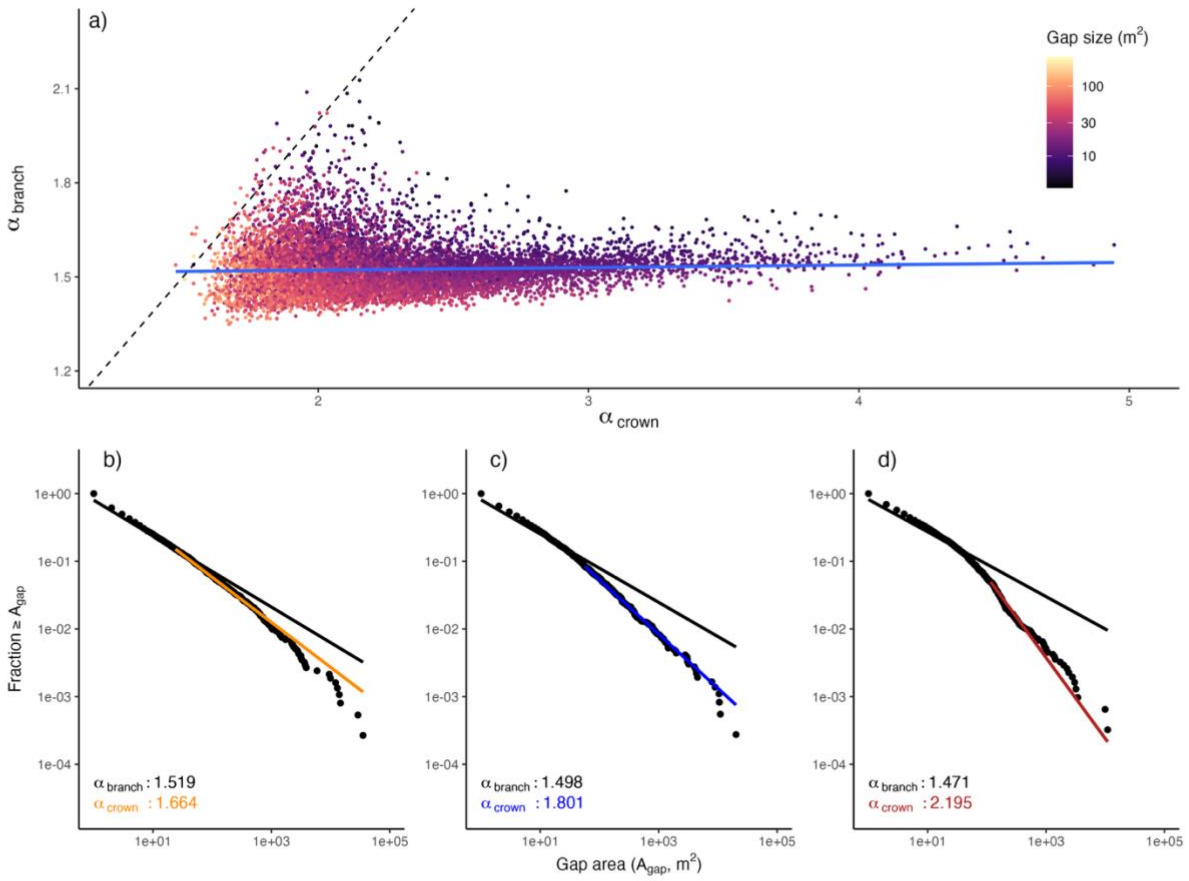

**Figure S2.4: Correlation between $\alpha_{branch}$ and $\alpha_{crown}$ (spikefree).** Shown is the correlation between $\alpha_{branch}$ and $\alpha_{crown}$ when calculated from a spikefree canopy height model (panel a), with the 1:1 line shown as dashed line, and coloured by gap size. Each point corresponds to a single 1 km$^2$ cell. The blue line represents an OLS regression line. Under true power law scaling, $\alpha_{branch}$ and $\alpha_{crown}$ should be nearly the same and thus tightly correlated. Here, there is next to no such relationship. Panels b, c, and d show the gap size frequency distributions (complementary cumulative distribution function) and power law fits for three 1 km$^2$ tiles at a single site in Danum Valley, Borneo. Tiles were chosen to illustrate the decoupling of $\alpha_{branch}$ and $\alpha_{crown}$ at a single site.

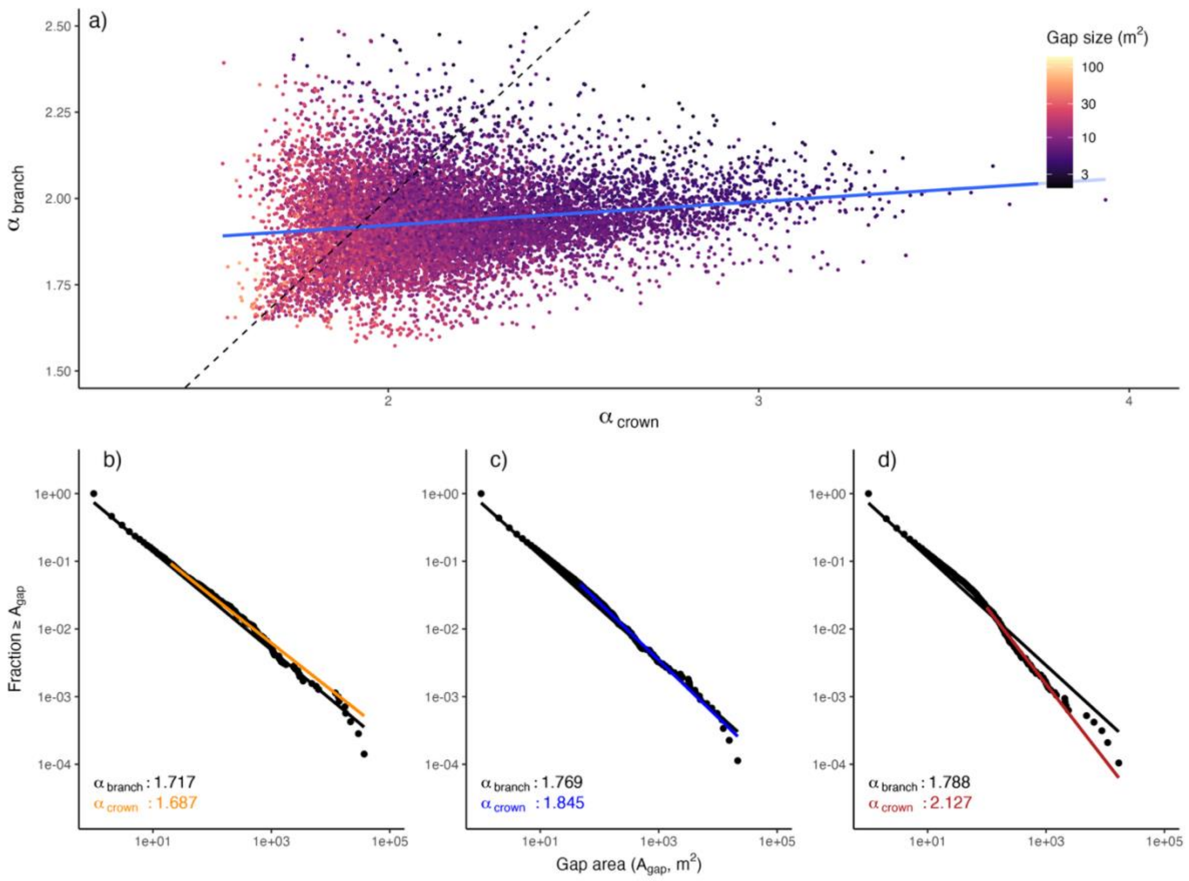

**Figure S2.5: Correlation between $\alpha_{branch}$ and $\alpha_{crown}$ (TIN).** Same as Fig. S2.4, but based on the TIN canopy height model (Delaunay triangulation of first returns). Under true power law scaling, $\alpha_{branch}$ and $\alpha_{crown}$ should be nearly the same and thus tightly correlated. Here, a lot more sites lie around the 1:1 line than in Fig. S2.4, indicating that power law behaviour may exist in some cases, but the figure shows that there is still little overall correlation.

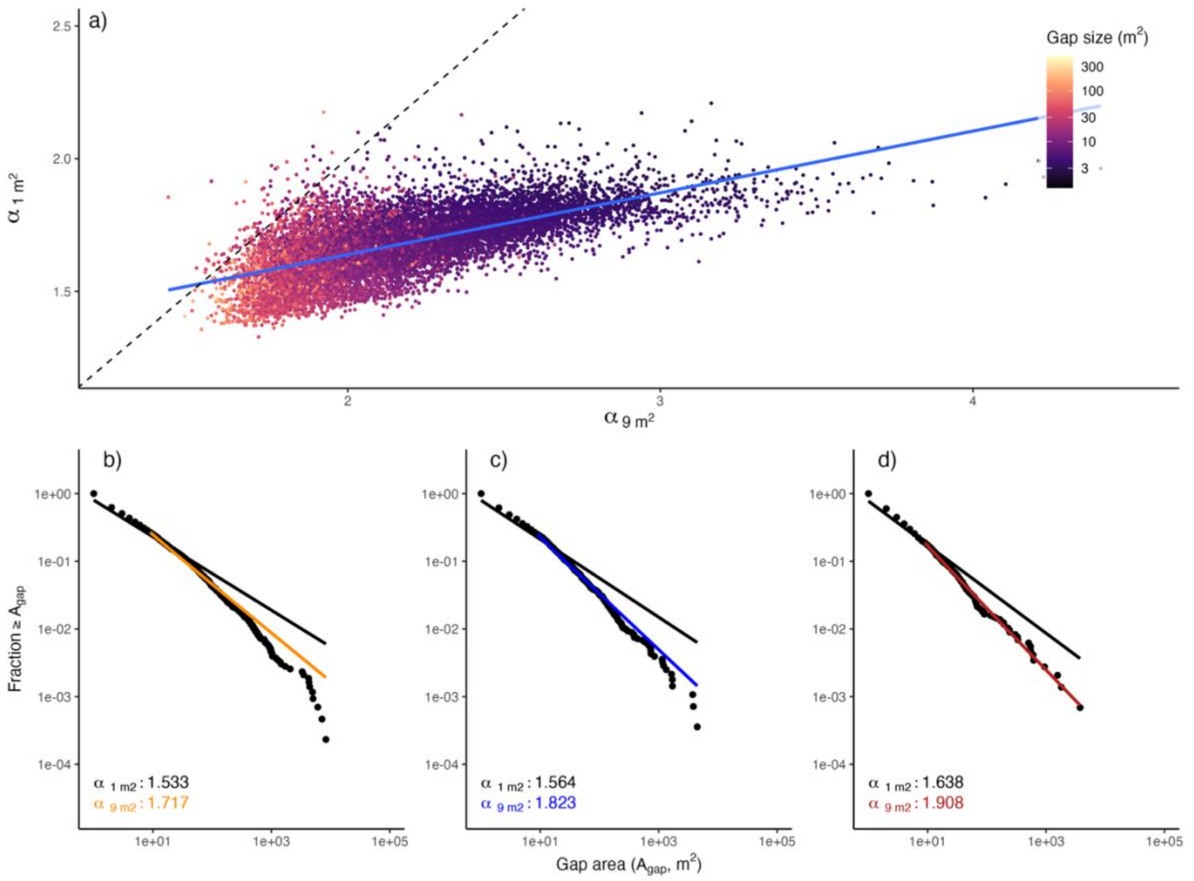

**Figure S2.6: Correlation between $\alpha$ values for gaps < 2 m in canopy height (spikefree).** Same as Fig. S2.4, but showing the correlation between $\alpha$ calculated for gaps >= 1 $m^2$ and $\alpha$ calculated for gaps >= 9 $m^2$, based on the spikefree canopy height model (panel a). Note how the inferred power law exponents change massively just by removing very small gaps (< 9 $m^2$), and that barely any points lie on the 1:1 line.

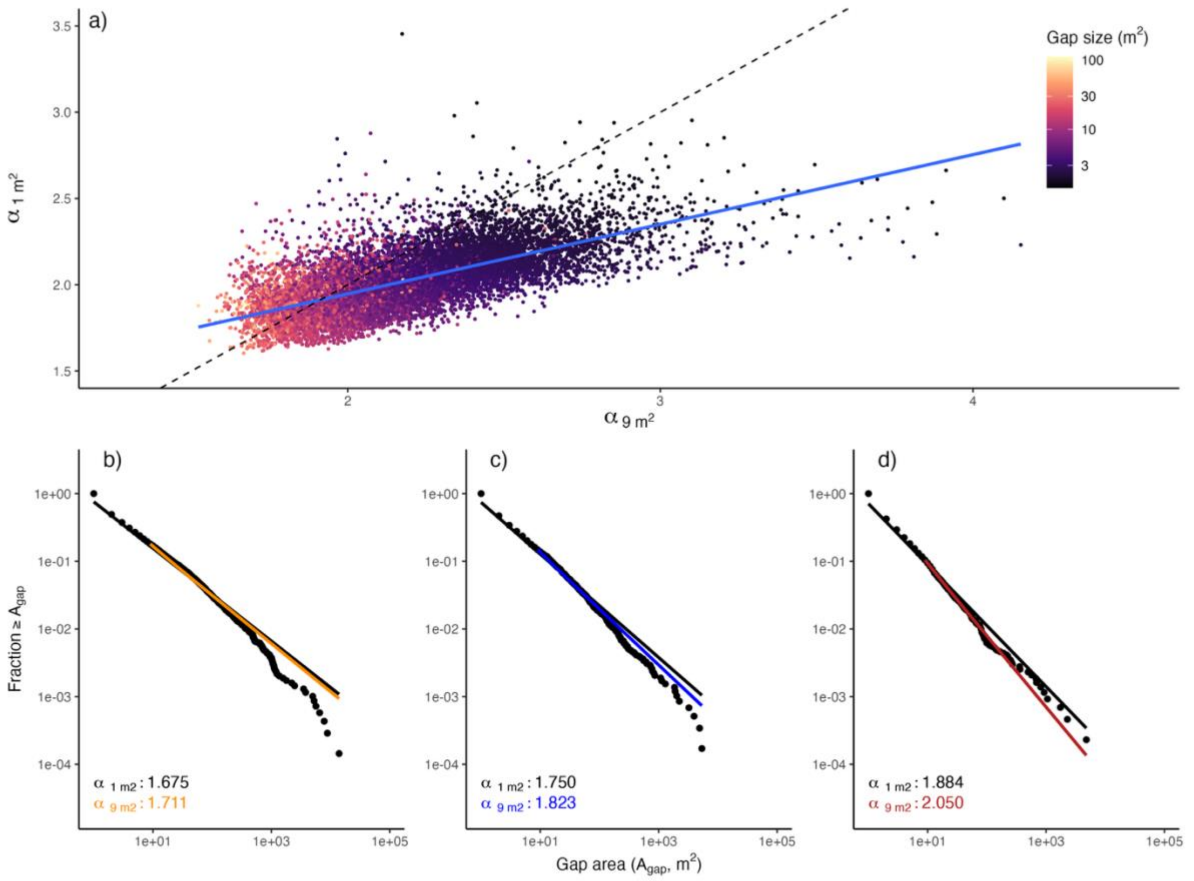

**Figure S2.7: Correlation between α values for gaps < 2 m in canopy height (TIN).** Same as Fig. S2.6, but with the TIN canopy height model. In this case, power law fits are slightly more stable, but note that there is still a clear deviation visible in panel a) and that this deviation is only due to removing the very smallest gaps (< 9 $m^2$ in size).

### Case Study #3

**S3.1 Inferring change rates from observed disturbances over varying scan intervals**

We assume a forest landscape where all non-forest area has been masked out and each pixel within forests is classified as either undisturbed canopy (set to 1) or disturbed gap area (set to 0). Without loss of generality, we will assume in the following that this classification is based on canopy height models, as in the main text (all areas < 10 m are gaps). Based on the binary disturbance classification, we can define a standardized disturbance rate *d* and standardized recovery rate *r*, which are the proportion of pixels that switch from 1 to 0 or 0 to 1, respectively, over a reference time period $t_{ref}$. The reference period $t_{ref}$ is arbitrary, but we here choose $t_{ref}$ = 1 yr, as this simplifies distinguishing periodic canopy dynamics (e.g., leaf shedding/flush during summer-winter or wet-dry season transitions) from discrete disturbance events.

To infer *d* and *r*, we need to calculate the observed proportion of pixels that switch from 1 to 0 (observed disturbance *D*) and from 0 to 1 (observed recovery *R*) over a time interval *t* between two observations. In cases where $t = t_{ref}$ and assuming no errors in the observations, we can trivially infer $d = D$ and $r = R$. However, in most cases, instrumentation errors exist, and observation intervals t are larger than tref. In such cases, *d* is not just $D / t$. For example, if we interpret d as probability for any pixel to switch from 1 to 0 over time period tref, then for any period $t > t_{ref}$, the pixel has a non-zero probability of switching twice, i.e., from 1 to 0 and back from 0 to 1. The initial disturbance event will thus be transient and invisible in D. Therefore $D / t < d$ for any $t > t_{ref}$. The same holds for transient recovery ($R / t < r$). In the following, we will derive a function to translate between *D* and *R* and *d* and *r* for any given time *t*.

To arrive at an analytical solution, we start with a simple recursion formula that interprets the probability of a pixel to be canopy at time $t + 1$ ($C_{t+1}$) as a function of the same probability at time $t$ ($C_t$). $C_{t+1}$ is the combined probability of staying in the undisturbed state $C_t$ with disturbance probability $(1 – d)$ or switching from the complementary gap state $(1 – C_t)$ with the recovery probability $r$:

$$C_{t+1} = C_t \times (1 - d) + (1 - C_t) \times\ r = C_t \times\ (1 - d - r) + r \tag{1}$$

This difference equation can be solved as follows:

$$C(t)\ =\ \frac{r}{d + r} + \frac{d}{d + r}\ \times\ (1 - d - r)^t \tag{2}$$

From this, we can derive the complementary probability that a pixel switches to non-canopy over period $t$:

$$D(t) = 1 - C(t) = \frac{d}{d + r}\ \times (1 -\ (1 - d - r)^t) \tag{3}$$

However, in most situations, $D(t)$ also includes noise or biases, such as small movements of branches between two scans, geolocation errors due to varying scan angles and flightlines, or uncertainty in ground reconstruction. These can lead to an arbitrary switching between canopy (1) and gap (0) state, particularly in pixels at the edge between canopy areas and gap areas. This leads to a positive offset in $D(t)$ which we capture through a constant $d_0$. So:

$$D(t) = d_0 + \frac{d}{d + r} \times (1 -\ (1 - d - r)^t) \tag{4}$$

Since $D(t = 0) = d_0$, we can also interpret the intercept term $d_0$ as the “disturbance” bias or artefact that would be observed between two temporally coincident scans.

Using Equation 4, we can also calculate the probability $D_{tot}$ that a pixel has been classified as disturbed at any point up to time $t$, ignoring its potential recovery. This is equivalent to setting $r$ = 0 (no transient disturbances):

$$D_{tot}\,(t) = d_0 + \,(1 - \,(1 - d)^t) \tag{5}$$

Equation 5 can also be interpreted as the sum of the true disturbance probability $D_{true}$ ($t$) = 1 – (1 – $d$)$^t$ and the background probability $d_0$ that accounts for instrument noise. The difference $D_{miss}$ ($t$) = $D_{tot}$ ($t$) - $D$($t$) is independent of $d_0$ and quantifies the extent of undetected disturbance as a function of time t.

In analogy to Equation 4, we can also derive a similar equation for recovery, namely:

$$R(t) = r_0 + \frac{r}{d + r} \times (1 - \,(1 - d - r)^t) \tag{6}$$

Again, $R$($t$ = 0) = $r_0$, so $r_0$ can be interpreted as the “recovery” artefact that would be seen when comparing two temporally coincident scans. Note that $r_0$ should be more uncertain than $d_0$, as, by definition, most closed-canopy forests will have less gap area than canopy area, but when converted to a fixed-area estimate and estimated for large areas $r_0$ ~ $d_0$.

**S3.2 Modelling observed rates of change**

Equation 4 describes the relationship between two empirical quantities $D$ and $t$ as a function of three biologically interpretable parameters $d_0$, $d$, and $r$. By fitting the equation to empirical data on $D$ and $t$, we can infer all three parameters. For practical purposes, we simplify the equation via reparameterization. We define:

$$\tau = -\log\,(1 - d - r) \tag{7}$$

and

$$\kappa = \frac{d}{d + r} \tag{8}$$

and obtain:

$$D(t) = d_0 + \kappa \times (1 - e^{-\tau t}) \tag{9}$$

For all following analysis here and in the main text, we chose a Bayesian approach to fit Equation 7 to the NEON dataset of 28 scan pairs, relying on the R package *brms* (Bürkner, 2018), which wraps the STAN software (Carpenter et al., 2017). We chose weakly informative priors for $d_0$ and $\kappa$, namely a beta distribution with shape parameters 1 and 5 (left skewed, bounded between 0 and 1), and also for $\tau$, namely a gamma distribution with parameters 2 and 10 (also left skewed, without upper bounds). We summarized posteriors for all parameters through median values and, using $d + r = 1 - e^{-\tau}$, converted them back into estimates of change rates:

$$d = \kappa \times (1 - e^{-\tau}) \tag{10}$$

and:

$$r = (1 - \kappa) \times (1 - e^{-\tau}) \tag{11}$$

Both *d* and *r* were then converted back from proportions of gap and disturbed area (%) into units of ha $km^{-2}$. When fitting the models, we carried out standard posterior predictive checks and various robustness tests (cf. Section 3.4). We note that the entire procedure could also have been carried out with Equation 6, for example, or a combination of Equation 4 and 6 (multivariate response), and that the modelling framework does not presuppose steady-state or equilibrium canopy dynamics. However, in our analysis, we introduce the implicit assumption that disturbance and recovery rates have remained approximately constant between 2012 and 2024. This makes it possible to more robustly quantify noise and errors, but is not generally necessary to apply the modelling framework (cf. Section 3.3).

**S3.3 Alternative method to obtain estimates of change rates from single pairs of scans**

In S3.2, we presented an approach to estimate steady-state canopy dynamics by fitting a non-linear function to observed change rates over varying time intervals. However, this requires a large number of scans. To assess how well we could estimate change rates from single pairs of scans, we also present an alternative approach, whereby one can derive direct estimates of *d* and *t* from a single pair of scans.

When *D*, *R*, and *t* are known and assuming $d_0 = r_0 = 0$ (or at least near 0), then Equations 4 and 6 have only two unknown parameters *d* and *r* and thus can be solved analytically. Setting $\lambda = (d + r)$, it holds

$$D + R = (1 - (1 - \lambda)^t) \tag{12}$$

Or:

$$\lambda = 1 - (1 - D - R)^{1/t} \tag{13}$$

And using Equations 3 and 11, we derive:

$$d = \frac{D}{(1 - (1 - \lambda)^t)} \times \lambda = \frac{D}{D + R} \times \lambda \tag{14}$$

And:

$$r = \frac{R}{(1 - (1 - \lambda)^t)} \times \lambda = \frac{R}{D + R} \times \lambda \tag{15}$$

These formulas only hold when $d_0 \sim 0$ and $r_0 \sim 0$, but this is unlikely to be the case. However, we can use empirical estimates of D (t = 0) = $D_0$ and R(t = 0) = $R_0$ from coincident scans (e.g., as in 3.4) to derive a corrected $D^* = D - D_0$ and $R^* = R - R_0$. For $D^*$ and $R^*$ we can calculate $d^* = \frac{D^*}{D^*+R^*} \times \lambda^*$ and $r^* = \frac{R^*}{D^*+R^*} \times \lambda^*$ directly from single pairs of scans.

**S3.4 Robustness checks**

We carried out four additional robustness checks of our results. (1) We repeated the model fitting with disturbances greater or equal 25 $m^2$ in size. We expected that this would considerably reduce the inferred value of $d_0$ (less instrument-dependent noise), with only a small effect on estimates of *d* and *r*. (2) We repeated the model fitting on an 8 $km^2$ subset of the area where the full set of NEON scans is available (8 NEON scans, 3 GLiHT scans, 1 3DEP scan), but excluded an additional scan acquired by 3DEP, as it was carried out in winter. Since the subset only covers ~5% of the total area, we expected that *d* and *r* would differ, but remain in a similar range as previous estimates. (3) We used the pairwise estimation approach from Section S3.3 with all 28 NEON scan pairs across the full area to derive an estimate of *d* and *r* for each scan pair and then derived average rate estimates. To account for random noise between scans, we used rough estimates of $d_0$ and $r_0$. Based on both the non-linear models in Section 3.2 and empirical data (cf. following point 4), we inferred these values to be ~1% of intact canopy area and ~10% of gap area (or ~1 ha $km^{-2}$ for each). We expected average values close to previous estimates, but with considerable noise around the mean. 4) Using the 8 $km^2$ subset, we derived an empirical estimate of $d_0$ by calculating the proportion of pixels that switched from 1 to 0 between GLiHT 2012 and NEON 2012, and vice versa, and between GLiHT 2017 and NEON 2017, and vice versa (n = 4 x 8 = 32 cells of 1 $km^2$). We expected this to be close to the values of $d_0$ and $r_0$ that were estimated as intercepts of a non-linear model with a fully independent data set.

| Dataset | Citation | Coordinate Reference System | Period | System | Area (km$^{2}$) | Pulse density (m$^{-2}$) |
|---|---|---|---|---|---|---|
| 3DEP | https://rockyweb.usgs.gov/vdelivery/Datasets/Staged/Elevation/LPC/Projects/USGS_LPC_MA_NE_CMGP_Sandy_Z18_2013 | NAD83(2011) / UTM zone 18N (EPSG:6347) | 16/11/2013 - 16/04/2014 | | 356.9 | 3.53 |
| G-LiHT-US | Cook et al. 2013. NASA Goddard's Lidar Hyperspectral and Thermal (G-LiHT) airborne imager. Remote Sensing 5:4045-4066 doi:10.3390/rs5084045. | WGS 84 / UTM zone 18N (EPSG:32618) | 19/06/2012 - 21/06/2012 | Riegl VQ-480 | 19.0 | 17.21 |
| G-LiHT-US | Cook et al. 2013. NASA Goddard's Lidar Hyperspectral and Thermal (G-LiHT) airborne imager. Remote Sensing 5:4045-4066 doi:10.3390/rs5084045. | WGS 84 / UTM zone 18N (EPSG:32618) | 09/08/2017 - 19/08/2017 | Riegl VQ-480i | 13.3 | 26.56 |
| G-LiHT-US | Cook et al. 2013. NASA Goddard's Lidar Hyperspectral and Thermal (G-LiHT) airborne imager. Remote Sensing 5:4045-4066 doi:10.3390/rs5084045. | WGS 84 / UTM zone 18N (EPSG:32618) | 06/08/2021 - 06/08/2021 | Riegl VQ-480i | 11.5 | 26.99 |
| NEON (proto) | | WGS 84 / UTM zone 18N (EPSG:32618) | 14/08/2012 - 14/08/2012 | Optech ALTM Gemini | 172.0 | 2.26 |
| NEON | NEON (National Ecological Observatory Network). Discrete return LiDAR point cloud (DP1.30003.001). https://data.neonscience.org (last accessed July 20 2023) | WGS 84 / UTM zone 18N (EPSG:32618) | 29/05/2014 - 02/06/2014 | Optech ALTM Gemini | 281.2 | 3.52 |
| NEON | NEON (National Ecological Observatory Network). Discrete return LiDAR point cloud (DP1.30003.001). https://data.neonscience.org (last accessed July 20 2023) | WGS 84 / UTM zone 18N (EPSG:32618) | 16/08/2016 - 28/08/2016 | Optech ALTM Gemini | 336.7 | 4.20 |
| NEON | NEON (National Ecological Observatory Network). Discrete return LiDAR point cloud (DP1.30003.001). https://data.neonscience.org (last accessed July 20 2023) | WGS 84 / UTM zone 18N (EPSG:32618) | 13/08/2017 - 23/08/2017 | Optech ALTM Gemini | 294.2 | 3.97 |
| NEON | NEON (National Ecological Observatory Network). Discrete return LiDAR point cloud (DP1.30003.001). https://data.neonscience.org (last accessed July 20 2023) | WGS 84 / UTM zone 18N (EPSG:32618) | 28/08/2018 - 05/09/2018 | Optech ALTM Gemini | 295.5 | 4.39 |
| NEON | NEON (National Ecological Observatory Network). Discrete return LiDAR point cloud (DP1.30003.001). https://data.neonscience.org (last accessed July 20 2023) | WGS 84 / UTM zone 18N (EPSG:32618) | 11/08/2019 - 26/08/2019 | Optech ALTM Gemini | 304.3 | 4.34 |
| NEON | NEON (National Ecological Observatory Network). Discrete return LiDAR point cloud (DP1.30003.001). https://data.neonscience.org (last accessed February 2 2024) | WGS 84 / UTM zone 18N (EPSG:32618) | 03/08/2022 - 14/08/2022 | Optech ALTM Gemini | 313.1 | 6.93 |
| NEON | NEON (National Ecological Observatory Network). Discrete return LiDAR point cloud (DP1.30003.001). https://data.neonscience.org (last accessed March 25 2025) | WGS 84 / UTM zone 18N (EPSG:32618) | 14/08/2024 – 04/09/2024 | Optech Galaxy Prime | 300.0 | 28.9 |

**Table S3.1: Airborne laser scans at Harvard Forest.** This table shows the acquisition characteristics of airborne laser scans at Harvard Forest.

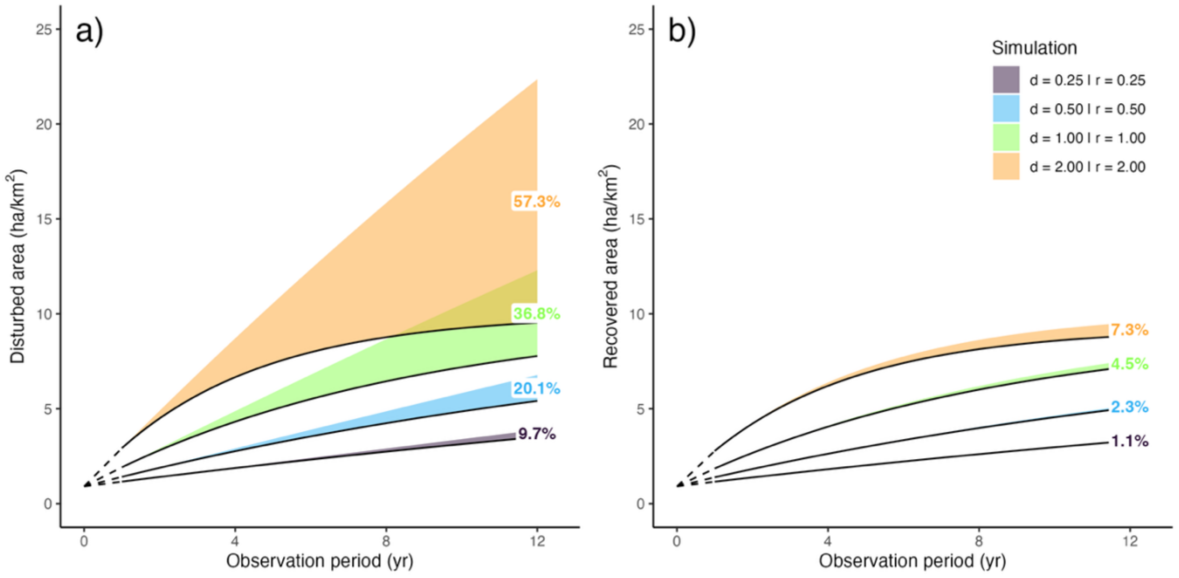

**Figure S3.1: Scenarios of short-term dynamics and the contribution of transient dynamics.** Shown are simulated disturbance and recovery trajectories over 12 years for a forest landscape with 90% forest cover and 10% gap area, i.e. mirroring the structure and temporal coverage at Harvard Forest in the main text (Fig. 4). All scenarios shown describe steady-state dynamics, i.e. equal disturbance and recovery rates, but at varying intensities, from low-turnover dynamics (disturbance and recovery rates of 0.25 ha $km^{-2}$ $yr^{-1}$, or 0.25%) to high-turnover dynamics (2 ha $km^{-2}$ $yr^{-1}$, or 2%). Predictions are based on the analytical equations in S3.1. The upper border of the colour-filled areas shows the true disturbance and recovery rate for each time interval, the black border at the lower end what would be observed between two observations over this interval. The percentages indicate the amount of transient (and thus potentially unobserved) dynamics for a 12 year-interval between two scans. For slowly changing forest landscapes – low disturbance rates and low recovery rates –, the amount of unobserved dynamics over long time intervals tends to be small (< 10% for the lowest rate). In stark contrast, it explodes for forest landscapes with high turnover. I.e., under a 2% turnover rate, nearly 60% of the area disturbed within 12 years is expected to close again within the 12-year period.

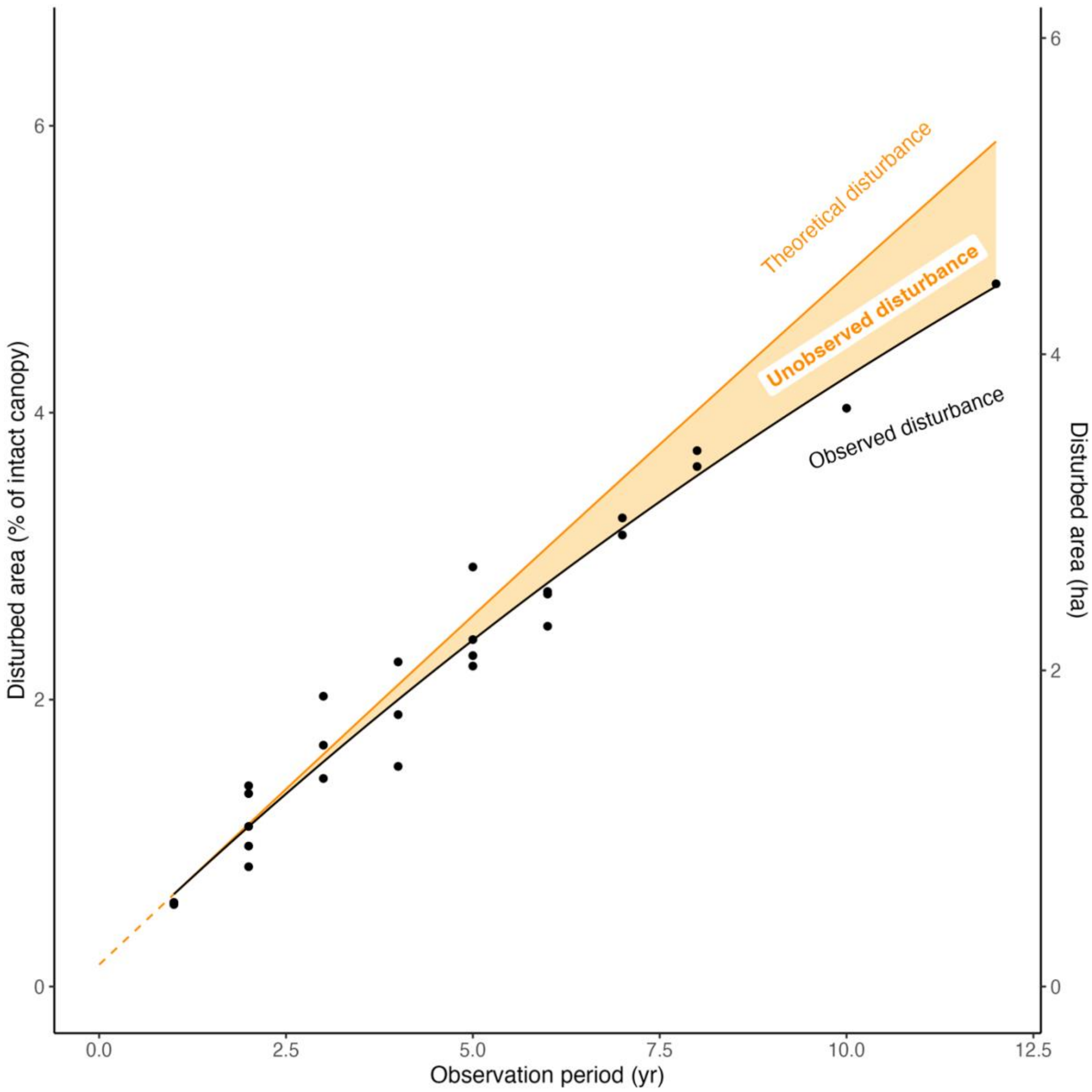

**Figure S3.2: Disturbance as function of time (only gaps ≥ 25 m²).** This mirrors Fig. 4a in the main text, but instead of showing disturbed area for any disturbance down to 1 m², restricts the disturbed area to a minimum of 25 m². The inferred 1-year disturbance rate d = 0.45 ha km$^{-2}$ is identical to the inferred 1-year disturbance rate from all disturbances, but, as expected, the acquisition bias (intercept term) is now vanishingly small ($d_0$ = 0.14 ha km$^{-2}$). This indicates that subsetting to larger gaps effectively removed noise between acquisitions. However, we note that subsetting to gaps of 25 m² or more lowers the estimated 1-year recovery rate (r = 0.34 ha km$^{-2}$ instead of 0.41 ha km$^{-2}$) and the estimated amount of transient dynamics after 12 years (18% instead of 21%). Since gap recovery (lateral ingrowth of branches, gradual ingrowth of regenerating trees from below) occurs in a more spatially disperse manner than gap creation (treefall), the introduction of minimum gap sizes may thus bias the inference of recovery rates.

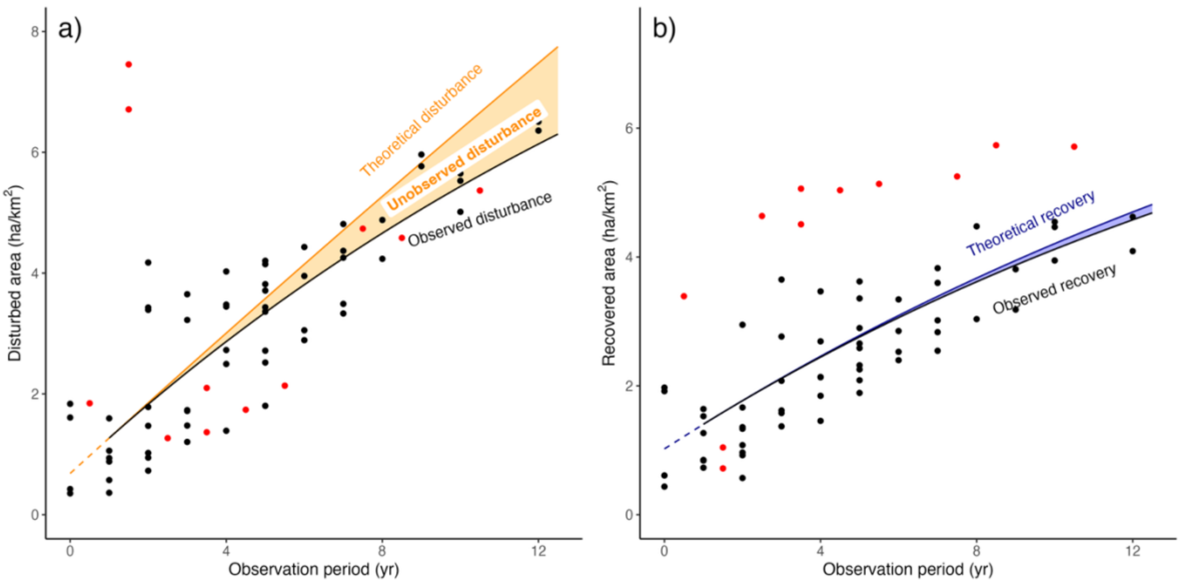

**Figure S3.3: Short-term dynamics at Harvard Forest (8 km² GLiHT subset).** This figure replicates panels a) and b) in Fig. 4 in the main text, but for a subset of the total scan area that is also covered by 3 GLiHT airborne laser scans (2012, 2017, 2021). This allows for a much larger number of comparisons across time, although the local dynamics may not fully represent dynamics across the wider Harvard Forest area. As in Fig. 4, each black dot represents a combination of scans. Red dots show additional scan combinations that involve a 3DEP scan in winter 2013/2014, but were not included in model fitting due to the confusion of leaf phenology effects ("leaf-off" condition) with structural change. The resulting biases can be seen in the positive "disturbance" outliers from 2012 to 2013/14 (panel a), and the general positive "recovery" outliers with respect to the 2013/2014 scan (panel b). Overall, 1-year disturbance rates are higher over the 8 km² than for the entire NEON area (d = 0.58 ha km$^{-2}$), while recovery rates (r = 0.38 ha km$^{-2}$) and the proportion of 12-year transient dynamics (19%) are slightly lower. Note also how laser scanning noise estimates are on the same order of magnitude as in Fig. 4, with a lower $d_0$ = 0.68 ha km$^{-2}$, but a higher $r_0$ = 1.02 ha km$^{-2}$.

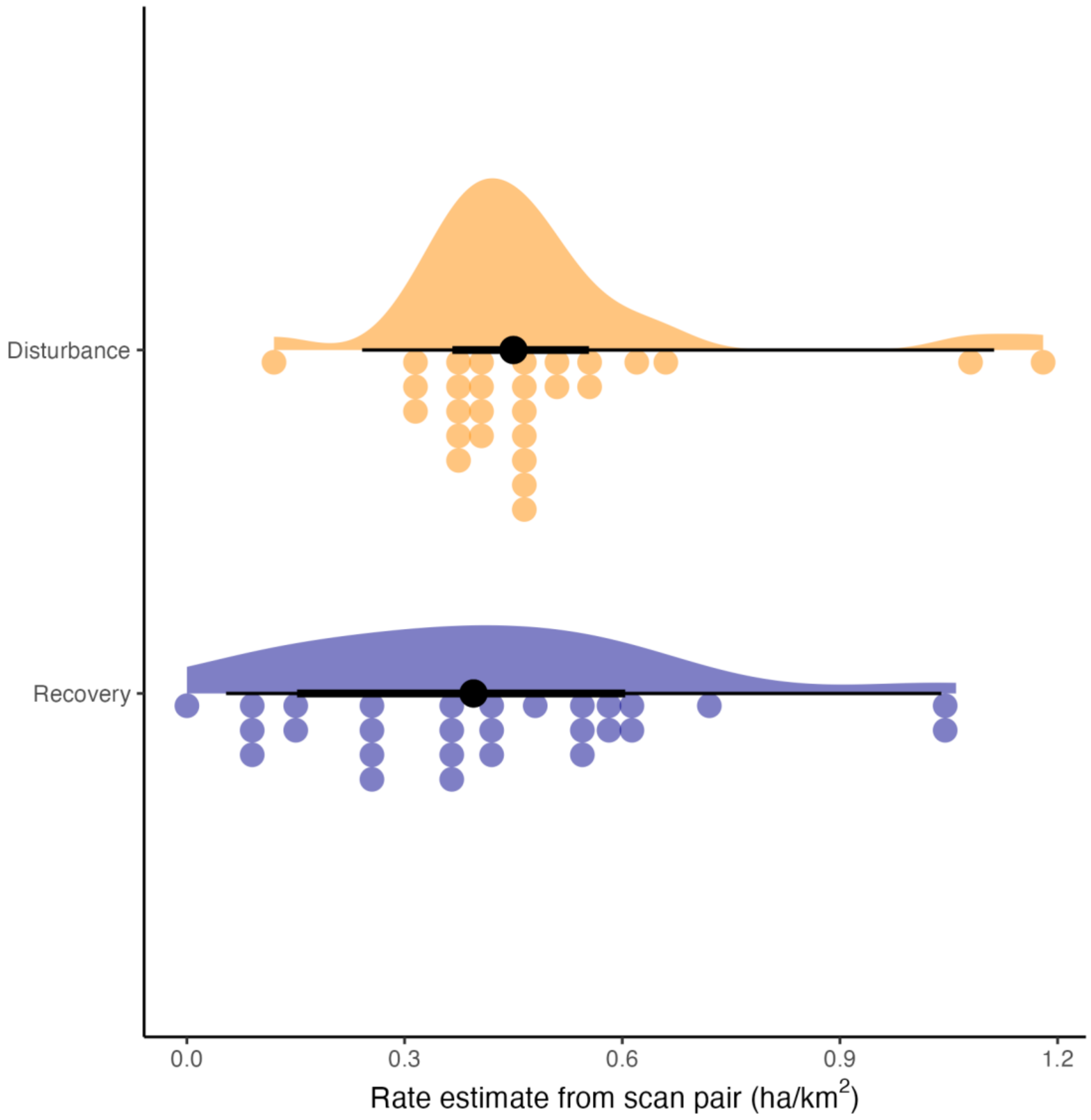

**Figure S3.4: Estimates of disturbance and recovery rates from scan pairs.** Shown are the distributions of estimates of *d* and *r* based on a pairwise inference procedure (Section S3.3) for all 28 scan pairs at Harvard Forest between 2012 and 2024. All values were corrected assuming a baseline measurement artefact of $d_0 = r_0 = 1$ ha km$^{-2}$ (or ~1% of canopy area, and 10% of gap area). The upper part of each plotting unit shows the density function, the lower part the binned values, and the black bars in between show the median, interquartile range and 95% range. Median values ($d = 0.45$ ha km$^{-2}$ and $r = 0.40$ ha km$^{-2}$) are near-identical to the values inferred from the models shown in Fig. 4 in the main text, with a lower uncertainty for disturbance rate estimates compared to recovery rate estimates. However, the ranges of inferred disturbance rates (95% interval: 0.24-1.11 ha km$^{-2}$) and recovery rates (0.05-1.04 ha km$^{-2}$) are extremely wide, reflecting both natural variation in disturbance rates and measurement uncertainty.

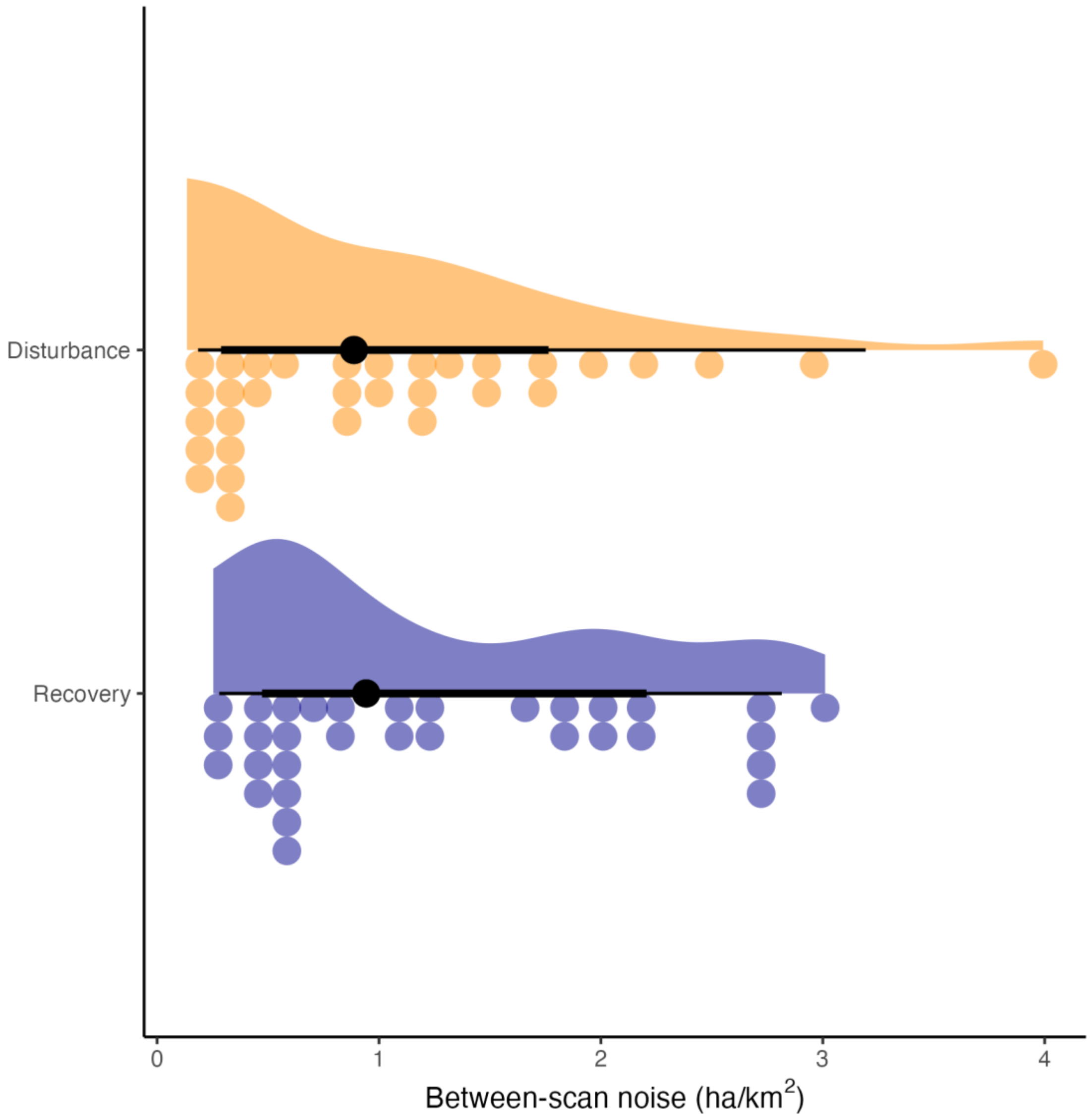

**Figure S3.5: Estimates of between-scan noise.** Shown are raw estimates of between-scan noise (disturbance and recovery artefacts $d_0$ and $r_0$) for 8 cells of 1 $km^2$ extent and 1 $m^2$ resolution at Harvard Forest. Values were obtained by comparing two sets of temporally coincident scans (NEON and GLiHT campaigns in the summers of 2012 and 2017), by treating each scan as first and last scan, respectively. This yielded a total of n = 32 comparisons. The upper part of each plotting unit shows the density function, the lower part the binned values, and the black bars in between show the median, interquartile range and 95% range. Note that median values lie close to 1 ha $km^{-2}$ and are thus in close agreement with the independently inferred intercepts $d_0$ and $r_0$ in Fig. 3 in the main text.
